# Temporal variability and its effects on diversity maintenance in an agroecological matrix

**DOI:** 10.64898/2026.07.10.737830

**Authors:** Verónica Zepeda, Luis Guillermo García-Jácome, Eugenio Azpeitia, Norma Leticia Abrica-Jacinto, Mariana Benítez

## Abstract

Agroecosystems are dynamic ecosystems, constituted by patches of vegetation and agricultural use, where biodiversity is shaped by spatial and temporal variability. While most studies have focused on spatial composition and configuration, the role of temporal variability remains poorly understood. Yet, temporal dynamics can strongly modify species composition, abundance, and persistence in ecological communities. Temporal variability is particularly relevant in agroecosystems with rainfed agriculture where environmental conditions shift dramatically between rainy and dry seasons. In this paper, we assess the role of temporal variability on biodiversity maintenance in an agricultural matrix using a metacommunity model that simulates an agricultural landscape under rainfed conditions, that is, with abrupt seasonal changes in the agricultural patches. This model couples a local community network dynamic with a migration dynamic and is based on empirically documented features of rainfed agricultural matrices. Our results show that temporal variability provides new opportunities for species to recover from low densities. However, the effect of temporal variability is not straightforward. It depends on the initial and final conditions, the migration and mortality rates and the intensity of temporal variability. Overall, our findings highlight the need to further investigate temporal variability to better understand its role in shaping biodiversity in agricultural landscapes.

## 1. Introduction

Agroecosystems –ecological systems modified for the production of food and other goods–are part of complex landscapes or agroecological matrices constituted by patches of vegetation and agricultural use (Perfecto et al. 2009). Agroecological matrices are often heterogeneous in space as a result of the different types of agricultural management practiced in a given region, and the arrangement and the size of vegetation and agricultural patches (Burel et al. 2005; Jeanneret et al. 2021). Moreover, these systems are also heterogeneous at different temporal scales (Burel et al. 2005), as a result of seasonal changes, interannual climate fluctuations and agricultural management practices. These temporal dynamics have important effects on the composition, abundance and persistence of species (Altieri 2004; Kremen et al. 2012a; Kremen et al. 2012b). In particular, temporal variability is more drastic in areas with marked rainy and dry seasons, where rainfed agriculture prevails, which can occupy up 80% of agricultural land (FAO 2011). Thus, to have a better understanding of the agroecosystems dynamics and their biodiversity maintenance we need to consider their temporal variability. However, such temporal dimension has often been neglected as much of the research has focused on spatial patterns of biodiversity within these systems.

Temporal variability characterizes all ecosystems and in many ecosystems this variability is expected to increase in the next century, affecting precipitation patterns and the frequency of storms and droughts (Adler et al. 2006). At population level, classic modelling efforts suggest that these time-dependent changes seem to be detrimental to the persistence of species due to increase in extinction risk (Kendall et al. 2002). On the other hand, at the community level, current evidence show positive effects for biodiversity maintenance as temporal variability might promote new opportunities for niche differentiation where species can coexist because species have different responses to the environment and because these responses might also modulate the intensity of biotic interactions (Adler et al. 2026; Chesson 2000). However, empirical evidence supporting a positive effect of temporal variability on biodiversity come from studies with only a few species, mostly with species of the same guild and without considering the surrounding landscape (Adler et al. 2026).

Agroecosystems are dynamic ecosystems in which biodiversity maintenance is determined by the establishment or movement of individuals through the landscape. They cannot be understood without considering their spatial composition and configuration, which determine the matrix permeability to biodiversity, this is, the so-called quality of the matrix (González González et al. 2016; Perfecto et al. 2010; Perfecto et al. 2009; Ramos et al. 2018). A high quality matrix allows metapopulations –a network of connected local populations– to be connected through the migration of individuals, which favors recolonization in places where the species had become locally extinct (Leibold et al. 2004; Levins 1969; Perfecto et al. 2009). Previous studies suggest that the type of vegetation or agriculture management (e.g. traditional or industrial) characterizing each patch also determine how permeable it is for the transit of wild species, thus facilitating or precluding species migration and recolonization (González González et al. 2016; Perfecto et al. 2010; Perfecto et al. 2009). Indeed, the ability of movement of the individuals through the landscape is vital to recolonize patches in which local extinction has occurred and avoid regional extinction, but also for the dynamics and functioning of the metacommunity (Loreau et al., 2003a; Loreau et al., 2003b).

Temporal variation in the quality of the agroecological matrix also affects its composition and configuration. In agroecosystems, temporal variability at larger scales (several years) can be result of crop rotation or reallotment affecting the size and the shape of cropping and vegetation areas; at short scales can be result of abrupt changes in land use, successive management operations or to periodic changes like crop phenology or seasonal variability (Bertrand et al. 2016; Burel et al. 2005; Gallé et al. 2018; Vasseur et al. 2013).

These factors might have an effect on resource availability due to variation in crop biomass and variation in resource availability (dos Santos et al. 2021; Ricketts et al. 2006). As a consequence, distribution patterns and abundance of species might be affected. For instance, in rainfed agriculture crops are planted during the rainy season of the year, they depend exclusively on rainwater as those systems are not irrigated and they drastically change from plots with high biomass in the rainy season to almost bare soil in the dry season. Thus, patches of rainfed agriculture may increase the quality of the agroecological matrix in the rainy conditions promoting diversity maintenance.

Another example of temporal variability in agroecological matrices is crop rotation. These have been shown to increase farmer’s gain and improve soil nutrients leading to a dynamic equilibrium, when used for a long time, as different species can use the different crops as complementary habitats in different seasons depending on management and agrochemical frequency applications (Ricketts et al. 2006). On the other side, abrupt changes in the cropping system can result in short-term high disturbance levels and loss or gain of land cover quality depending on species or functional group (dos Santos et al. 2021; Rebollar et al. 2017).

Although temporal variability has been considered to be a relevant driver of metapopulation and metacommunity dynamics in agroecological matrices, it remains largely unknown how temporal changes affect species abundance and diversity, and their consequences for species sustained coexistence in heterogeneous matrices (dos Santos et al. 2021). In previous studies (González González et al. 2016; Ramos et al. 2018) a spatially explicit metacommunity model was developed to assess the impact of the quality of an agroecological matrix on biodiversity. They found that biodiversity declines as the level of agricultural intensification increases and also is drastically affected by some configurations of heterogeneity of habitat patches. However, these studies were only focused on studying the role of spatial heterogeneity and do not explore the role of temporal changes in agroecological matrices.

In this paper, we develop a model to assess the role of different frequencies of temporal variation on the diversity of communities of wild species embedded in a mixed rainfed and irrigated agricultural matrix. In order to do this, we used a metacommunity model that simulates an agricultural landscape and couples a local community network dynamic with a migration dynamic.

## 2. Methods

### 2.1 Simulation of species networks

We first generated 300 graphs representing the structure of ecological communities, each corresponding to a network of 10 interacting species. The networks only included predatory interactions. Depending on their position in the network, species were classified as basal (species that are eaten by other species and do not eat other species), intermediate (species that are eaten and eat other species) and top species (only eat other species). To generate these networks, we used the niche model described by Williams and Martínez (2000), as previously done in the exploration of metacommunities in agroecological matrices that do not vary with time (González González et al. 2016; Ramos et al. 2018). The model consist of an algorithm that generates the structure of trophic networks from wo input parameters: the number of species (S) and the network mean connectivity (C). In this study, S = 10 and C = 0.2 were considered. We removed the networks with no basal species and with isolated species. After this procedure we kept 149 networks. From these networks, we built interaction matrices *α*, where every value *a_i,j_ = 1* whenever species *j* feeds on species *i*, otherwise *a_i,j_ = 0*. Then we use this information to set up a model of ordinary differential equations that describes the population dynamics of the interacting species. The equation describing the dynamic of each species *i* in the network had the form:

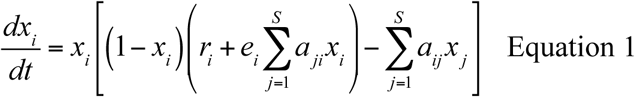

where *x_i_* is the abundance of species *i*, *r_i_* is the intrinsic growth rate of species *i*, *e_i_* is the assimilation (consumption) efficiency, S = 10 is the number of species included in the network/community. The equation’s initial values and parameters were obtained in the following way:

1. *x_i_* was randomly chosen from a uniform distribution, with values between 0 and 1.
2. *r_i_*was defined positive for basal species (primary producers) and negative for the rest as positive growth for basal species is expected in absence of its consumers and a decrease of the other species in absence of their prey. The magnitude of *r_i_* was randomly chosen from a uniform distribution, with values between 0 and 2.5 for basal species and −1 and 0 for the rest of the species. This is because previous works, with a different equation for the community dynamics, have shown these values to prevent non-computable explosive behaviors and several extinctions (González González et al. 2016).
3. Following Kéfi and collaboraors (2016) *e_i_* was defined to 0 for basal species, 0.66 for primary consumers and 0.85 for top consumers.
4. *a_i,j_* are the interaction relationships from the interaction matrix generated by the niche model.

### 2.2 Simulation of an agricultural matrix

To assess the effect of temporal variation over biodiversity maintenance of an agricultural landscape we simulated an agricultural system with two types of farming, rainfed agriculture and irrigated agriculture, and some patches of vegetation. This landscape was a lattice of 10 by 10 square patches, with alternated stripes of rainfed and irrigated agriculture. Then, 30% of the patches were assigned as patches of vegetation. While this is a completely hypothetical landscape, the identity of the patches and their proportion, this is, its composition, was inspired by an agricultural landscape previously characterized by our group in Oaxaca, Mexico (Fig. 1;(Urrutia et al. 2020). In such landscape, as in many agricultural regions in Mexico: 1) agriculture is the most abundant type of patch, 2) rainfed and irrigated agriculture coexist in an heterogenous mosaic and 3) rainfed plots usually exhibit a traditional and more biodiversity-friendly management than irrigated plot (Flores-Gutierrez et al. 2020; González González et al. 2020; León-Cortés et al. 2024), but change drastically from diversified agroecosystems like milpa to almost bare soil along the year (Rebollar et al. 2017)(Fig.1).

**Figure 1.**
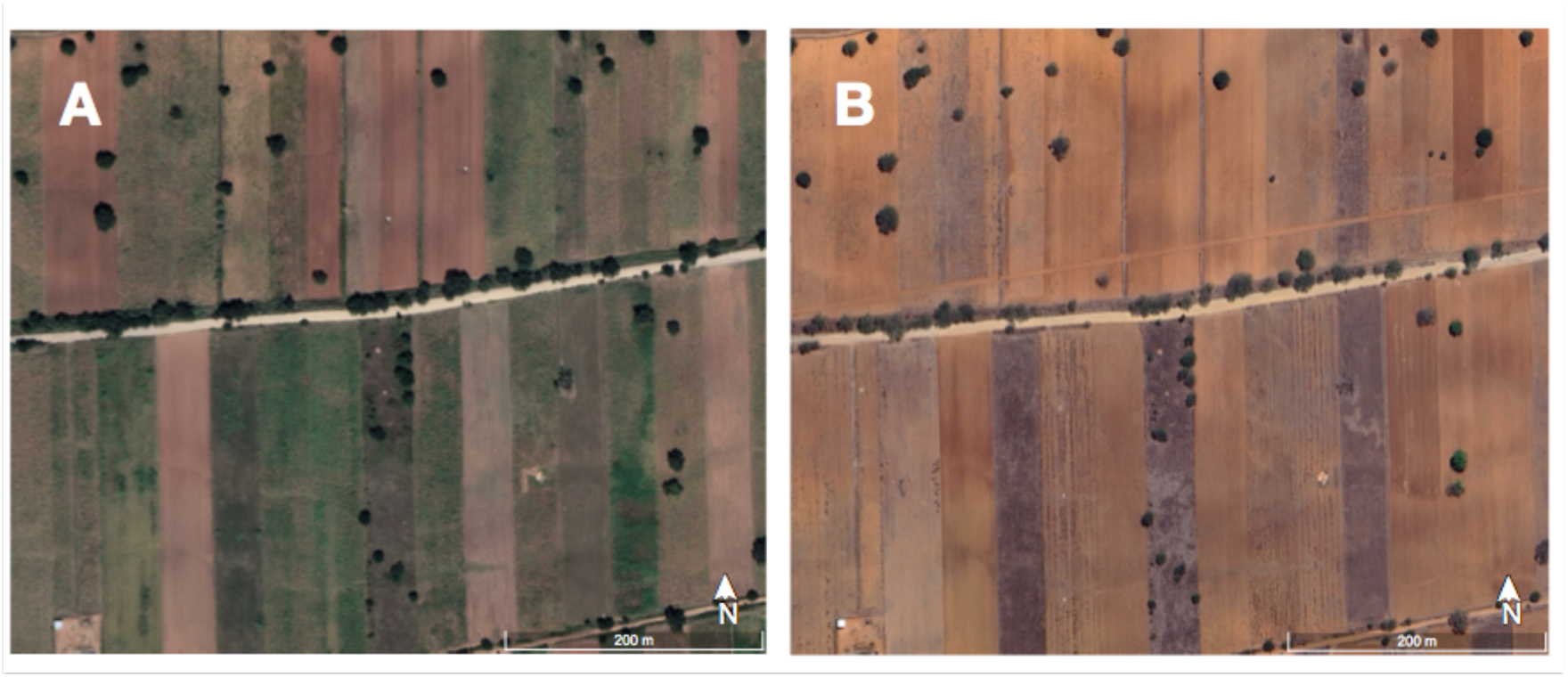
Plots of rainfed agriculture in Zaachila, Mexico, an agricultural area of central Valleys of Oaxaca, in rainy (A) and dry season (B). Images were taken from Google Maps.

Community dynamics only happens in the vegetation patches. As a consequence, species can only establish permanently and interact with other species in the vegetation patches as we assume that cells around habitat are not fit for reproduction and are merely to be migrated across (González González et al. 2016; Hanski et al. 2004).

### 2.3 Metacommunity model

We follow a metacommunity model described in previous works (González González et al. 2016; Ramos et al. 2018) that couples a local community network dynamic with a migration dynamic. This model takes as entries a community (characterized by its interaction matrix, dynamic equations and initial abundances, as described above) and the migration rates, which are assumed to be the same for all species in the network. The model iterates between community dynamics in continuous time and discrete steps of migration and mortality. Previous studies, and our own simulations, have found that, under the conditions described above, 100 units of time are sufficient to reach equilibrium under constant conditions (González González et al. 2016; Ramos et al. 2018), and that there were not oscillatory, explosive or chaotic dynamics in the model. During each unit of time, each local community interacts in habitat patches following the dynamic described by equation 1. Equation 1 was solved by Euler’s method using the ode function from the deSolve package in R (Soetaert et al. 2010). Thus, in each unit of time there were 1000 equally spaced subintervals, which corresponds to an integration step of *Δt* = 0.1. The choice of this value aims to ensure a balance between numerical accuracy and computational efficiency of the method. After community dynamics were run, a proportion of each population migrates to the eight direct neighboring patches. This process was repeated five times. Once those migration steps have been completed, a proportion of the individuals that arrive at each of the patches dies. The proportion of migration and mortality depends on the identity of the patches reflecting the fact that the quality of the patch has been shown to be affected by the type of agriculture, land management and spatial configuration (Fig 2) (Papaïx et al. 2015; Perfecto et al. 2009). The output of the model is the spatial distribution of species across the landscape in steady state, through which we can assess the changes in the diversity and the abundance of simulated wild species.

**Figure 2.**
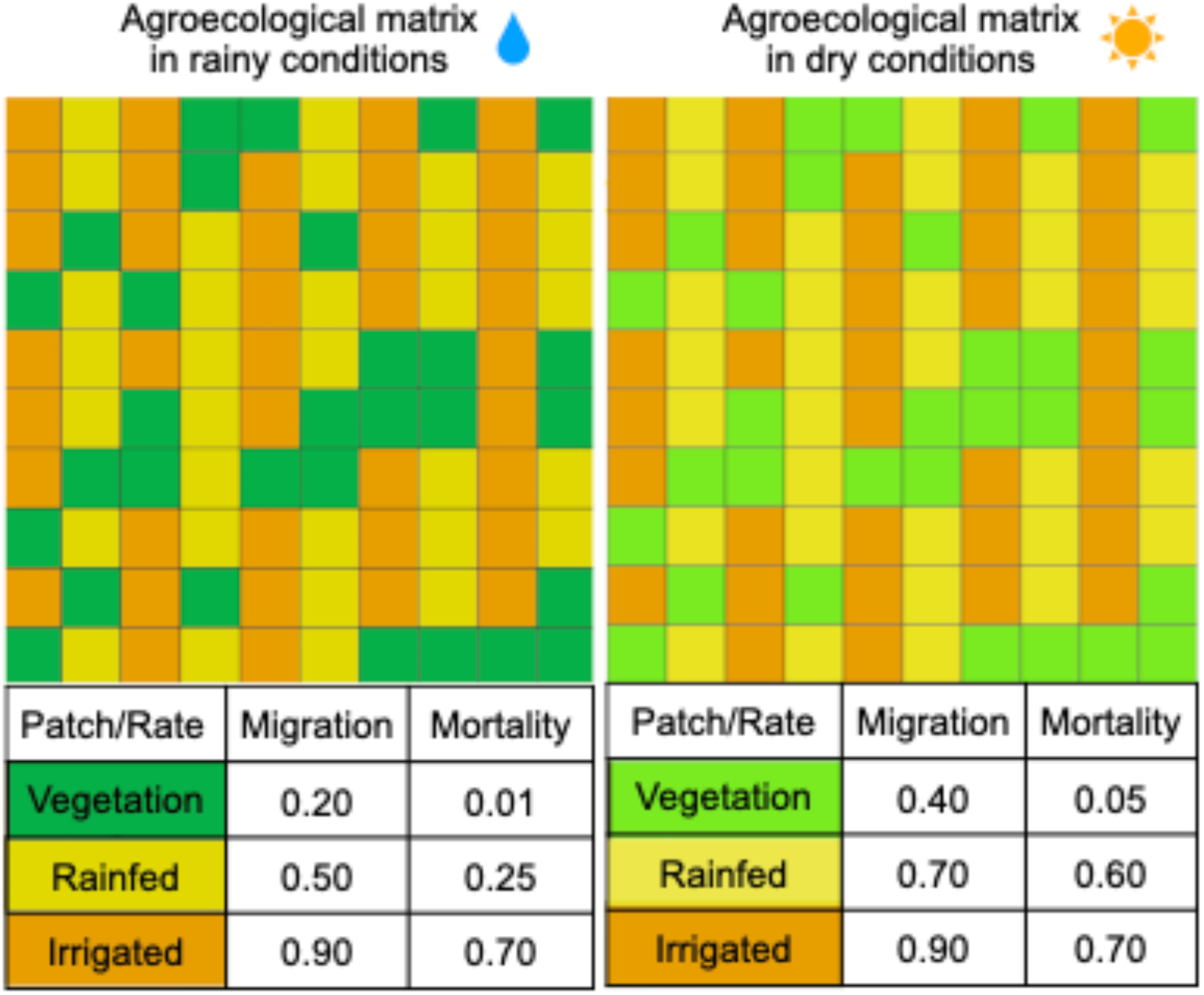
Temporal variability in the simulated agricultural matrix was simulated by changing the migration and mortality rates for each type of patch: vegetation, rainfed agriculture and irrigated agriculture.

We run several sets of simulations to couple the trophic interactions dynamics and the migratory steps to define the number of migratory and dead steps. To this, each of the 149 simulated communities was systematically tested for coupling. This was necessary so the spatiotemporal dynamics were not driven only by species interactions nor migration and dead steps, but by the joint action of these two processes (Appendix A).

### 2.4 Simulated experiments

To explore the effects of temporal variation on the richness and abundance of wild species we simulated experiments with different levels of temporal variation by changing the value of the migration and mortality parameters.

The quality of the matrix might be different in different seasons (Rebollar et al. 2017; Tavares Dias et al. 2018) (Fig. 1), with lower migration and mortality rates in the rainy conditions than in dry conditions (Fig. 2). To establish a baseline for comparison with temporally varying dynamics, we first simulated experiments without temporal variability by iterating the metacommunity model 100 times with the rainy condition and another experiment by iterating the metacommunity model 100 times with the dry condition.

To assess the role of different temporal periods, we alternate the rainy and dry parameters by partitioning the 100 iterations of the metacommunity model to simulate experiments with low, moderate and high temporal variability. For experiments with low variability, we iterated the model 50 iterations by starting with a rainy state in the sequence of migration and mortality parameters. After this step we used the spatial distribution of the species across the landscape to run another 50 iterations but now with the dry condition parameters (Fig. 3 low variability A). For experiments with moderate variability we partitioned the 100 iterations in five equal parts. Thus, we started with 20 iterations of the metacommunity model and the rainy condition parameters. Then we repeated the same procedure described before until we completed the 100 iterations (Fig. 3 moderate variability A). Finally, for experiments with high variability, we partitioned the 100 iteration of the metacommunity model into ten equal parts. Thus, we iterated the model 10 times with the parameters of rainy conditions and then another 10 iterations with the dry conditions parameters and so on until complete the 100 iterations (Fig. 3 high variability A). Then, we repeated all the sets of experiments but starting with the dry condition parameters as it can be seen in figure 3 B.

**Figure 3.**
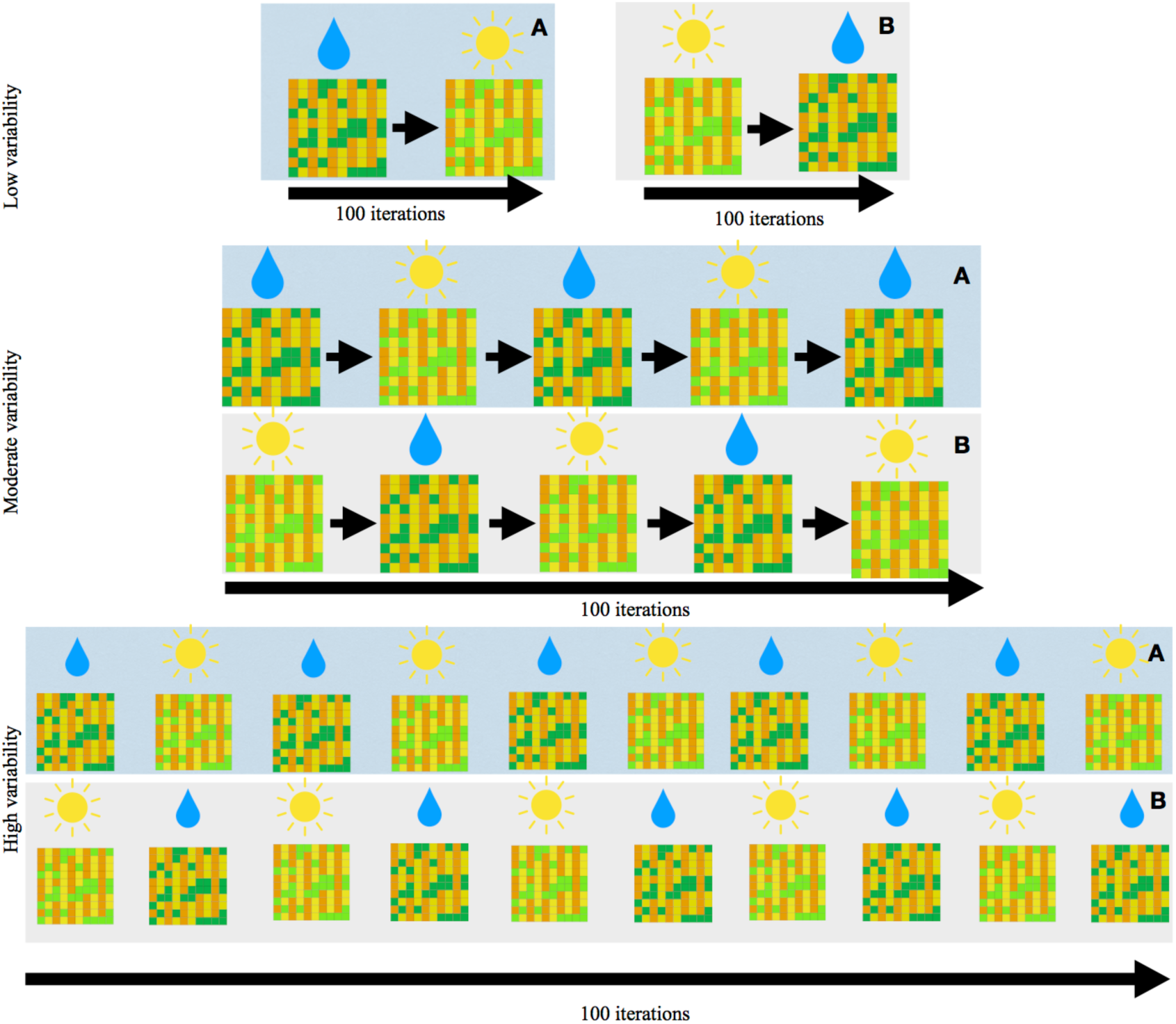
Set of experiments with temporal variability. Panel A represents experiments starting under rainy conditions, whereas Panel B represents experiments starting under dry conditions.

### 2.5 Diversity measures and analysis

After the 100 iterations, for each network in each of the experiments, we recorded the abundance of each of the species in all the patches. Species richness was measured as the number of species with abundance above 10^-6^ in the final time step. We also compute Hill numbers, in which the importance of the relative abundance of each of the species in the community is regulated by the parameter *q*. Thus, in ^1^*D*, *S* is weighted by the proportions of the species abundances and can be interpreted as the effective number of species equally abundant within an assemblage, which is equivalent to the exponent of Shannońs entropy (Jost 2006). In ^2^*D*, the diversity values favor the most abundant species, and the less abundant or rare species are almost not accounted for; consequently, ^2^*D* can be roughly interpreted as the effective number of dominant or the most abundant species in the community (Jost 2006). To compute Hill numbers we use the hill_taxa function from the hillR package in R (Li 2018). For each experiment, we computed the mean total abundance, the mean species richness, the mean ^1^*D* and ^2^*D* values across the 149 communities, along with their 95% confidence intervals for each diversity measure and for species abundances.

## 3. Results

Our results show that both the highest and lowest species abundance and diversity occurred in experiments with no temporal variability (i.e., under constant conditions).

Species abundance and diversity were consistently higher in experiments that ended up in rainy conditions, regardless of their initial setup (Fig. 4). This pattern weakened as temporal variability increased. In contrast, all experiments that ended up under dry conditions showed similar mean values of species abundance and diversity, regardless of the initial parameter conditions or the magnitude of temporal variation (Fig. 4). When comparing species richness between scenarios with no variability and those with low, moderate, and high variability, we observed a slight increase in diversity (Figs. 4 and 5).

**Figure 4.**
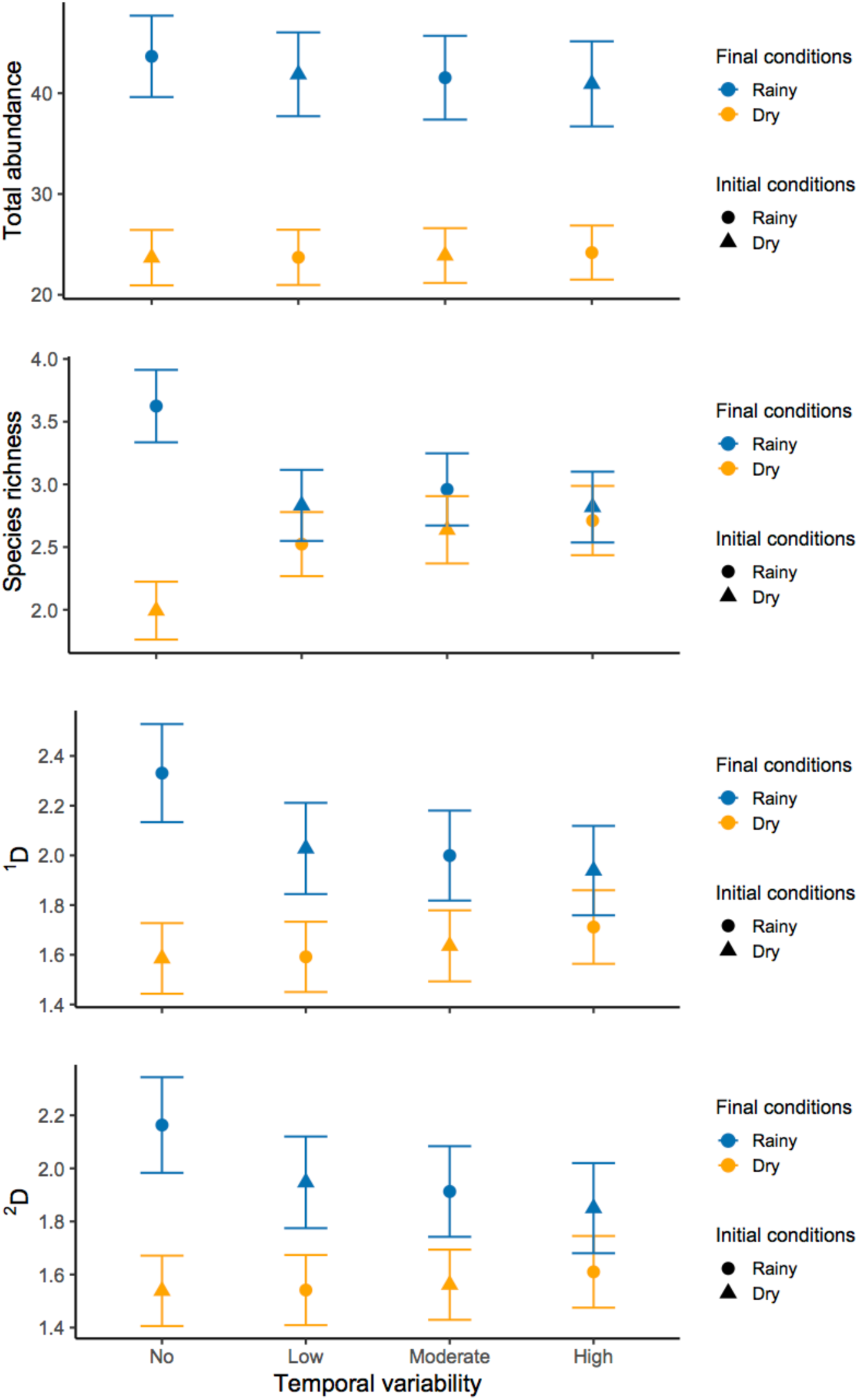
Results of total abundance and species diversity at the end of each experiment. Each point represents the mean of the 149 simulated communities and the error bar is the 95% IC for each of the treatments. Blue lines indicate the experiments that ended up in rainy conditions and orange ones the experiments that ended up in dry conditions.

**Figure 5.**
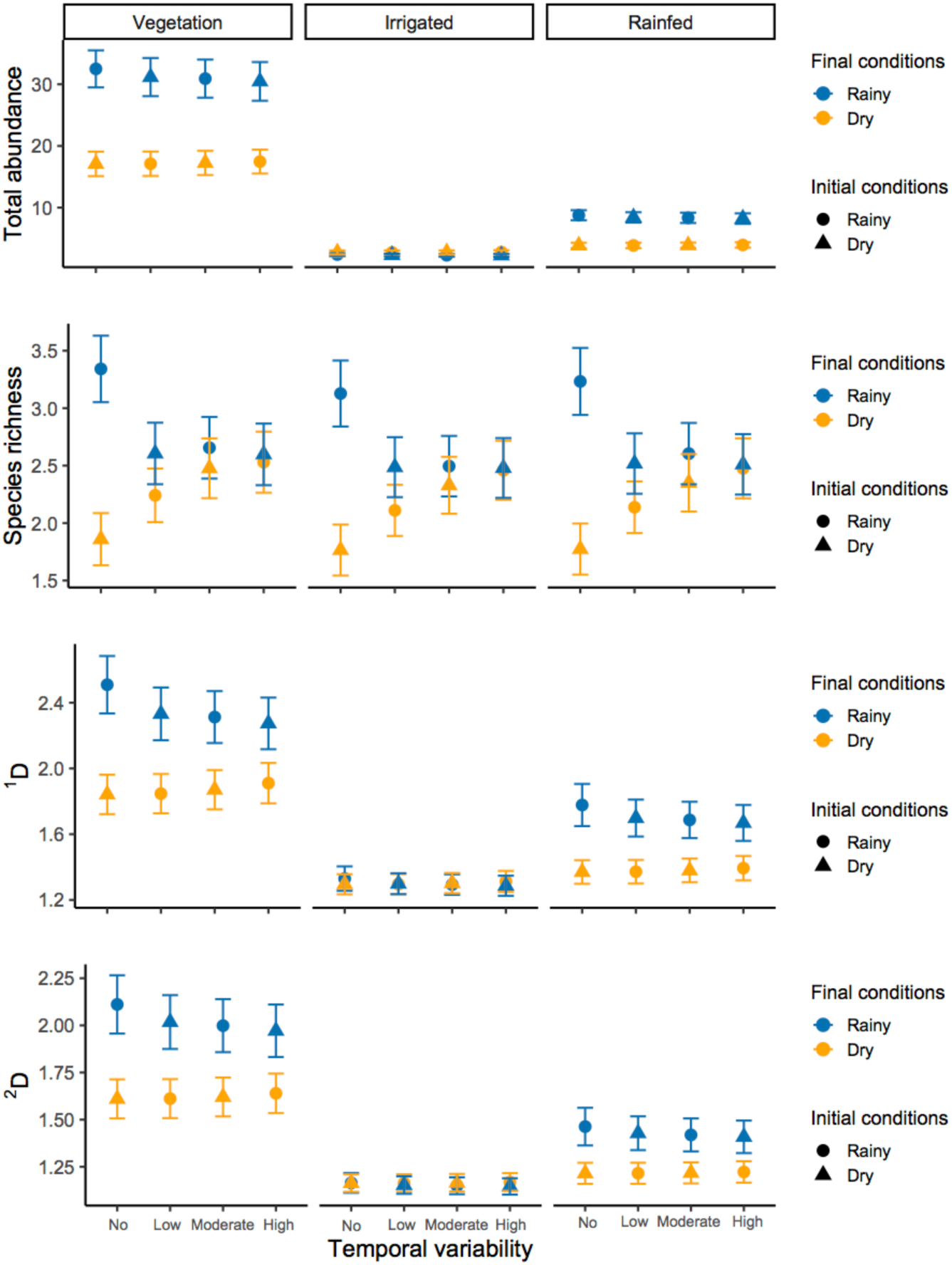
Results of total abundance and species diversity at the end of each experiment by patch type. Each point represents the mean of the 149 simulated communities and the error bar is the 95% IC for each of the treatments. Blue lines indicate the experiments that ended up in rainy conditions and orange ones the experiments that ended up in dry conditions.

We also computed the mean species abundance and diversity of each network over the 100 iterations for all experiments. Overall, abundance and species diversity tended to be higher in experiments that began under rainy conditions. In contrast, all metrics declined as temporal variability increased. However, in most cases, the confidence intervals overlapped (Appendix B).

We found the same trend described before in species abundances and species richness when we take into account the type of patch. However, the mean of ^1^*D* and ^2^*D* was always higher in habitat patches as a result of greater abundance, followed by patches of rainfed agriculture (Fig. 5). Along the same lines, ^1^*D* and ^2^*D* collapsed in patches of irrigated agriculture as the abundances of the species were too low in these patches (Fig. 5).

The mean values of species abundances, ^1^*D* and ^2^*D* were consistently higher in habitat patches, followed by rainfed agriculture (Fig. 5). In contrast, ^1^*D* and ^2^*D* declined sharply in irrigated agriculture, where species abundances were too low to sustain higher diversity values (Fig. 5).

Finally, when analyzing community dynamics across the 100 iterations, we observed that in experiments starting under dry conditions with low variability, some populations recovered from near extinction, leading to a recovery of species diversity within the community. This pattern was observed in 70 out of the 149 communities (see example in Fig. 6 or Appendix C for full disclosure of the community dynamics).

**Figure 6.**
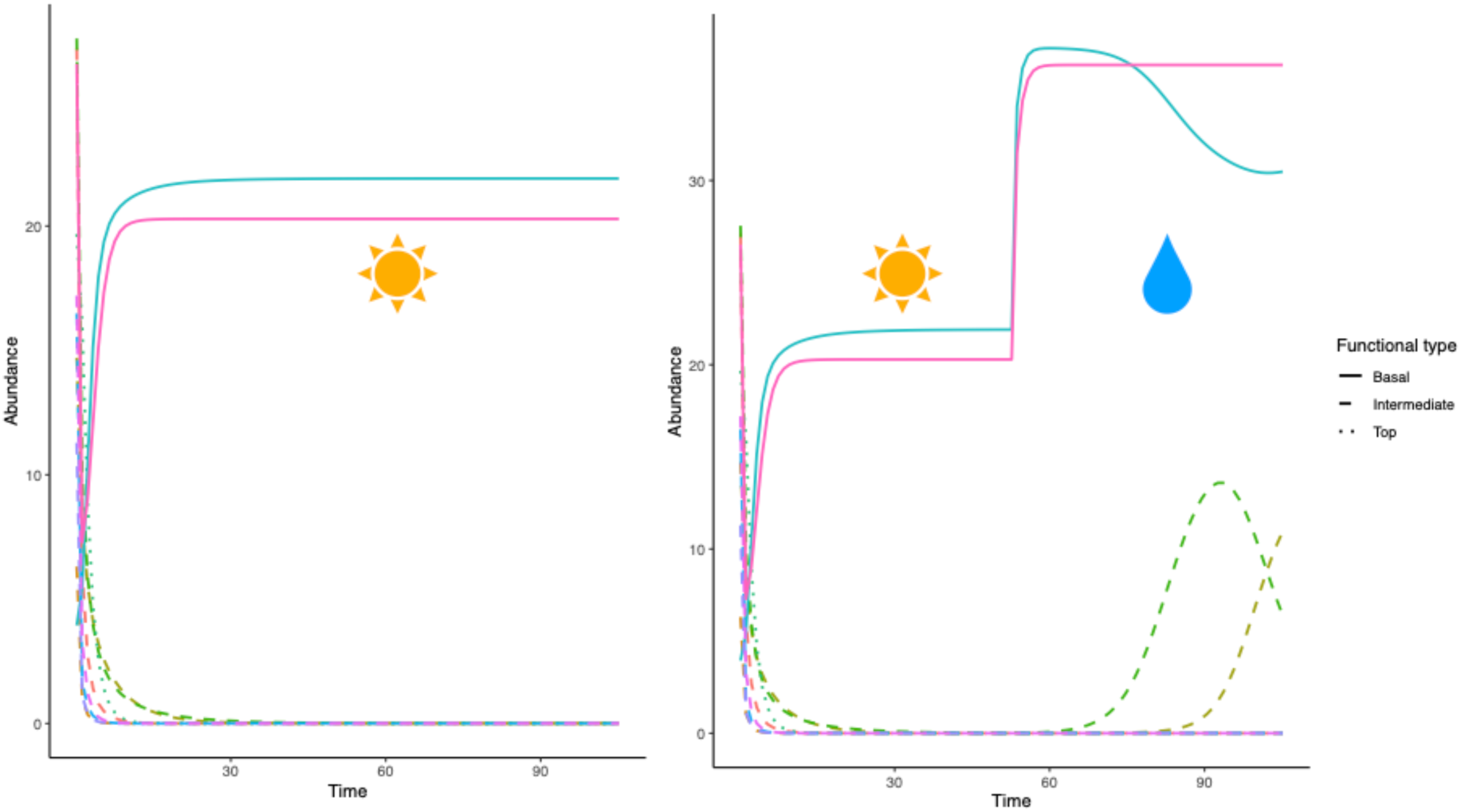
Example of a community dynamics in the experiments with no variability (dry condition) and the experiment with low variability, starting in dry conditions followed by rainy conditions.

Diamonds indicate the experiments that started in rainy conditions and circles the experiments that started in dry conditions.

Diamonds indicate the experiments that started in rainy conditions and circles the experiments that started in dry conditions.

## 4. Discussion

This study explored the impacts of seasonal changes in the quality of a simulated agricultural matrix on the abundance of individuals and species diversity in metacommunities. Our results showed that the effects of the temporal variability on diversity maintenance are context-dependent as its influence is related to the initial and ending conditions of the systems as well as the interval of time in which rainy conditions change to dry conditions and vice versa.

Constant conditions have an important influence on species abundances and diversity. The experiments with constant rainy conditions had the highest mean species abundance and the highest diversity values. In contrast, the experiments with dry constant conditions had the lowest species abundances and species diversity. These results are in line with the assumption that species diversity is promoted when the quality of the matrix is improved by increasing the permeability of a number of patches (Perfecto et al. 2010; Perfecto et al. 2009).

Temporal variability plays a context-dependent role in shaping biodiversity outcomes, with its effects varying according to baseline environmental conditions. All experiments that included temporal variability showed lower biodiversity and species abundances compared to the treatment under constant rainy conditions. However, when temporal variability simulations are compared with the constant dry treatment, positive effects on biodiversity emerge, as both species abundance and diversity increase under temporally variable conditions. This pattern was even more marked in the experiments with high temporal variability than in those conducted under constant environmental conditions. Moreover, species abundances and diversity were consistently higher in experiments that ended up under rainy conditions than in those that ended under dry conditions, regardless of the initial conditions. Under our modeling framework, temporal changes in community parameters were implemented as variation in matrix quality and in the duration of rainy and dry conditions experienced by each network. Thus, these results are also consistent with the higher matrix quality under rainy conditions, where species exhibit greater survival and establishment success. They also indicate that the model captures expected dynamics in seasonal environments (Zeppel et al. 2014), as is the case of rainfed agroecological matrices, with most species thriving under rainy conditions and declining during drought.

Water availability was introduced in our model as “rainy conditions” and was overall associated with higher abundance and species diversity. However, in real agroecological matrices, water availability depends not only on precipitation, but also on agroecological practices that help maintain soil moisture (Wezel et al. 2025). Some of these practices are cover cropping (Moore 2023), agroforesty and tree planting (Acharya et al. 2018), crop rotation and diversification (Wezel et al. 2025) or windbreaks and shelterbelts (Enescu et al. 2025). By incorporating these agroecological practices, farmers can effectively enhance moisture retention, improve soil health and, more generally, improve the quality of agroecological matrices.

Temporal variability may provide opportunities for species to recover from low population densities. The fact that some populations were able to recover when rare under low variability and dry initial conditions scenarios suggests that the interval between rainy and dry periods may be critical for biodiversity maintenance, highlighting the importance of precipitation events following droughts in rescuing vulnerable populations. This pattern also suggests that there is a lag in species’ responses to favorable conditions, as such dynamics were not observed in scenarios with moderate and high variability. Probably, that was because the period of time that communities experienced the transition between dry and wet conditions was too small to give a change to species to recover from low densities. Additionally, a qualitative analysis of these dynamics suggests that species recovering from low abundances were able to do so because basal species maintained sufficiently high abundances to support intermediate and/or top species within the community (See Appendix C). This highlights the importance of basal species for the persistence of higher trophic levels and important key drivers for community resilience. Moreover, these results also suggest that temporal variability does not impact all trophic levels in the same way and suggest asynchronous responses for the different trophic levels in which lower trophic levels enable subsequent recovery of intermediate and top species.

Landscape heterogeneity was an important driver for diversity maintenance in the agroecological matrix. Across patch types, clear differences emerged in how communities responded to environmental conditions and temporal variability. Vegetation and rainfed patches exhibited strongest temporal dynamics, highlighting their dependence on precipitation. In contrast, irrigated patches only showed changes in species richness to shifts between rainy and dry conditions. Species abundances were always too low as mortality and migration parameters do not allow them to increase.

## 5. Conclusions

Our results provide some insight about how temporal variability in agricultural landscapes might impact biodiversity as evidence in this matter is still scarce. The model presented in this work provides new opportunities to explore the role of temporal variability in matrices with different compositions, configurations and types of land management. Further work using this type of model could help explore how different matrix compositions (number and relative area of the patches) and configurations (spatial arrangement of the patches) interact with variability, among other relevant processes.

Variability has heterogeneous effects on metacommunity persistence and diversity in agroecological matrices. Yet, as it often happens in complex ecological systems, their effects depend on the context, in this case the effects of temporal variability are the result of the initial and final conditions, the value of the parameters associated to the quality of the matrix (migration and mortality), and the frequency of variability. Our model provides a theoretical basis to continue exploring the role of variability in agroecological systems.

## Supporting information

Supplemental figure migration steps

Supplemental figure time average

Supplemental community dynamics graphs

## Acknowledgments

VZ thanks to the Postdoctoral Grant provided by the DGAPA and the support of the Instituto de Ecología, UNAM. LA also thanks to the Centro de Ciencias Matemáticas, UNAM and to the UNAM postdoctoral program, DGAPA. MB was supported by UNAM PASPA – DGAPA.

## Author contributions: CRediT

**Verónica Zepeda:** Conceptualization, Formal analysis, Investigation, Methodology, Project administration, Software, Validation, Visualization, Writting – orignal draft, Writing – review & editing. **Luis Guillermo García-Jácome:** Methodology; Software, Validation, Writing – review & editing. **Norma Leticia Abrica-Jacinto:** Methodology; Software, Validation, Writing – review & editing. **Eugenio Azpeitia:** Methodology; Writing – review & editing. **Mariana Benítez:** Conceptualizacion; Supervision, Methodology; Writing – review & editing.

