## Supplemental figure migration steps for "Temporal variability and its effects on diversity maintenance in an agroecological matrix"

**Appendix A**

Once the 149 communities to be analyzed were obtained, the next step was to evaluate the appropriate coupling between trophic dynamics and the number of migration and mortality steps. Using too few migration steps makes trophic dynamics effectively too fast, causing spatial movement of individuals to have a negligible effect on species persistence. Conversely, using too many migration steps leads migration dynamics to dominate, altering population sizes without allowing trophic interactions to meaningfully influence the system.

To address this, we systematically explored different durations of trophic dynamics combined with varying numbers of migration and mortality steps under both constant rainy and dry conditions. Initially, we considered one mortality step per migration step; however, this configuration caused species abundances to decline too rapidly. We therefore adopted a modified approach in which a single mortality step is applied after a sequence of migration steps, preventing unrealistically fast declines in population sizes.

Regarding trophic dynamics, we first ran the metacommunity model (trophic dynamics + migration steps + one mortality step) using five time units for trophic dynamics, progressively increasing this duration in increments of five up to 50 time units. However, species abundances again declined too rapidly, and spatial dynamics had little effect on species persistence. Consequently, we reduced the trophic dynamics duration to one time unit and increased it stepwise up to five time units. Finally, we coupled the number of migration steps to trophic dynamics, starting with one migration step and increasing incrementally up to seven.

We found that one unit of time of the trophic dynamic and at least four migration steps were enough to see differences in species richness between rainy and dry conditions (Fig. 1).

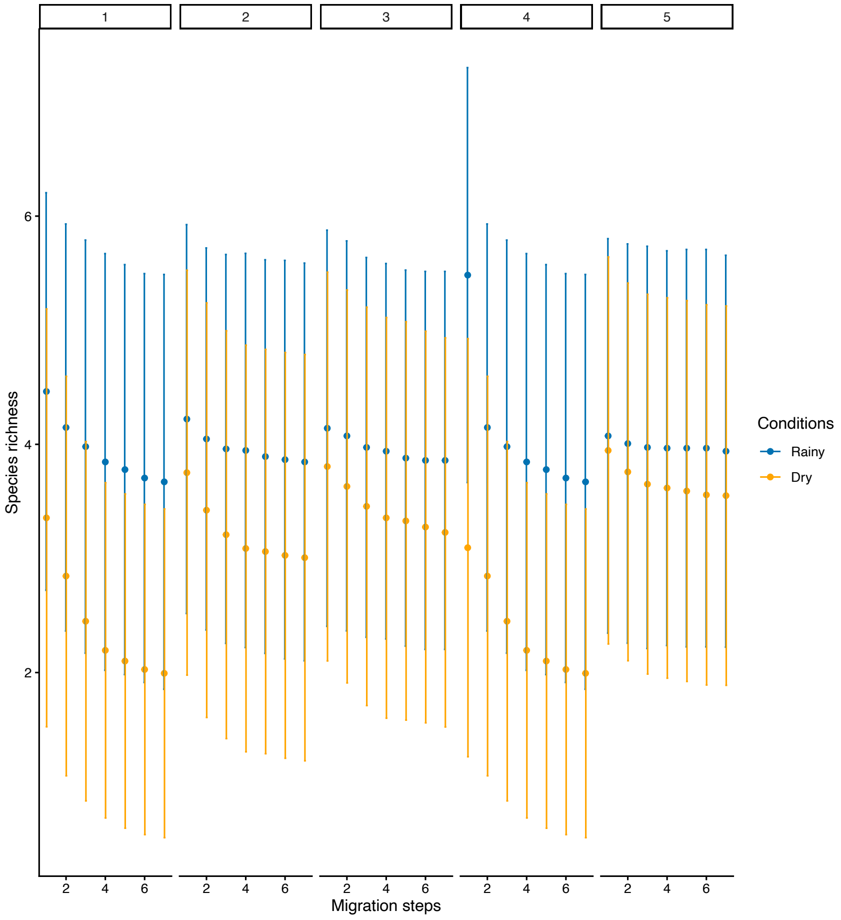


**Figure A.1** Species richness at the end of each combination of trophic dynamics duration, migration steps, and a single mortality step. Each point represents the mean across the 149 simulated communities, and error bars indicate the standard deviation. Blue lines correspond to simulations run under rainy-condition migration and mortality parameters, whereas orange lines represent dry-condition parameters. The top panels indicate the duration (in units of time) over which trophic dynamics were simulated.
