## Supplemental figure time average for "Temporal variability and its effects on diversity maintenance in an agroecological matrix"

### Appendix B

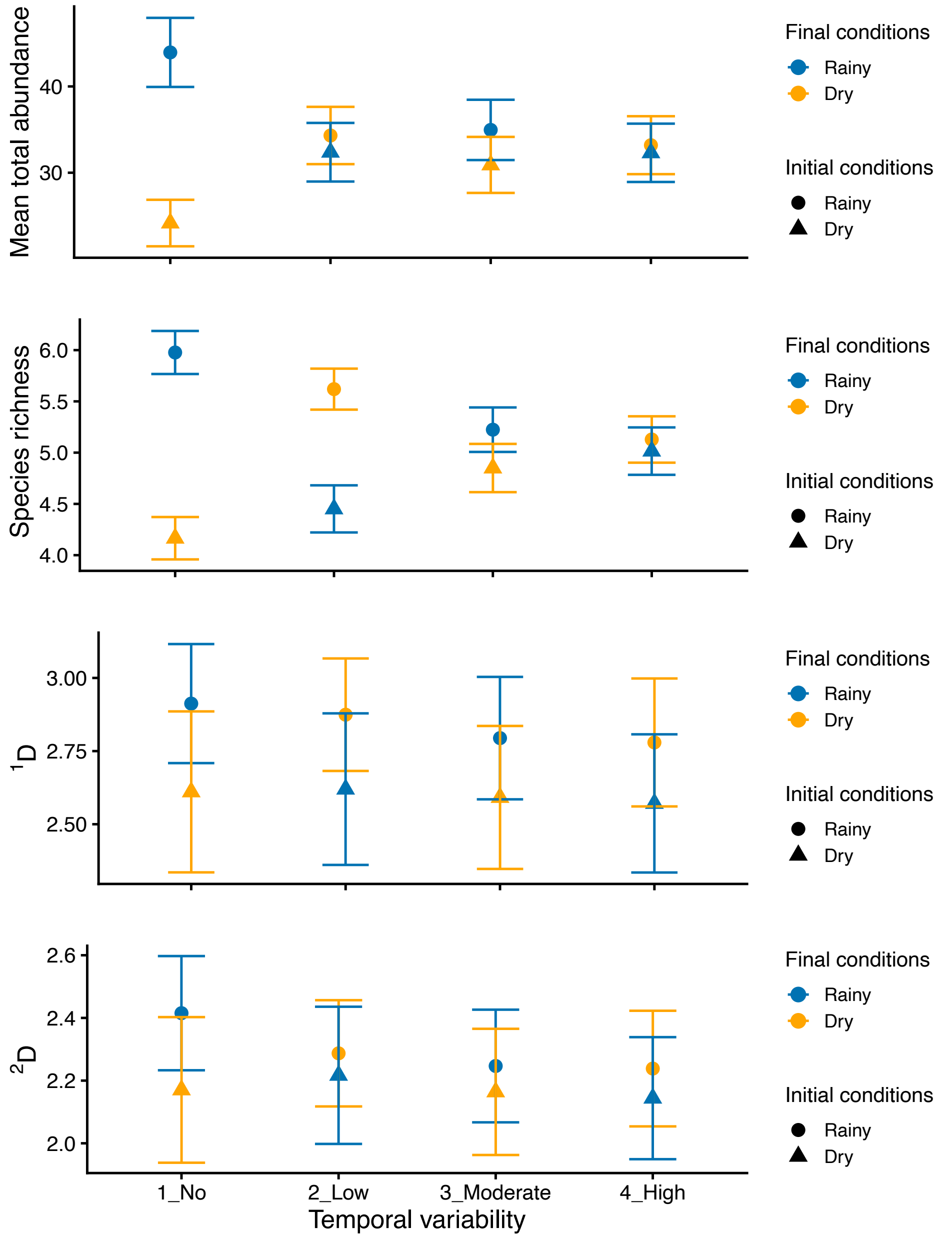


**Figure B.1** Total abundance and species diversity. Each point represents the mean value across the 149 simulated communities, averaged over 100 iterations. Error bars indicate the 95% confidence interval for each treatment. Blue lines indicate the experiments that ended up under rainy conditions, whereas orange lines represent those that ended up under dry conditions.

Diamonds indicate experiments that started under rainy conditions, and circles indicate those that started under dry conditions.
