## Supplemental community dynamics graphs for "Temporal variability and its effects on diversity maintenance in an agroecological matrix"

**Appendix C:** Community dynamics of experiments with low temporal variability, starting under dry conditions. Here we only show networks in which species recover when rare under rainy conditions.

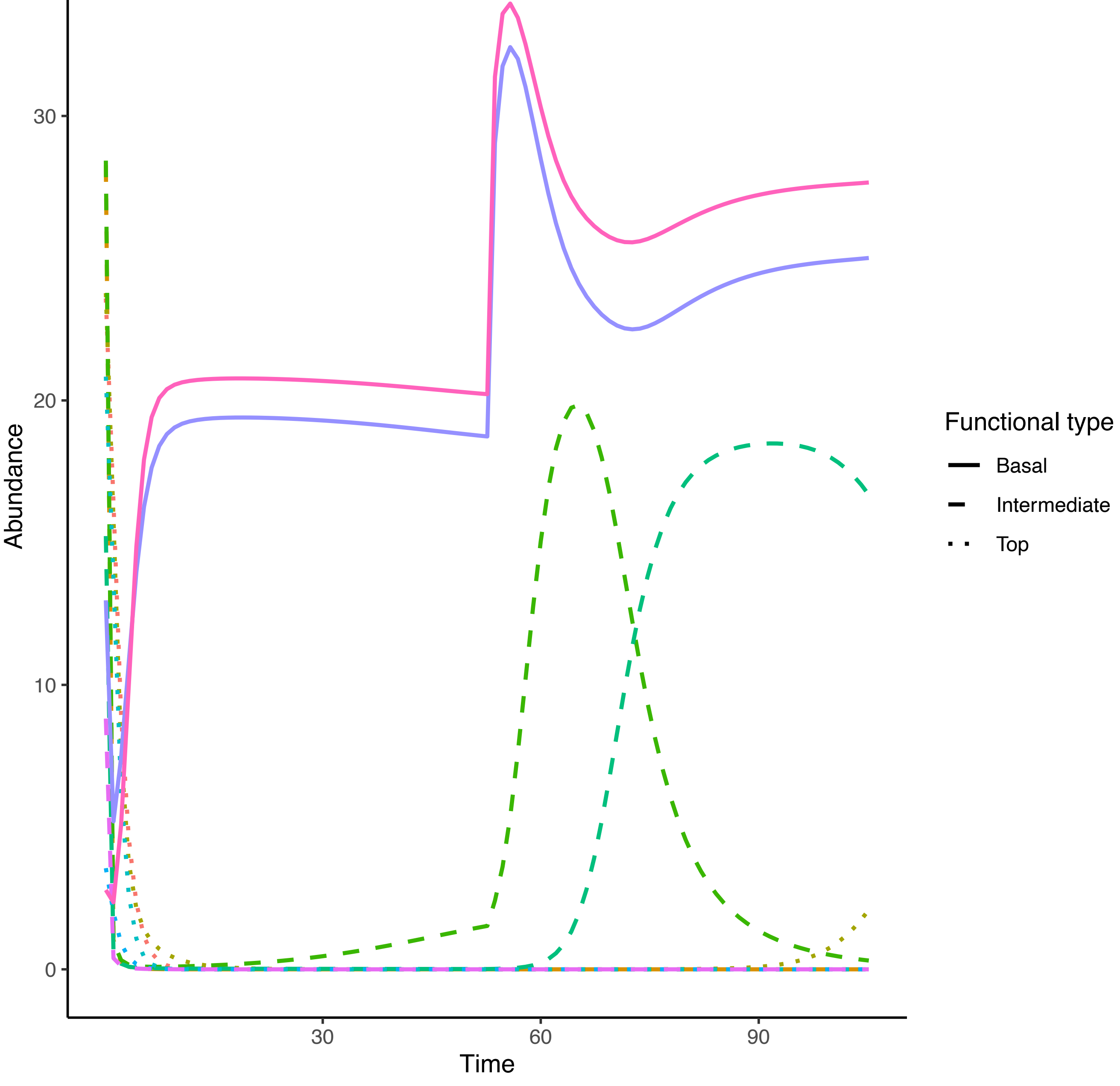

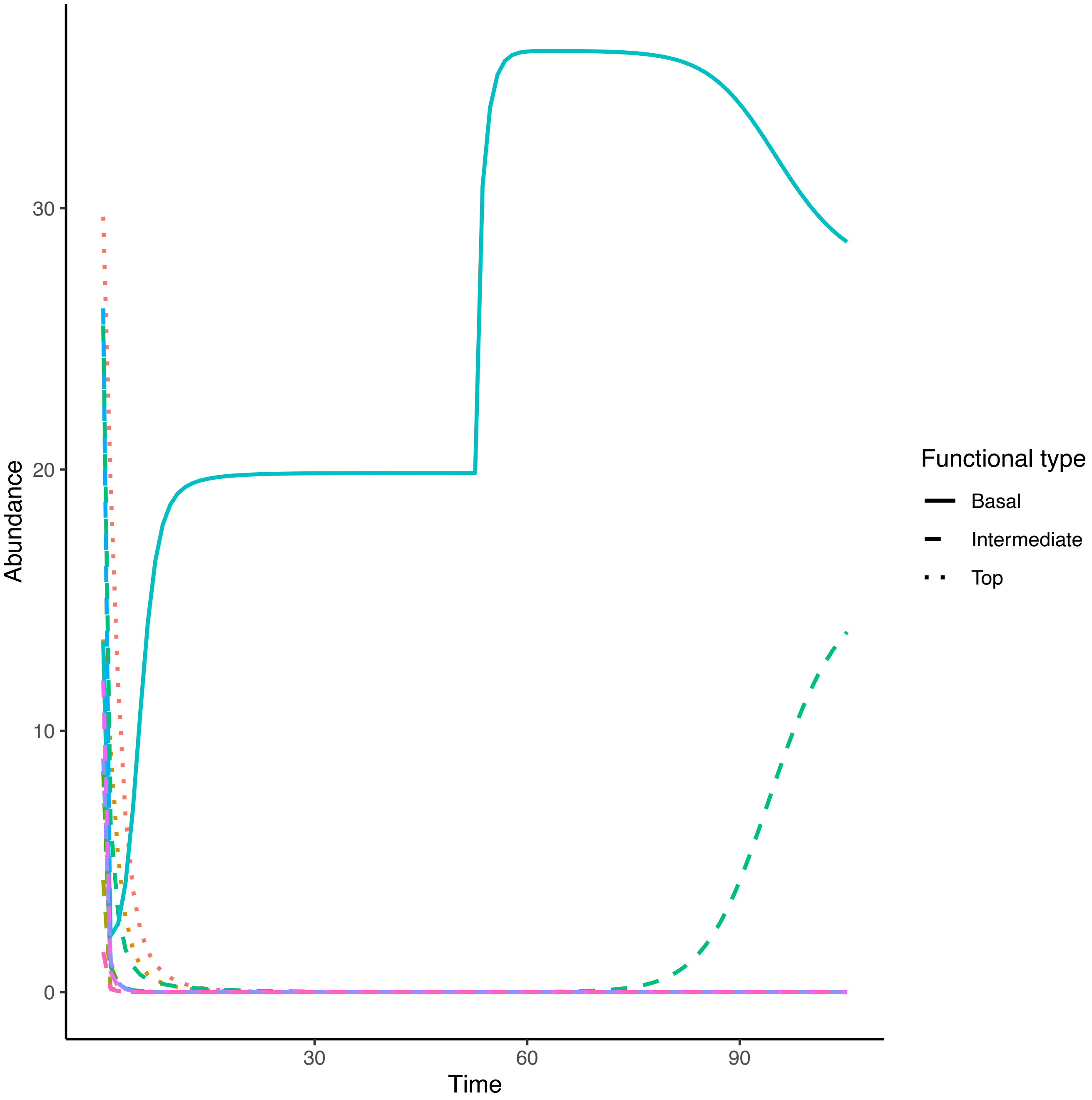
Abundance
30
20
10
0
30
60
90
Time
Functional type
Basal
Intermediate
Top

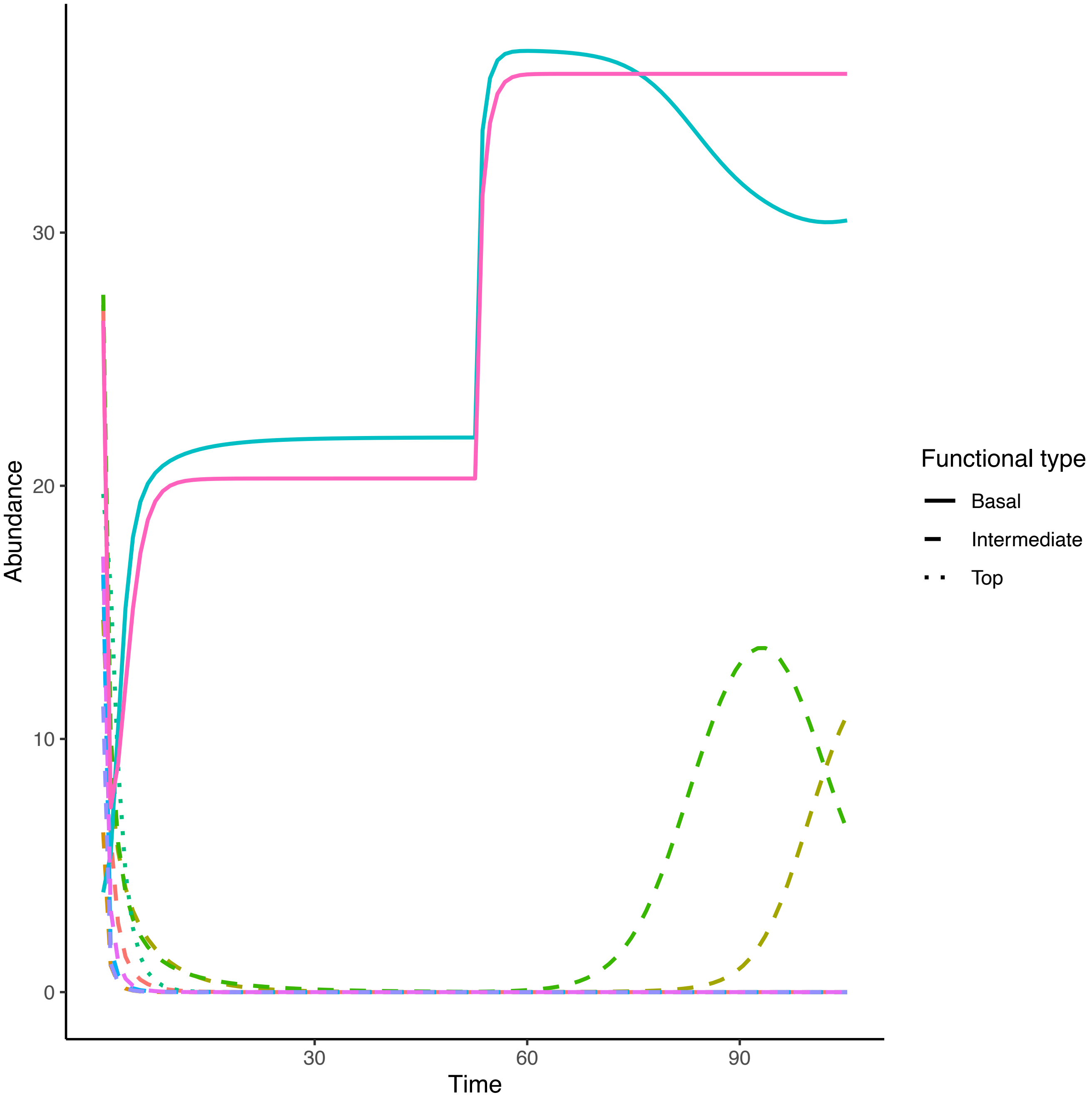

Abundance
0
10
20
30
Time
30
60
90
Functional type
Basal
Intermediate
Top

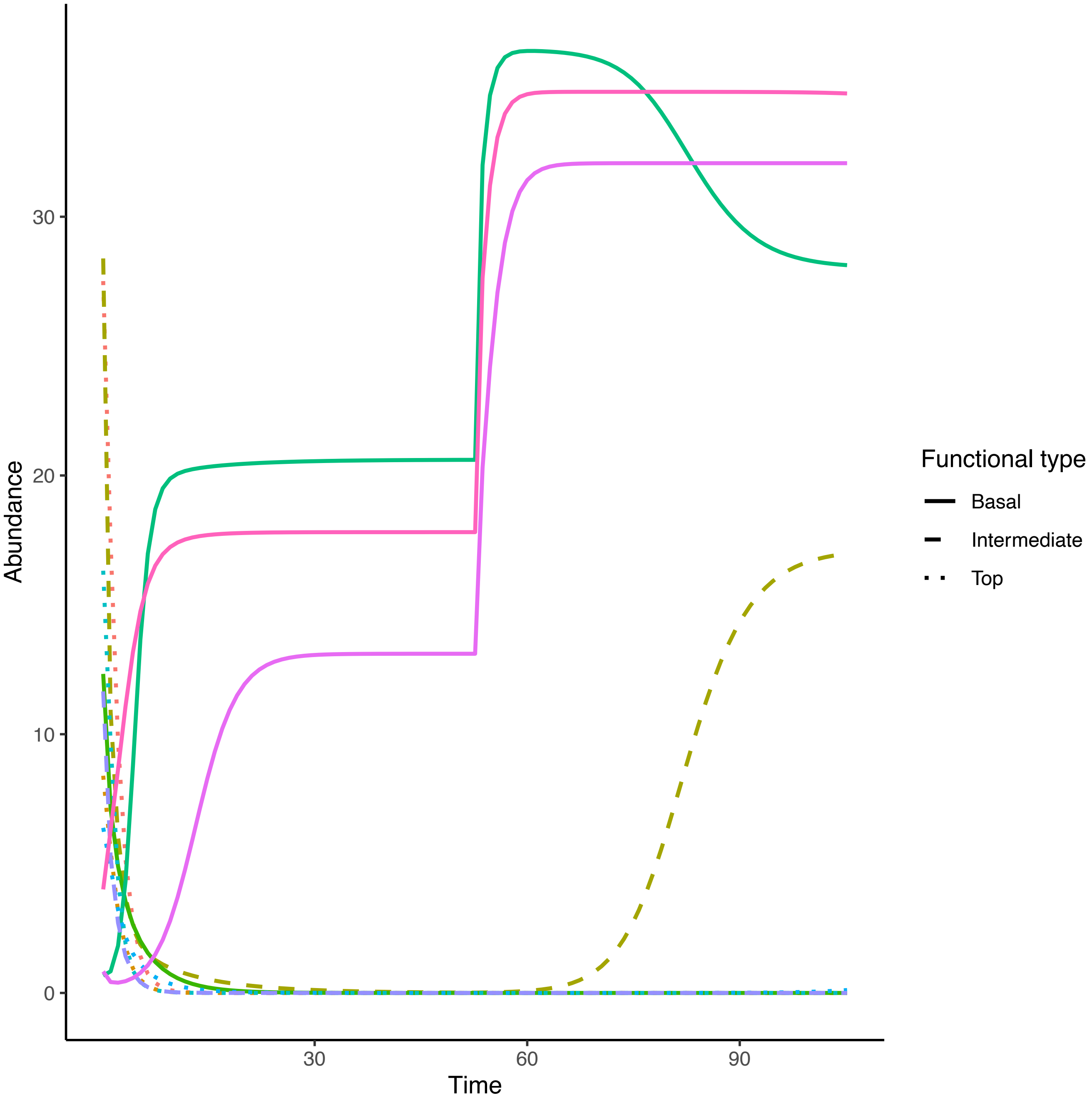

Abundance
0
10
20
30
Time
30
60
90
Functional type
Basal
Intermediate
Top

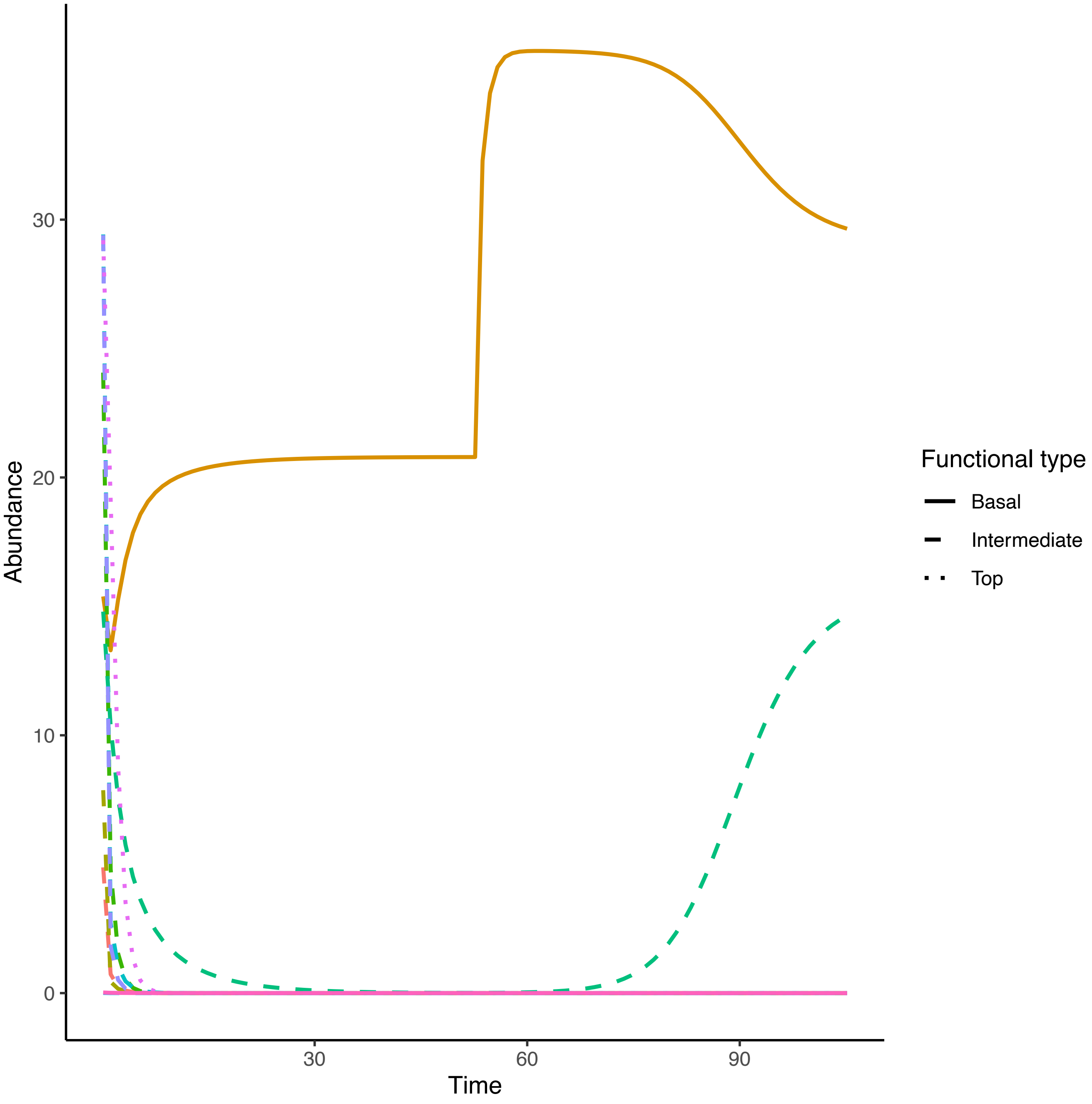

Abundance
30
20
10
0
30
60
90
Time
Functional type
Basal
Intermediate
Top

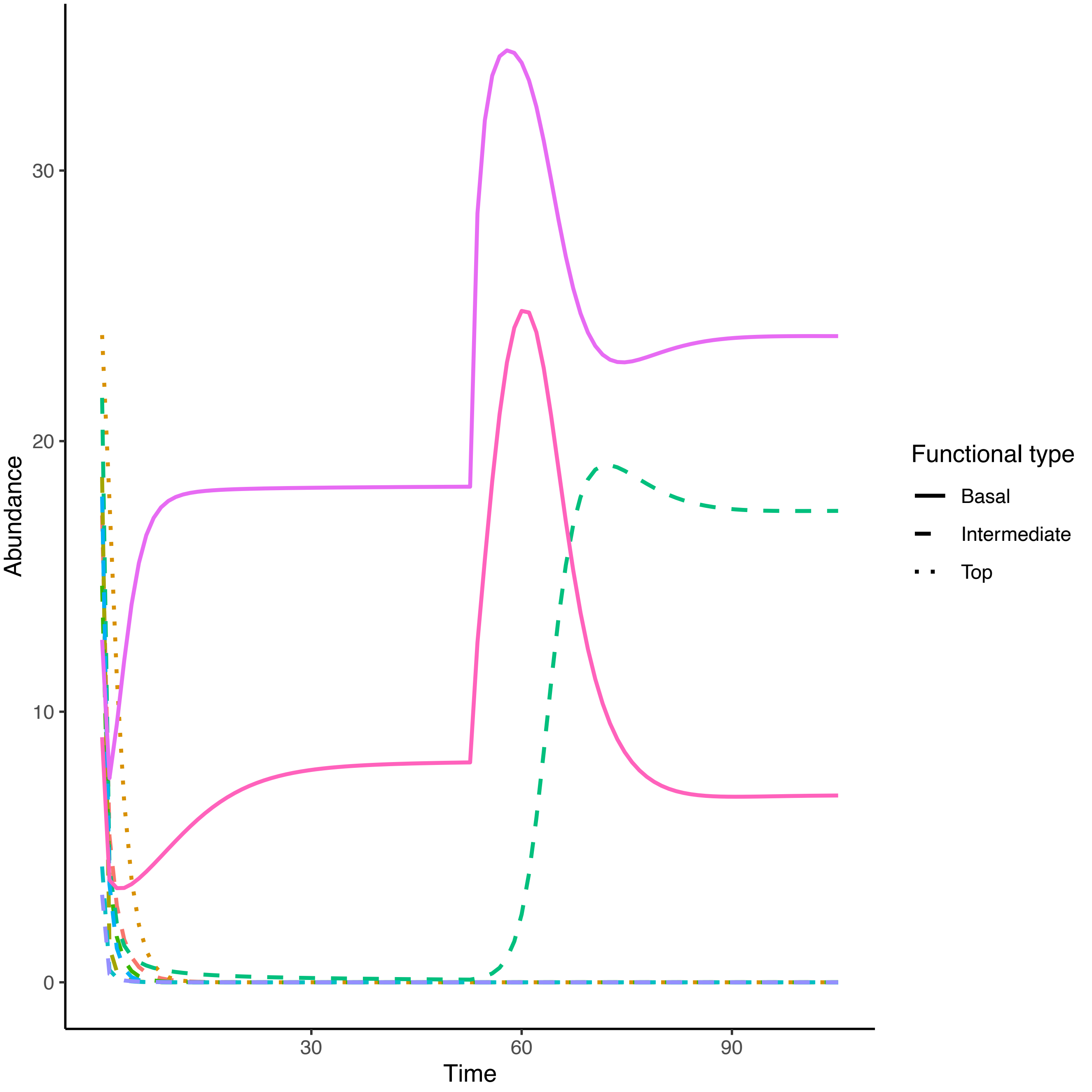

Abundance
30
20
10
0
30
60
90
Time
Functional type
Basal
Intermediate
Top

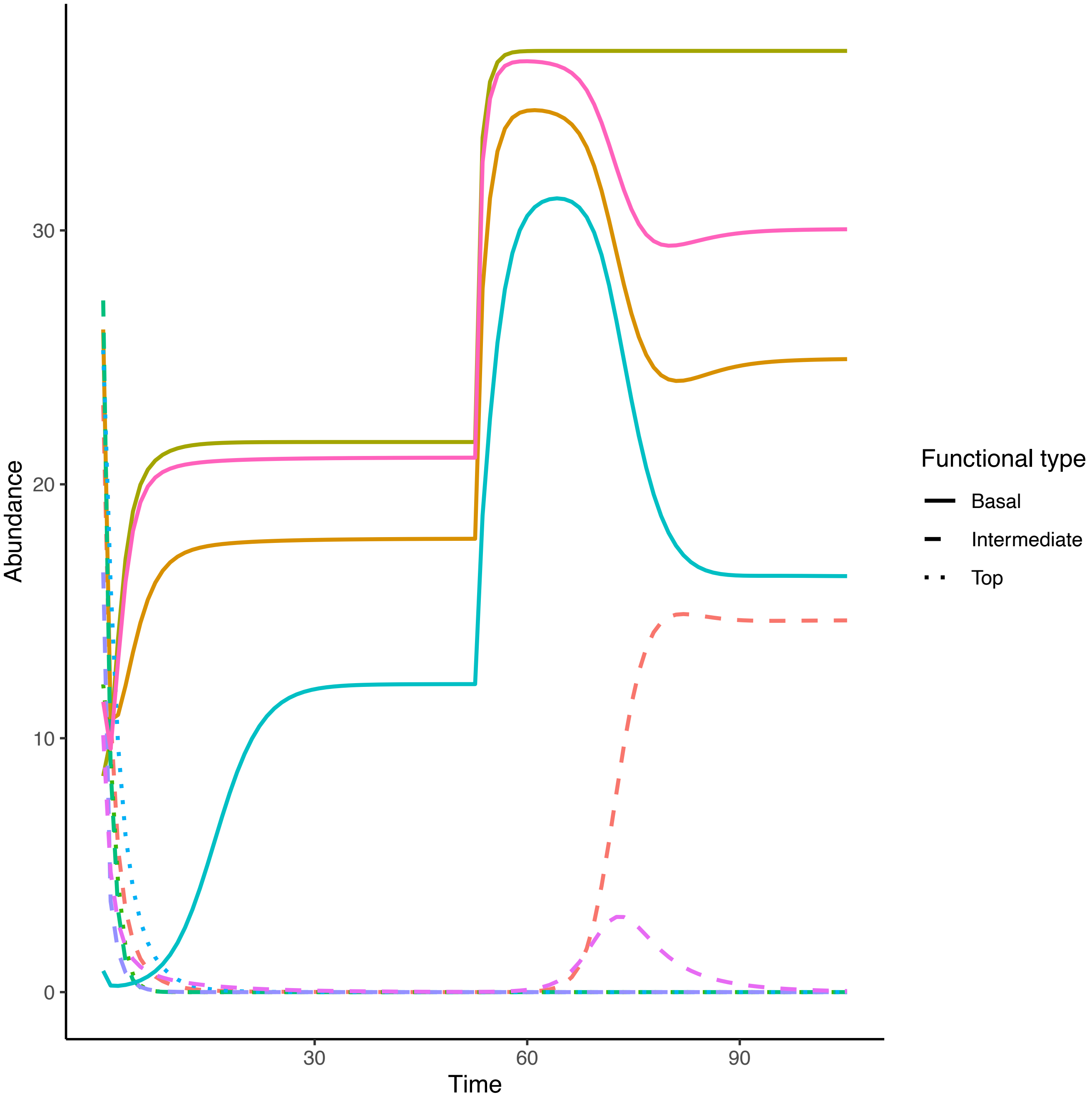

Abundance
Time
0
10
20
30
30
60
90
Functional type
Basal
Intermediate
Top

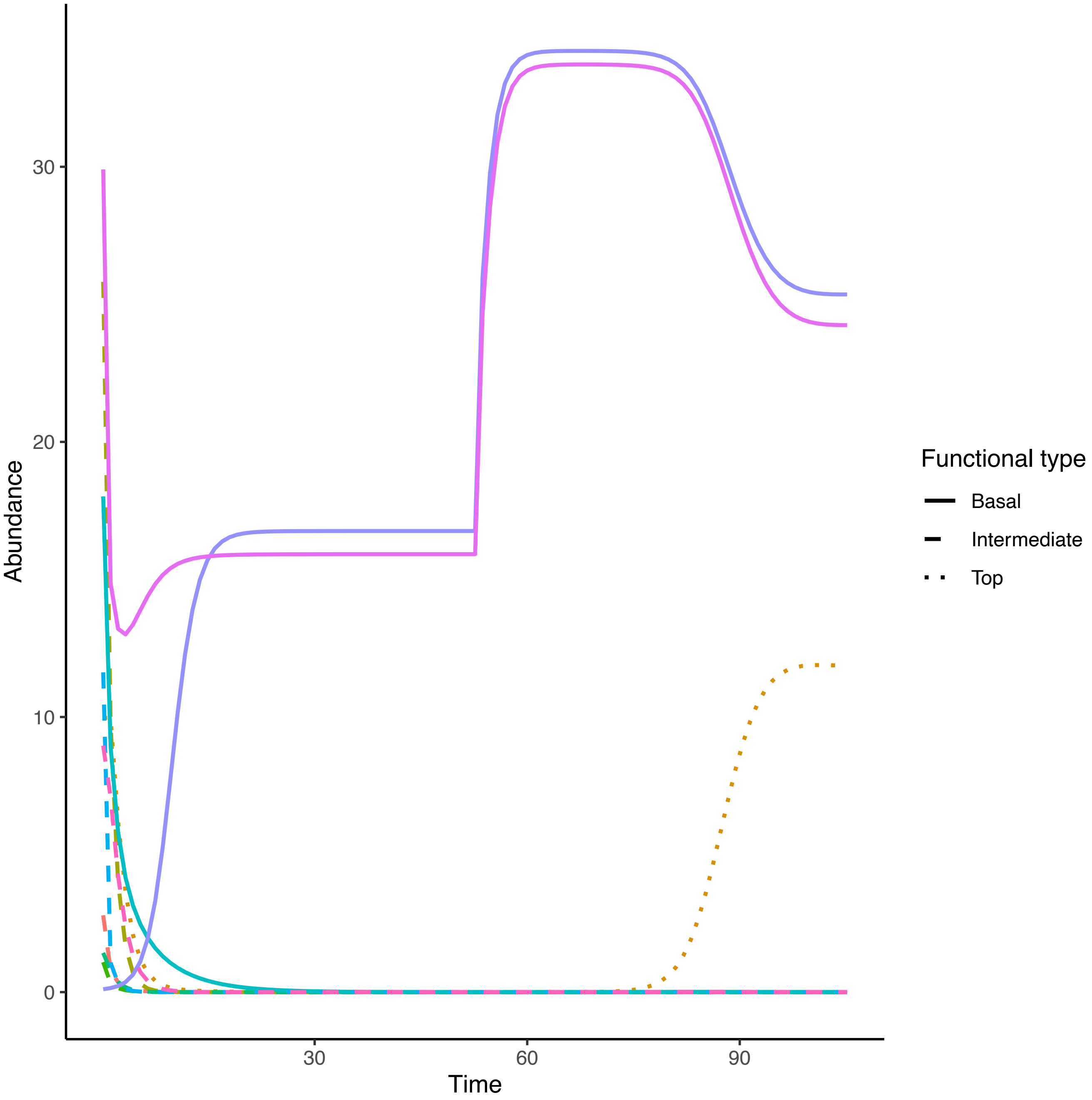

Abundance
0
10
20
30
Time
30
60
90
Functional type
Basal
Intermediate
Top

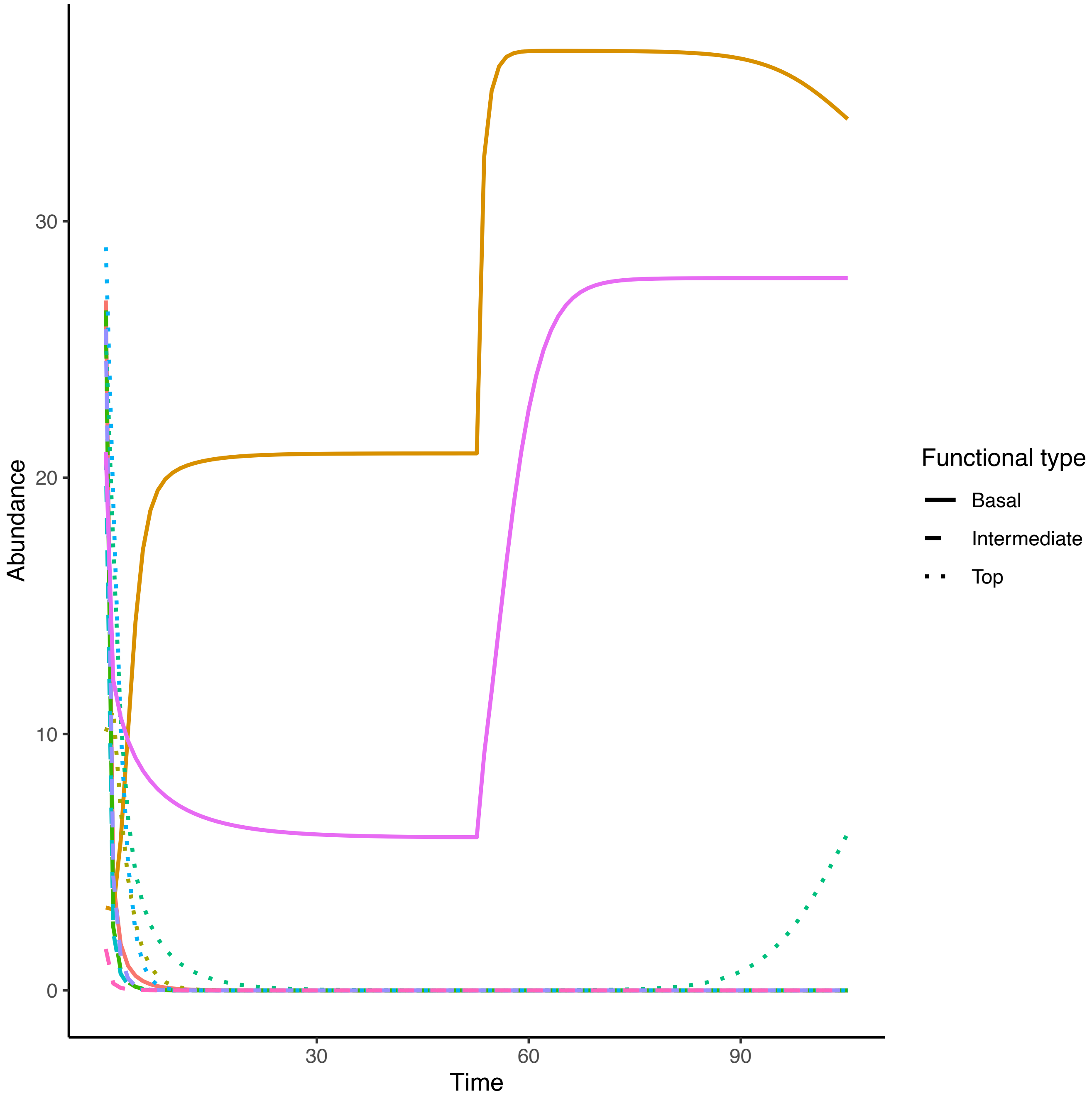

Abundance
30
20
10
0
Time
30
60
90
Functional type
Basal
Intermediate
Top

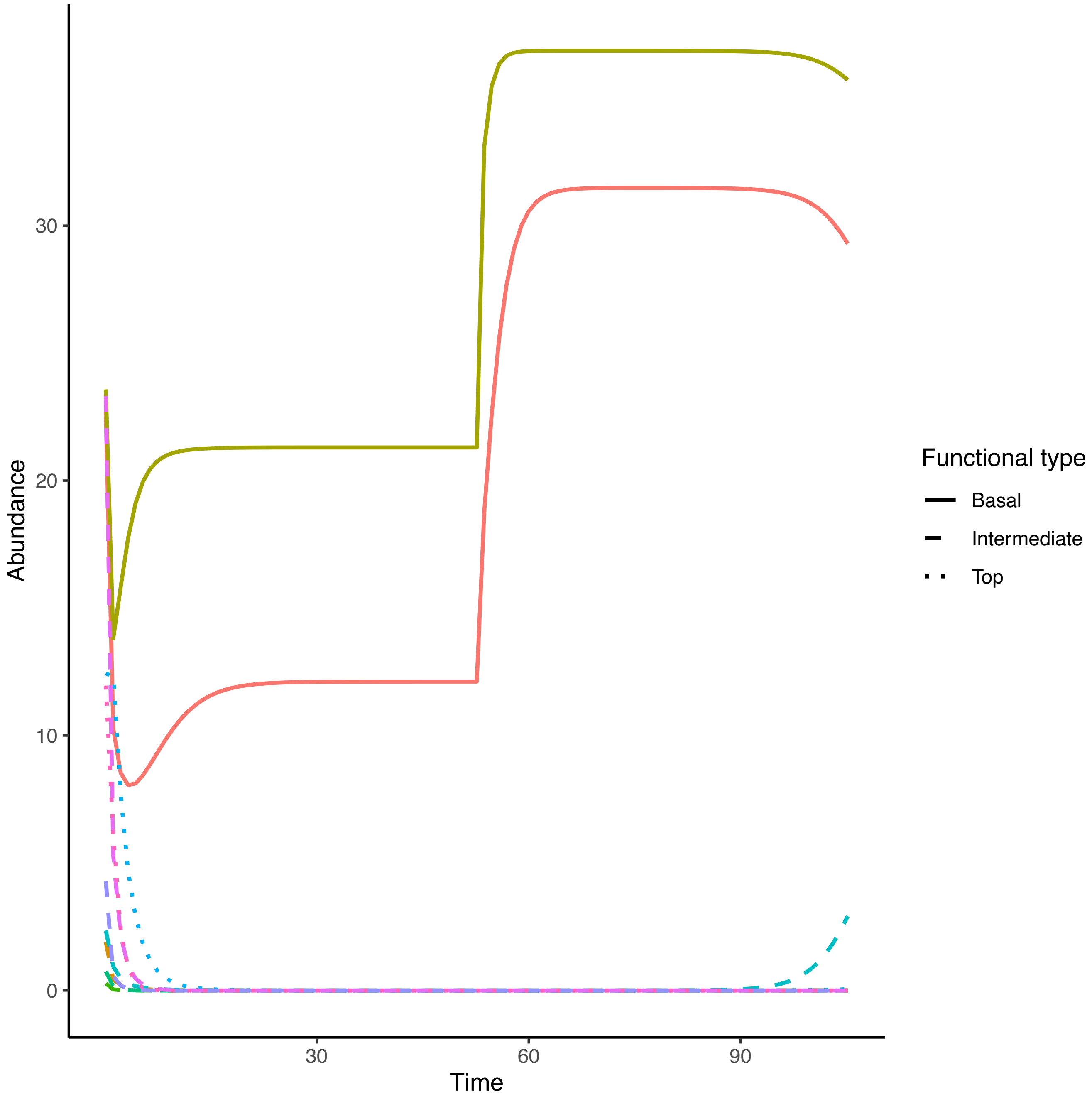

Abundance
Time
0
10
20
30
30
60
90
Functional type
Basal
Intermediate
Top

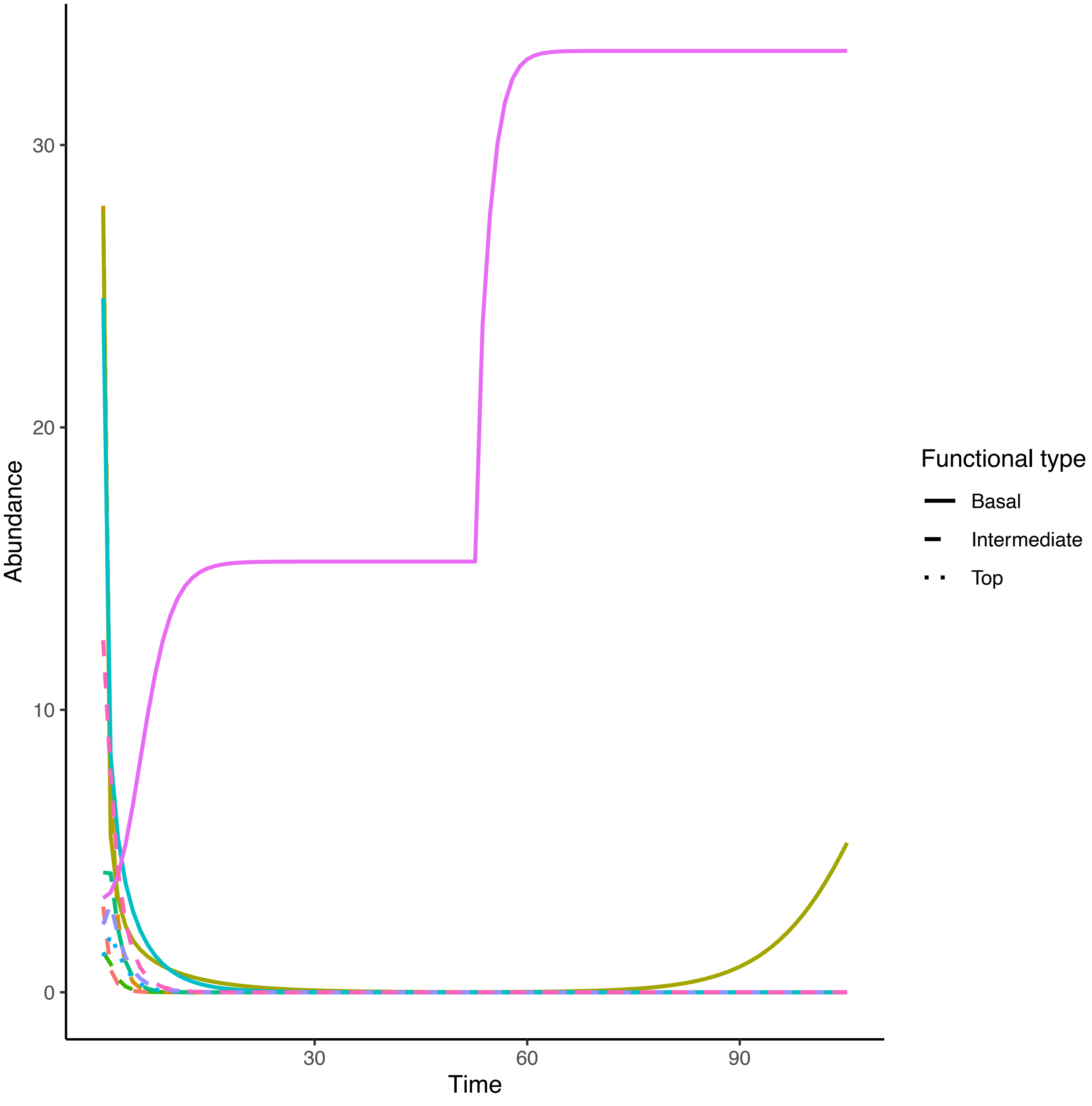

30
20
10
0
Abundance
30
60
90
Time
Functional type
Basal
Intermediate
Top

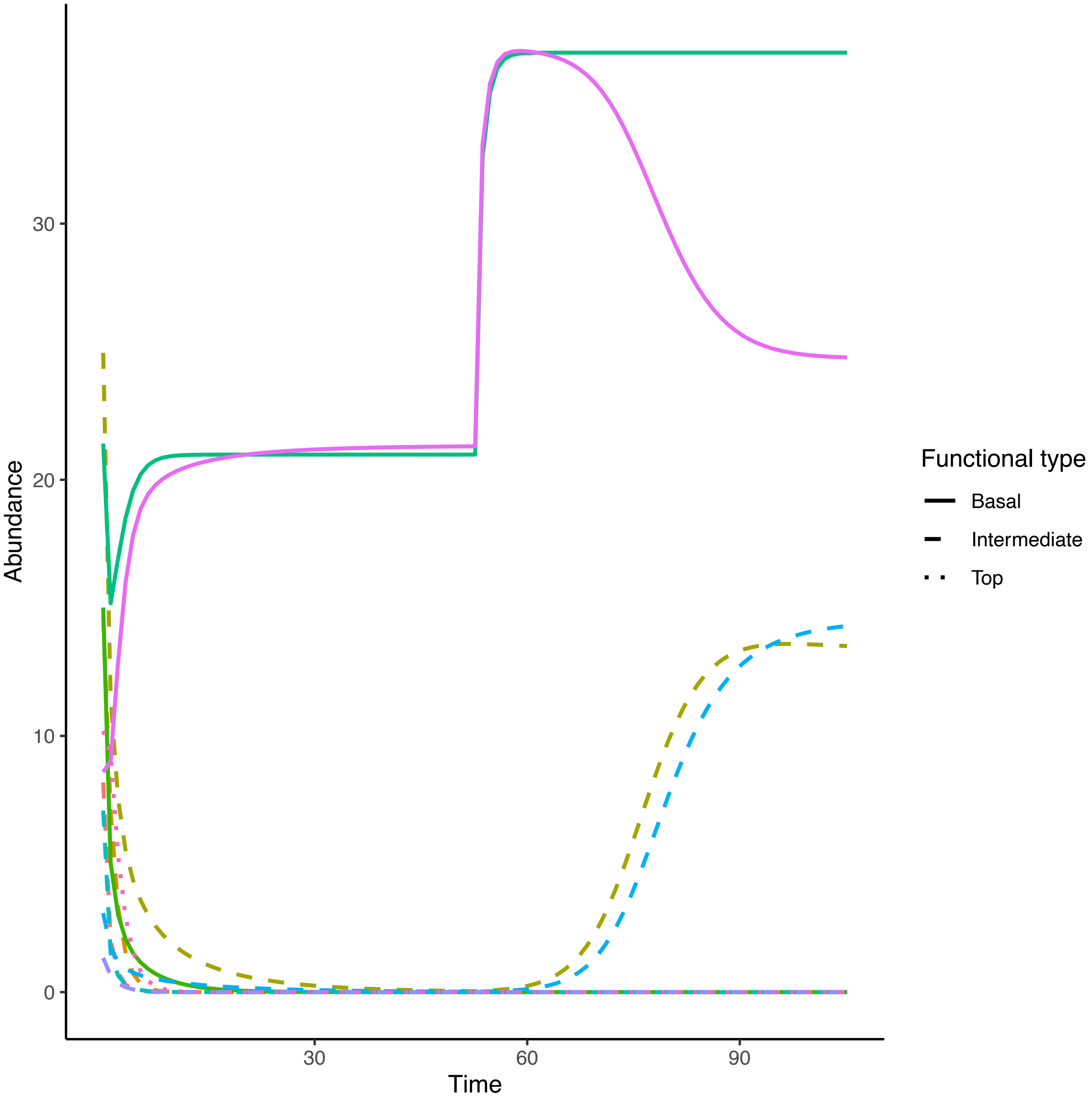

Abundance
Time
0
10
20
30
30
60
90
Functional type
Basal
Intermediate
Top

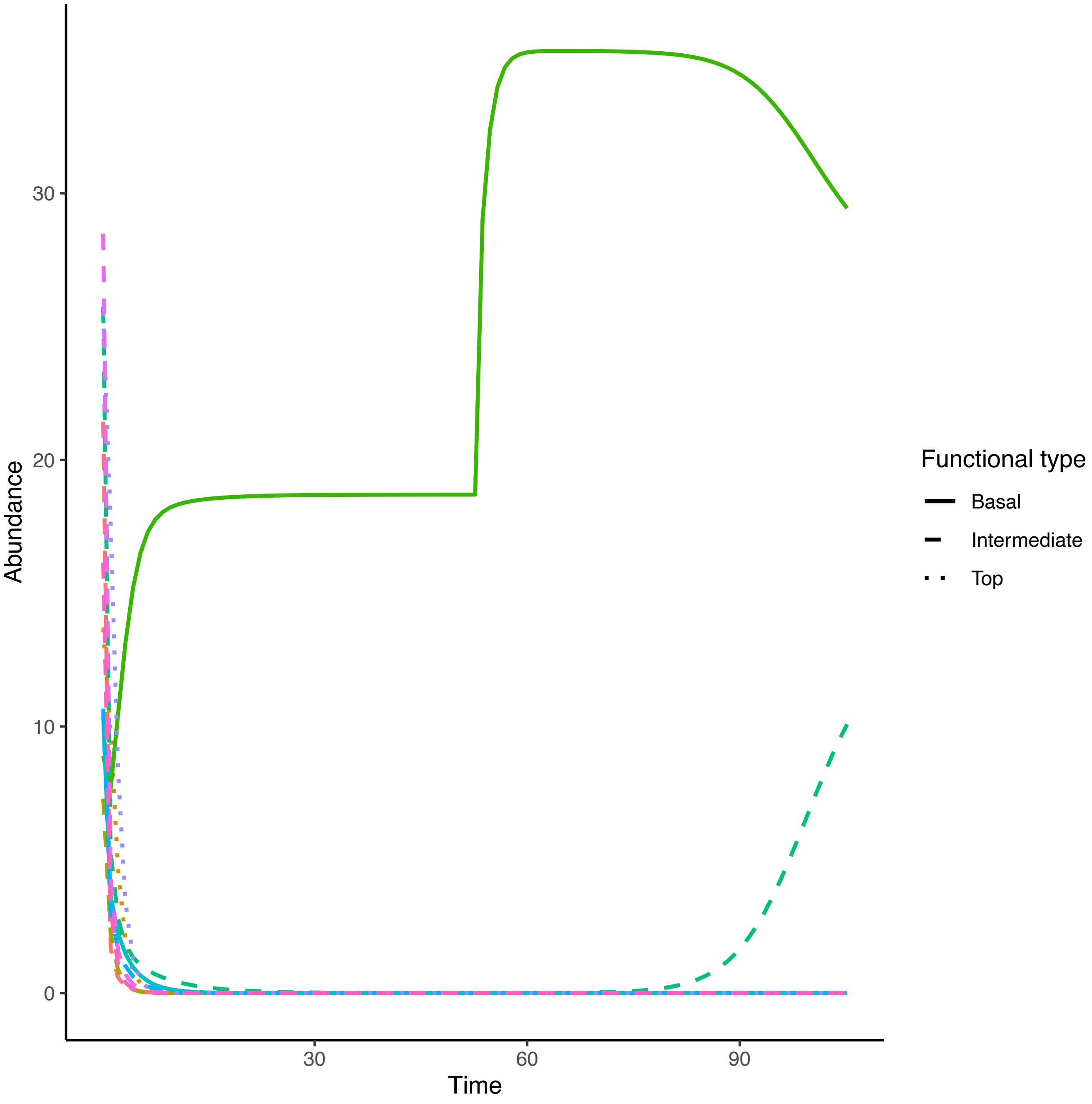

Abundance
Time
0
10
20
30
30
60
90
Functional type
Basal
Intermediate
Top

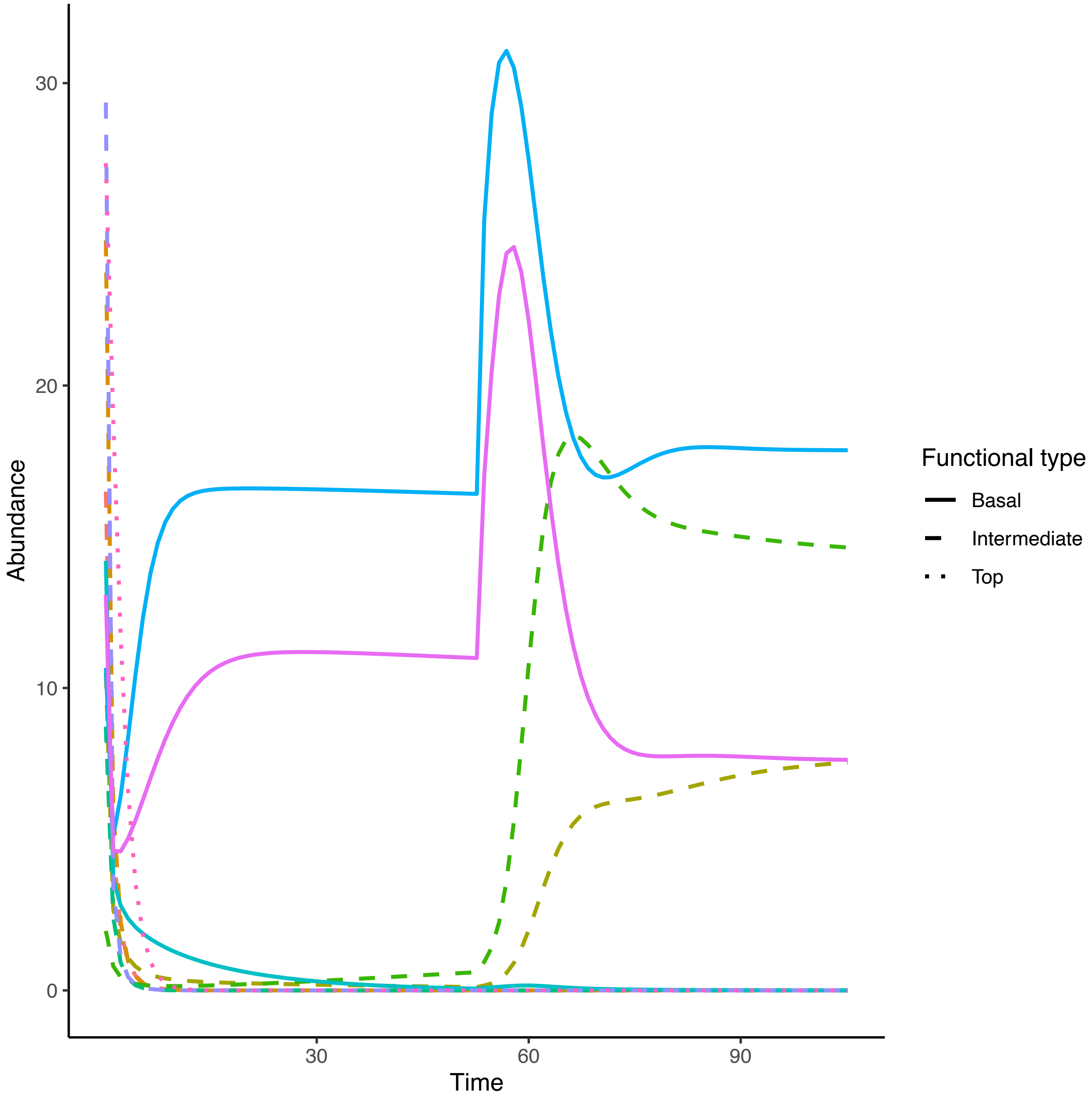

Abundance
30
20
10
0
30
60
90
Time
Functional type
Basal
Intermediate
Top

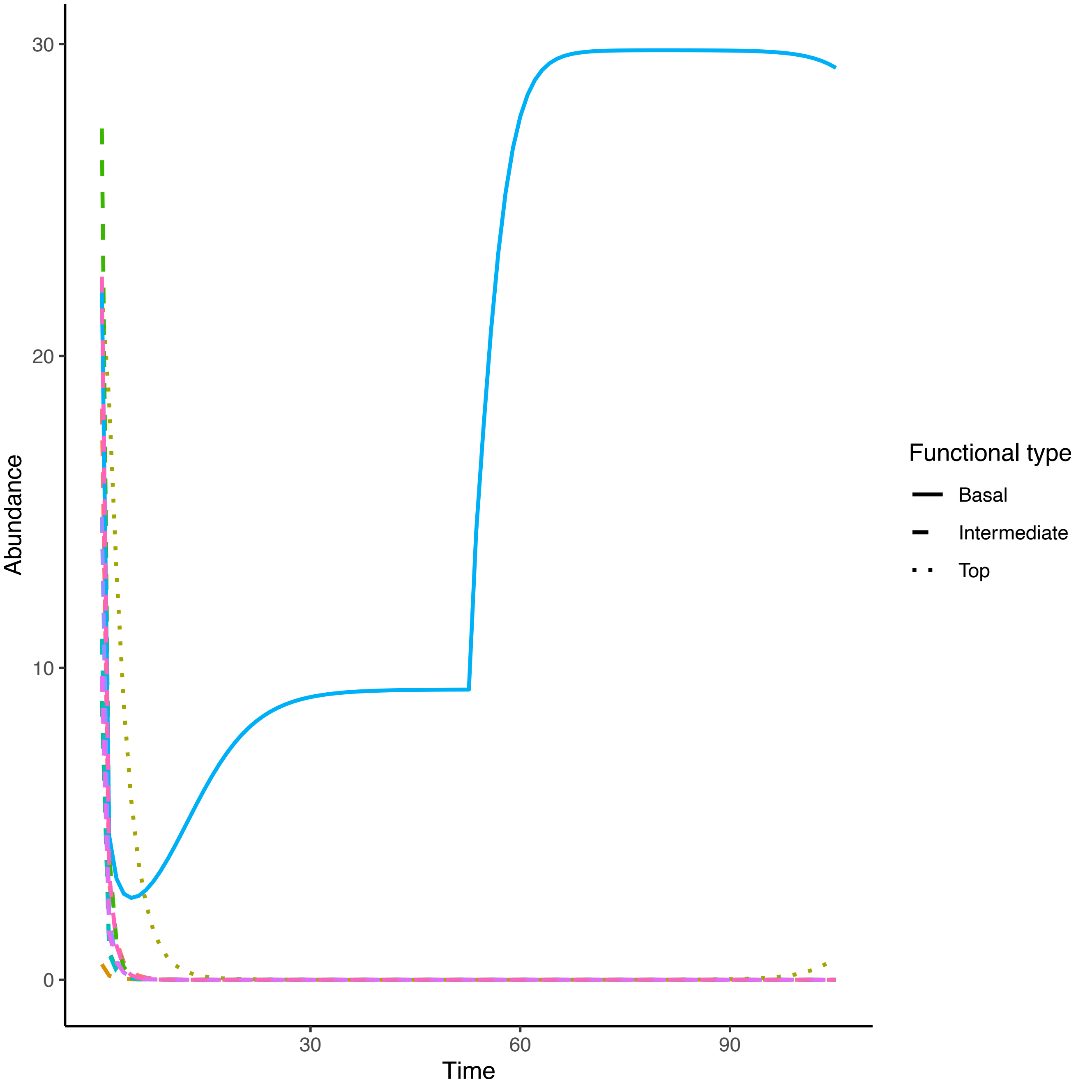
30
20
10
0
Abundance
30
60
90
Time
Functional type
Basal
Intermediate
Top

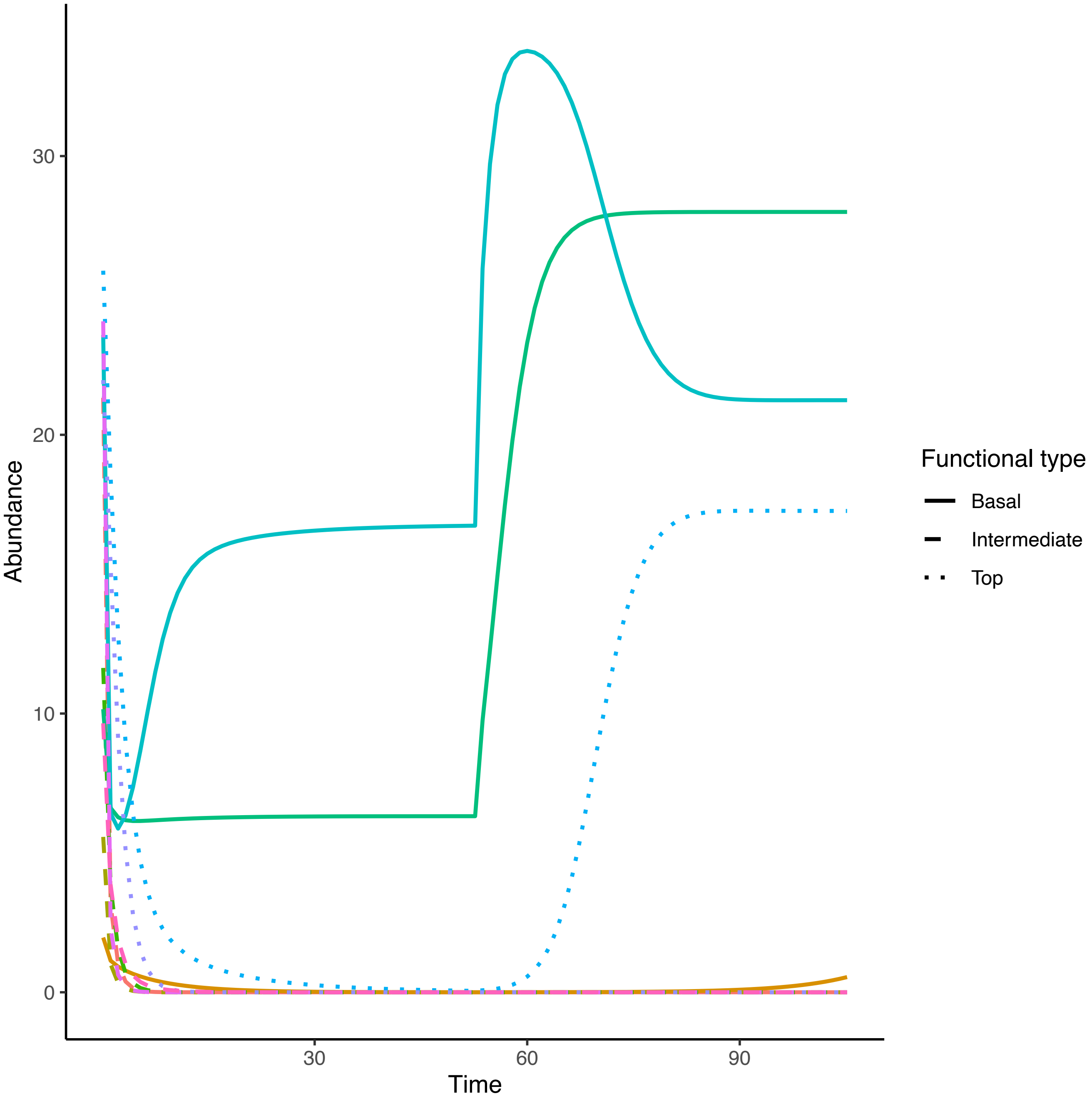

30
20
10
0
Abundance
30
60
90
Time
Functional type
Basal
Intermediate
Top

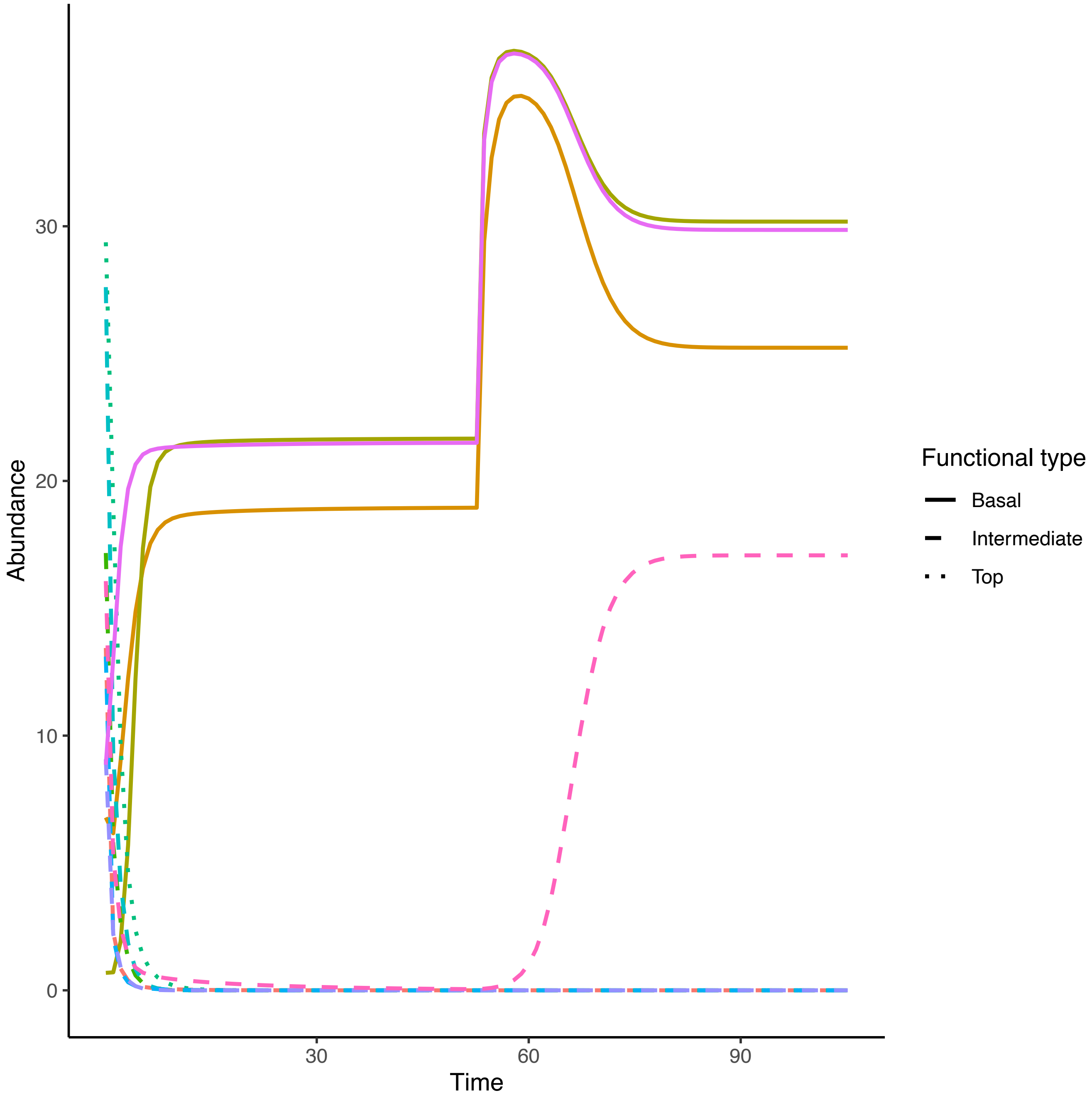

Abundance
30
20
10
0
Time
30
60
90
Functional type
Basal
Intermediate
Top

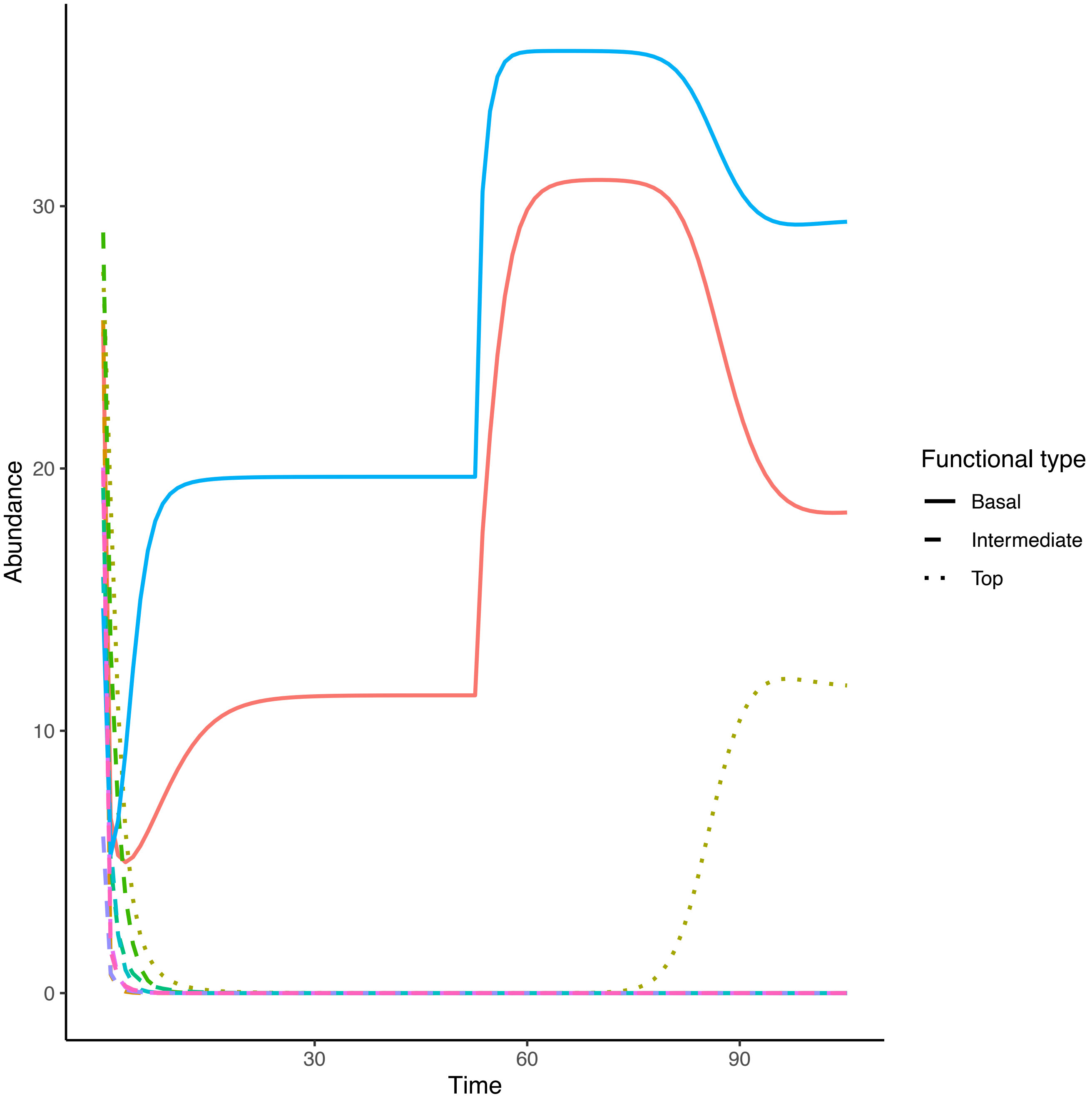

Abundance
30
20
10
0
30
60
90
Time
Functional type
Basal
Intermediate
Top

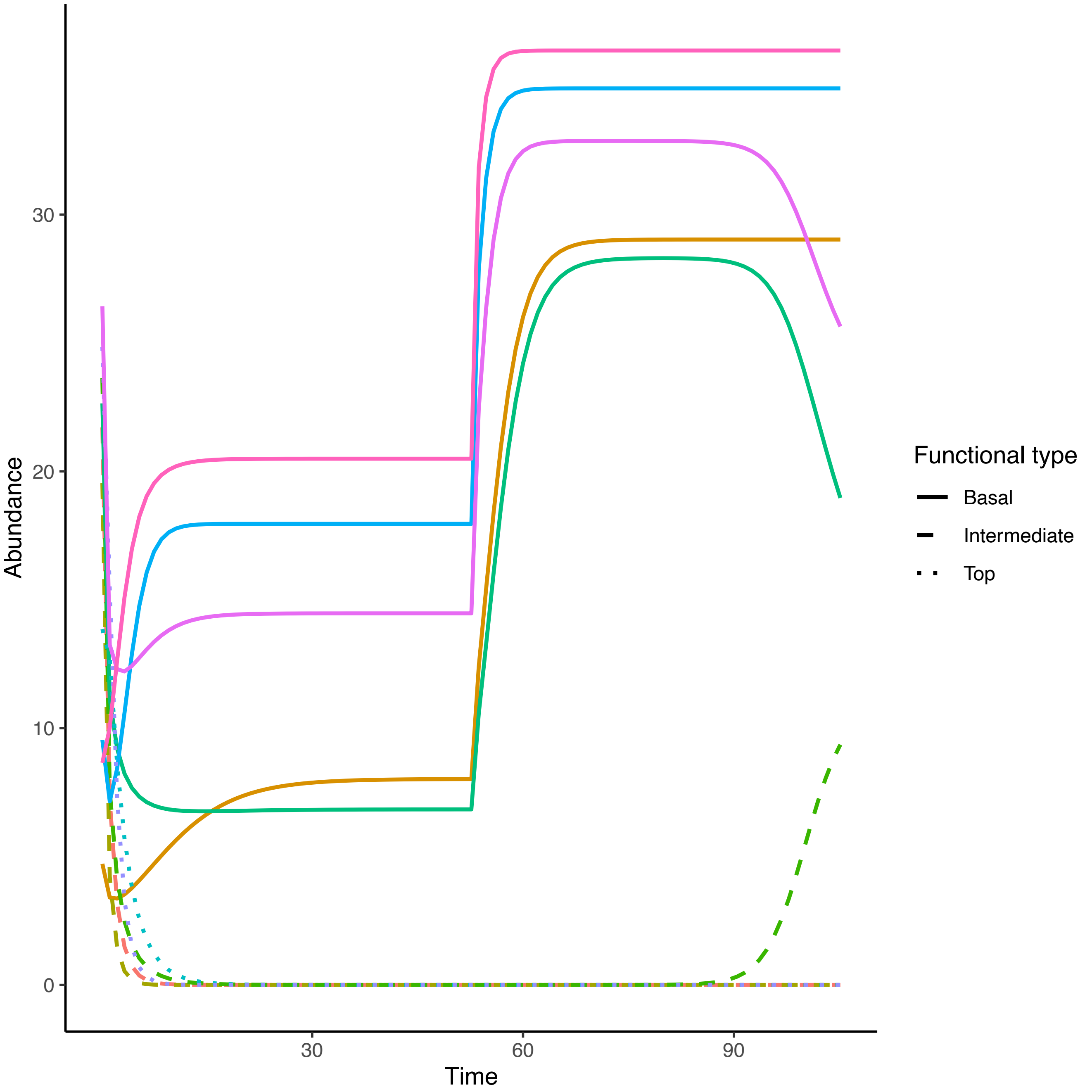
Abundance
Time
0
10
20
30
30
60
90
Functional type
Basal
Intermediate
Top

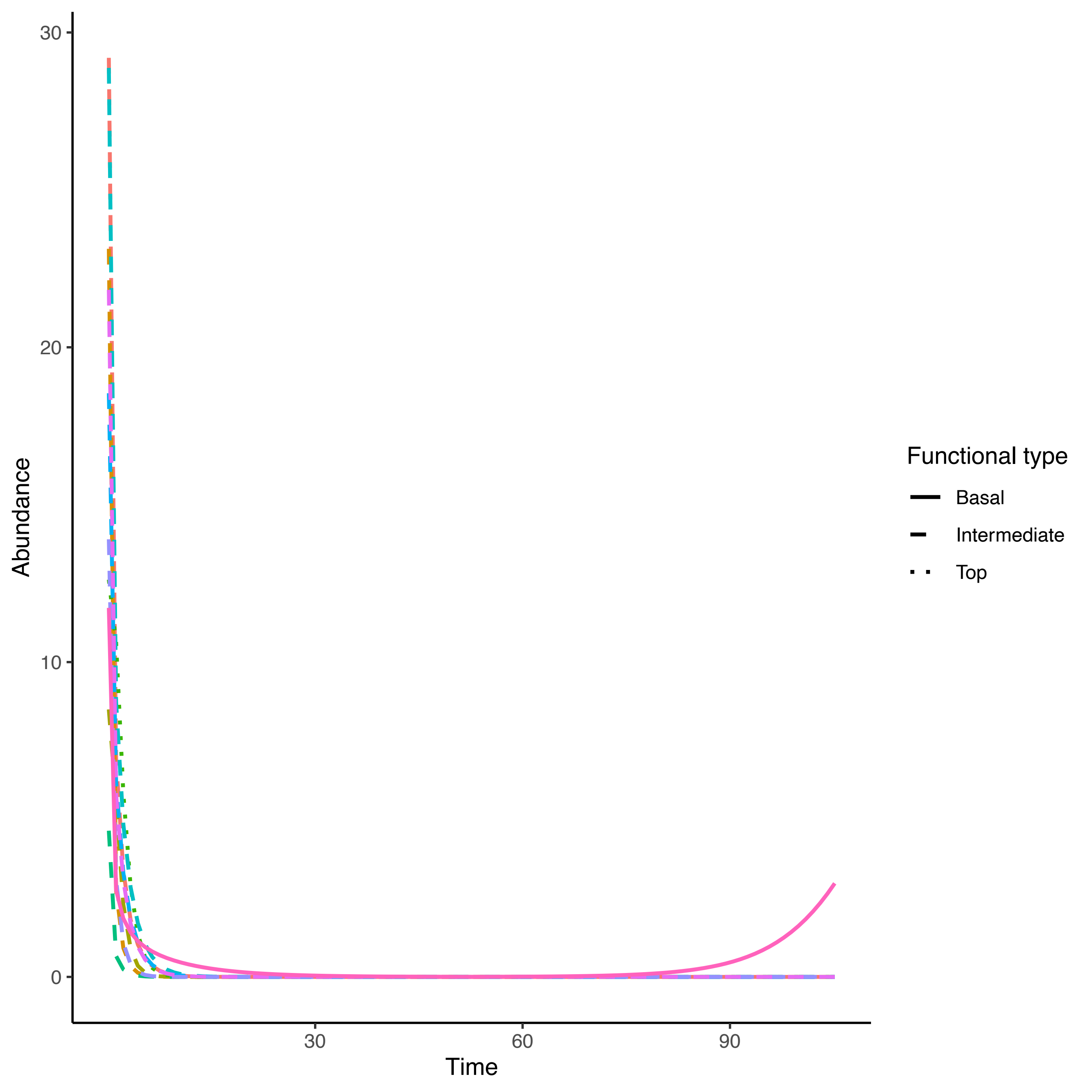

30
20
10
0
Abundance
30
60
90
Time
Functional type
Basal
Intermediate
Top

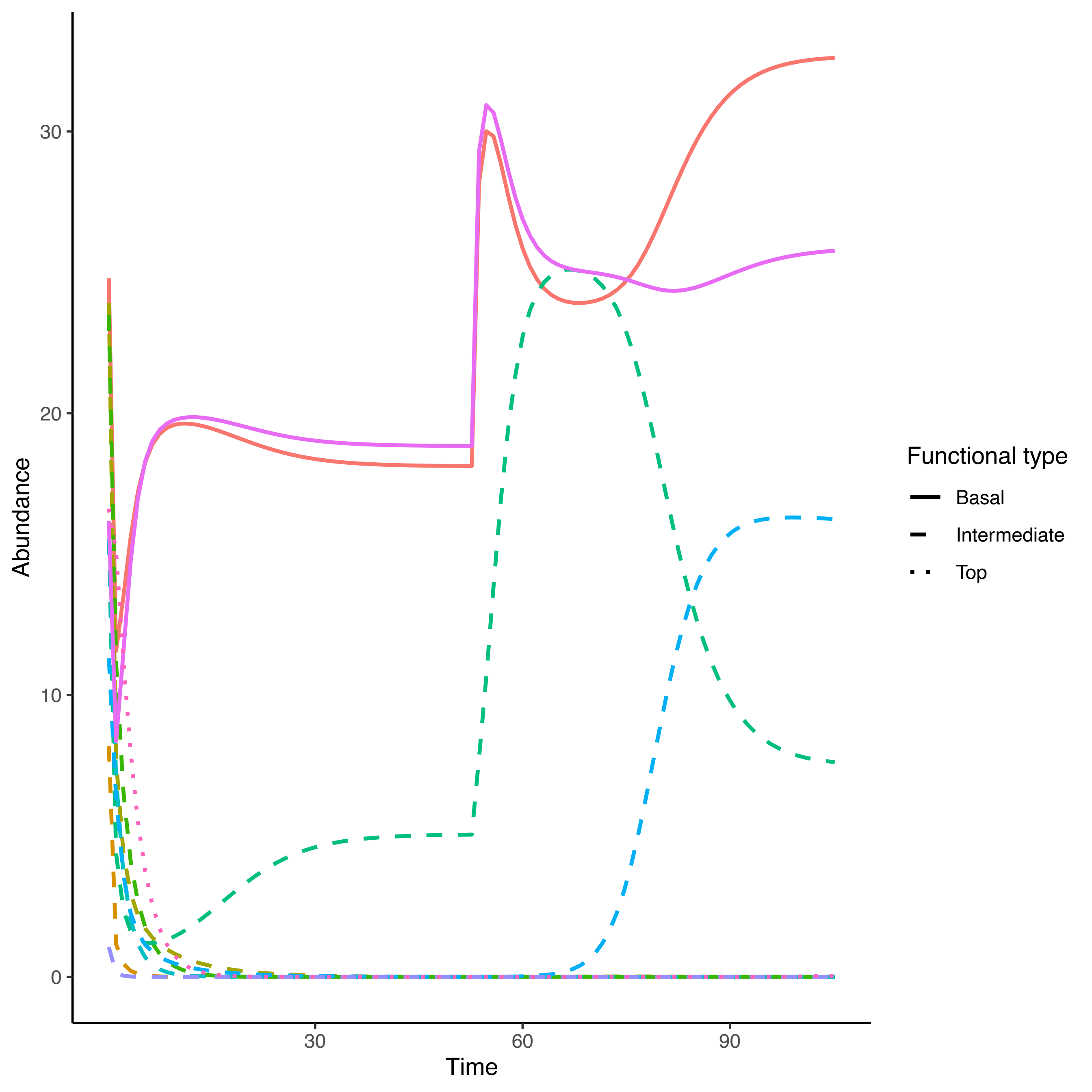
Abundance
0
10
20
30
Time
30
60
90
Functional type
Basal
Intermediate
Top

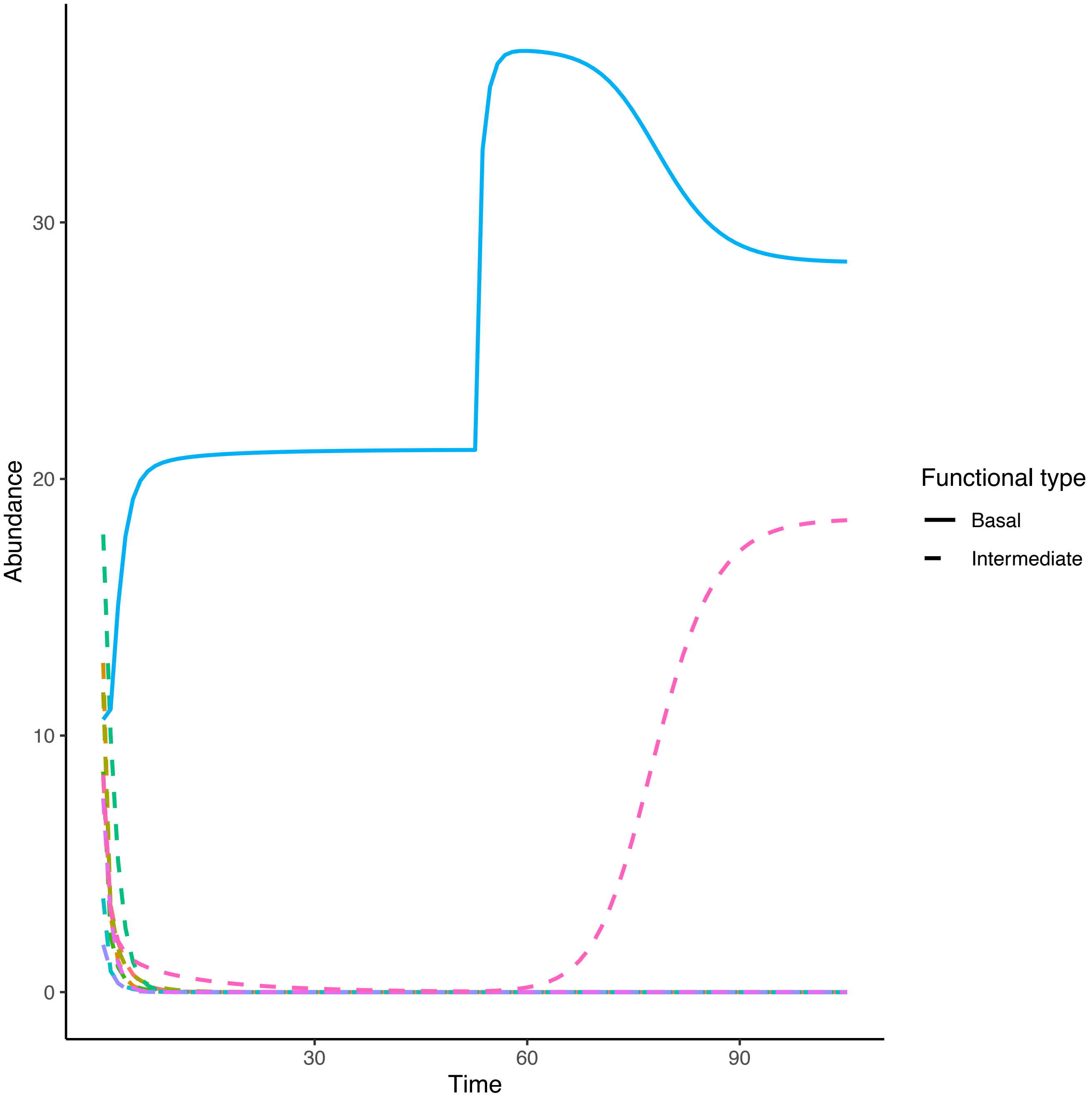

Abundance
30
20
10
0
30
60
90
Time
Functional type
Basal
Intermediate

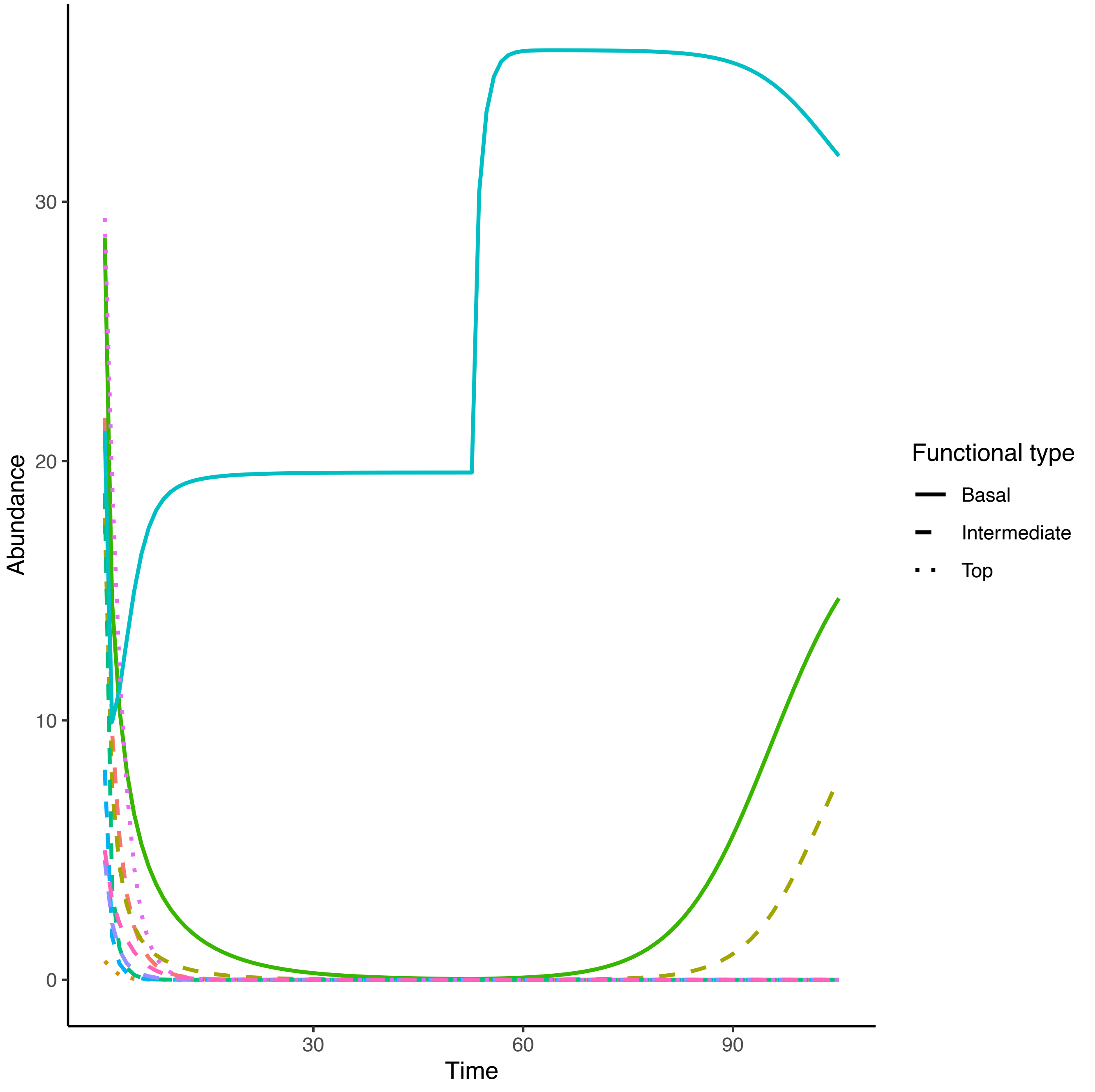

Abundance
30
20
10
0
30
60
90
Time
Functional type
Basal
Intermediate
Top

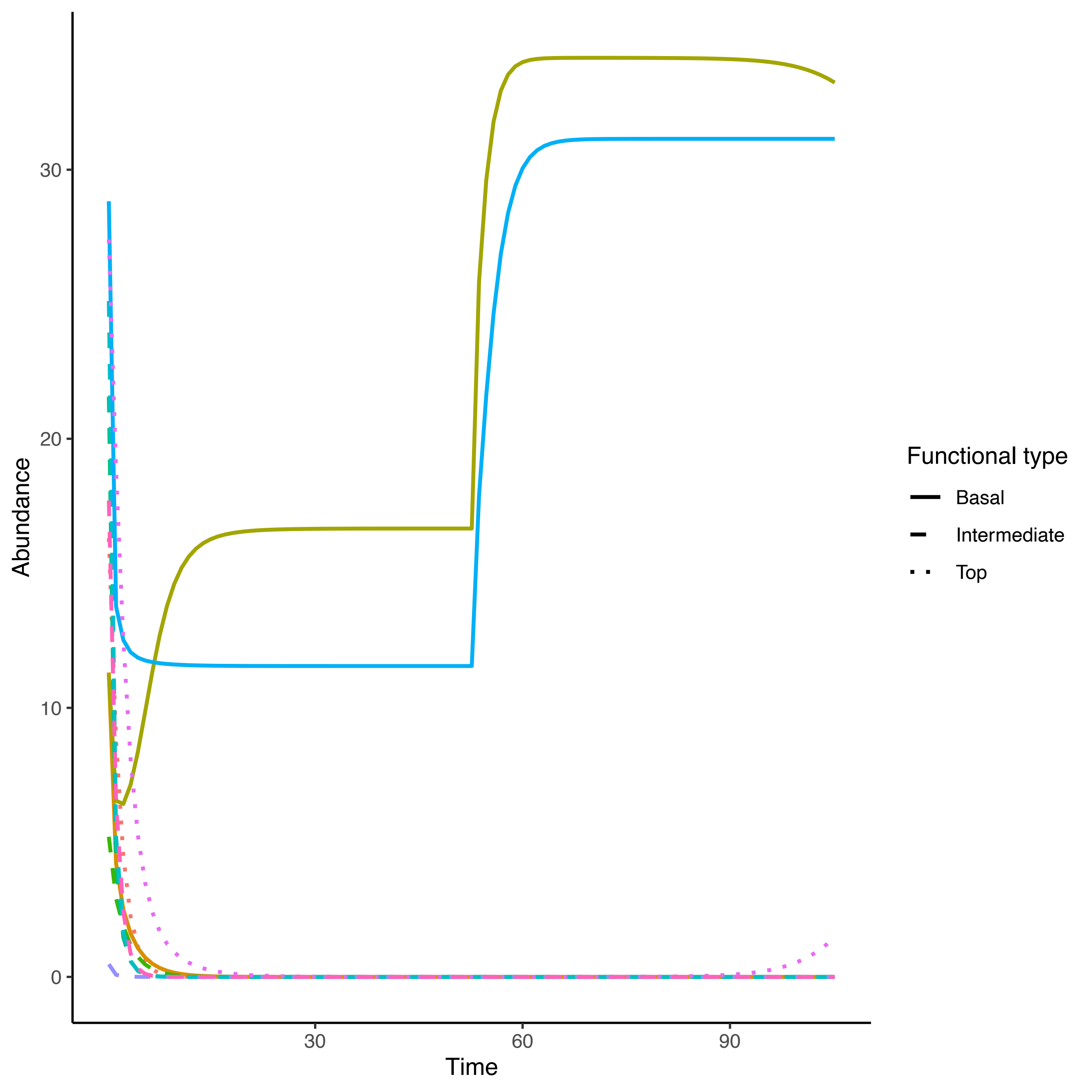

Abundance
30
20
10
0
30
60
90
Time
Functional type
Basal
Intermediate
Top

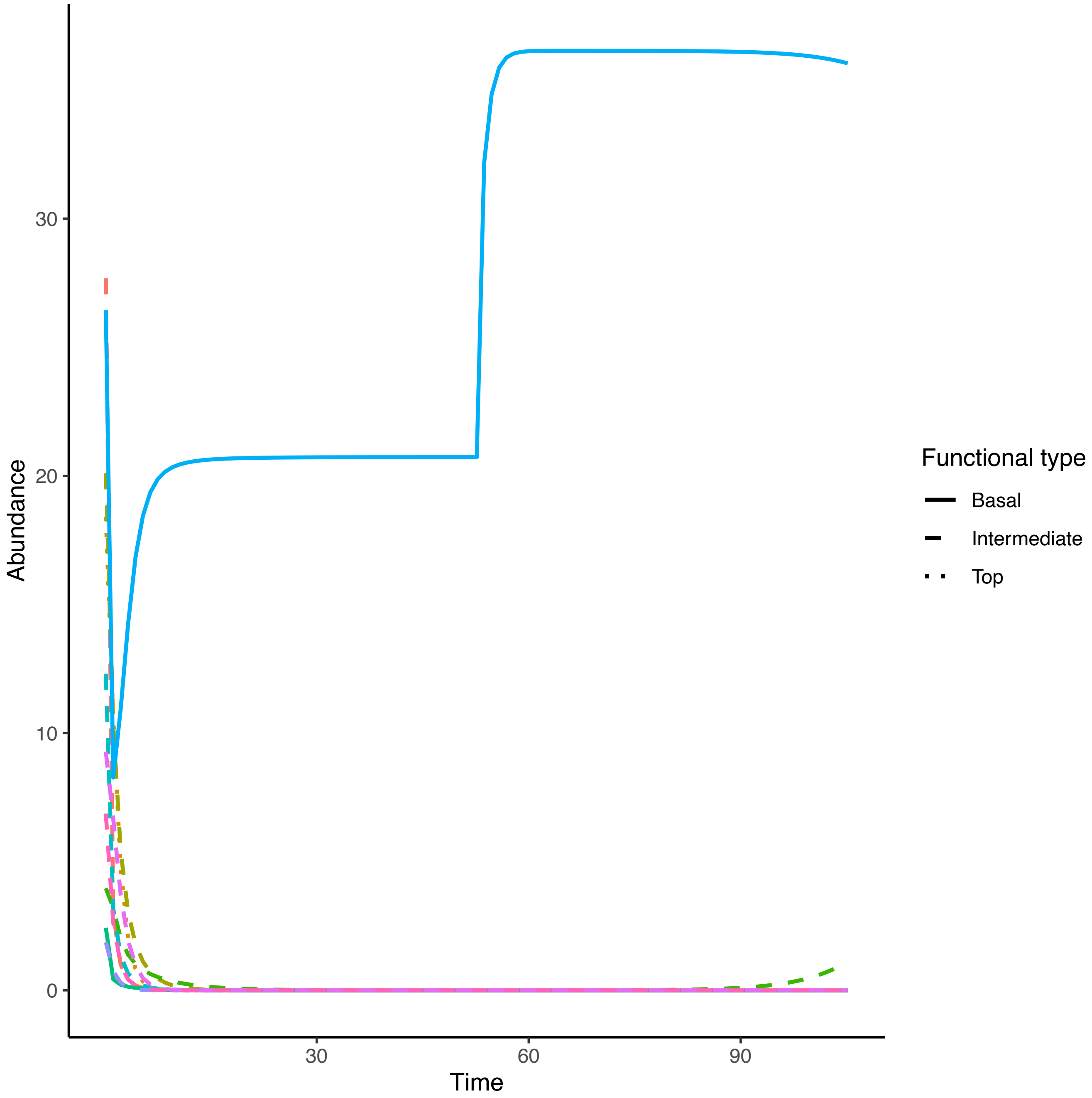

Abundance
30
20
10
0
Time
30
60
90
Functional type
Basal
Intermediate
Top

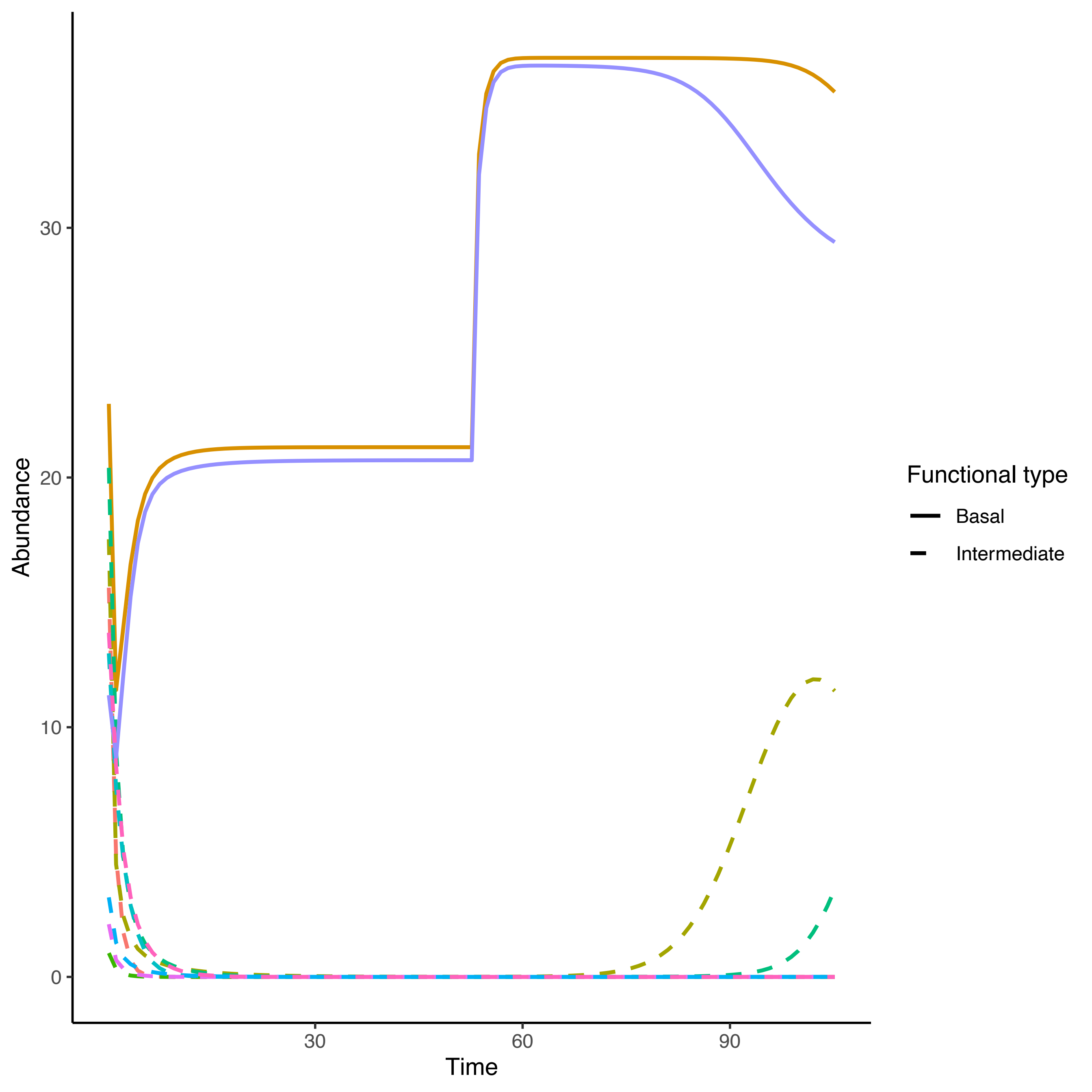

Abundance
30
20
10
0
Time
30
60
90
Functional type
Basal
Intermediate

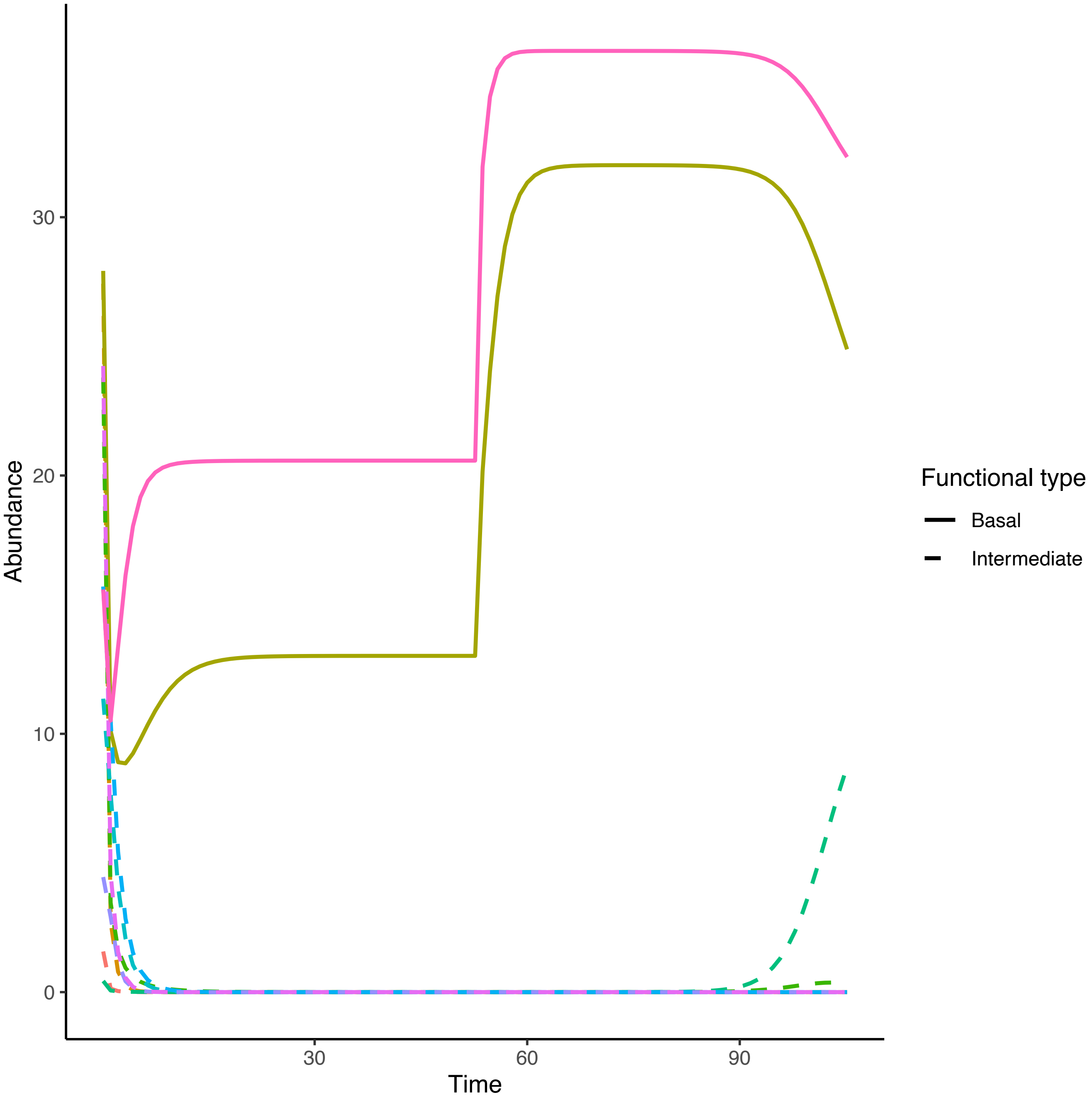

Abundance
30
20
10
0
30
60
90
Time
Functional type
Basal
Intermediate

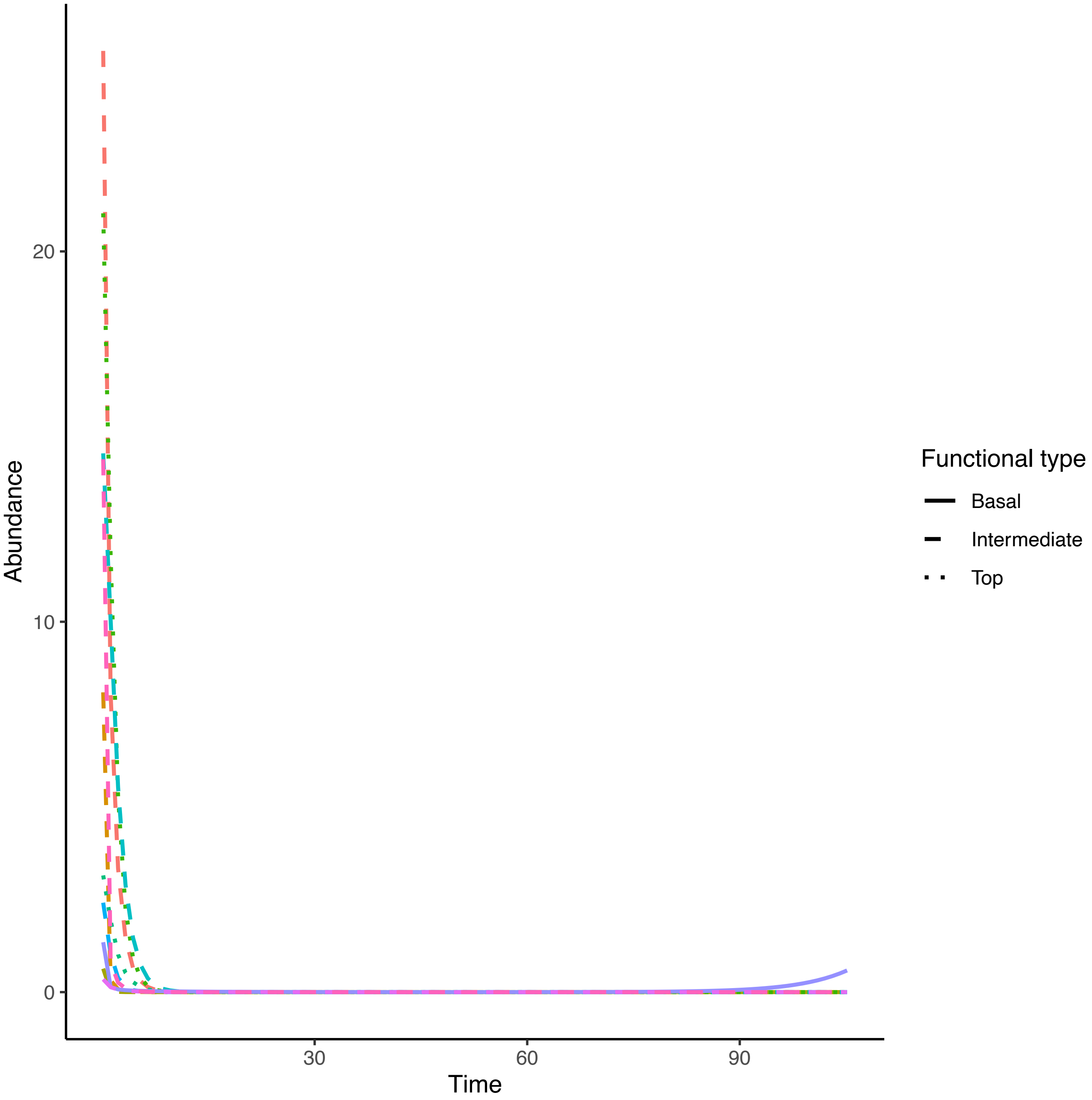

Abundance
20
10
0
30
60
90
Time
Functional type
Basal
Intermediate
Top

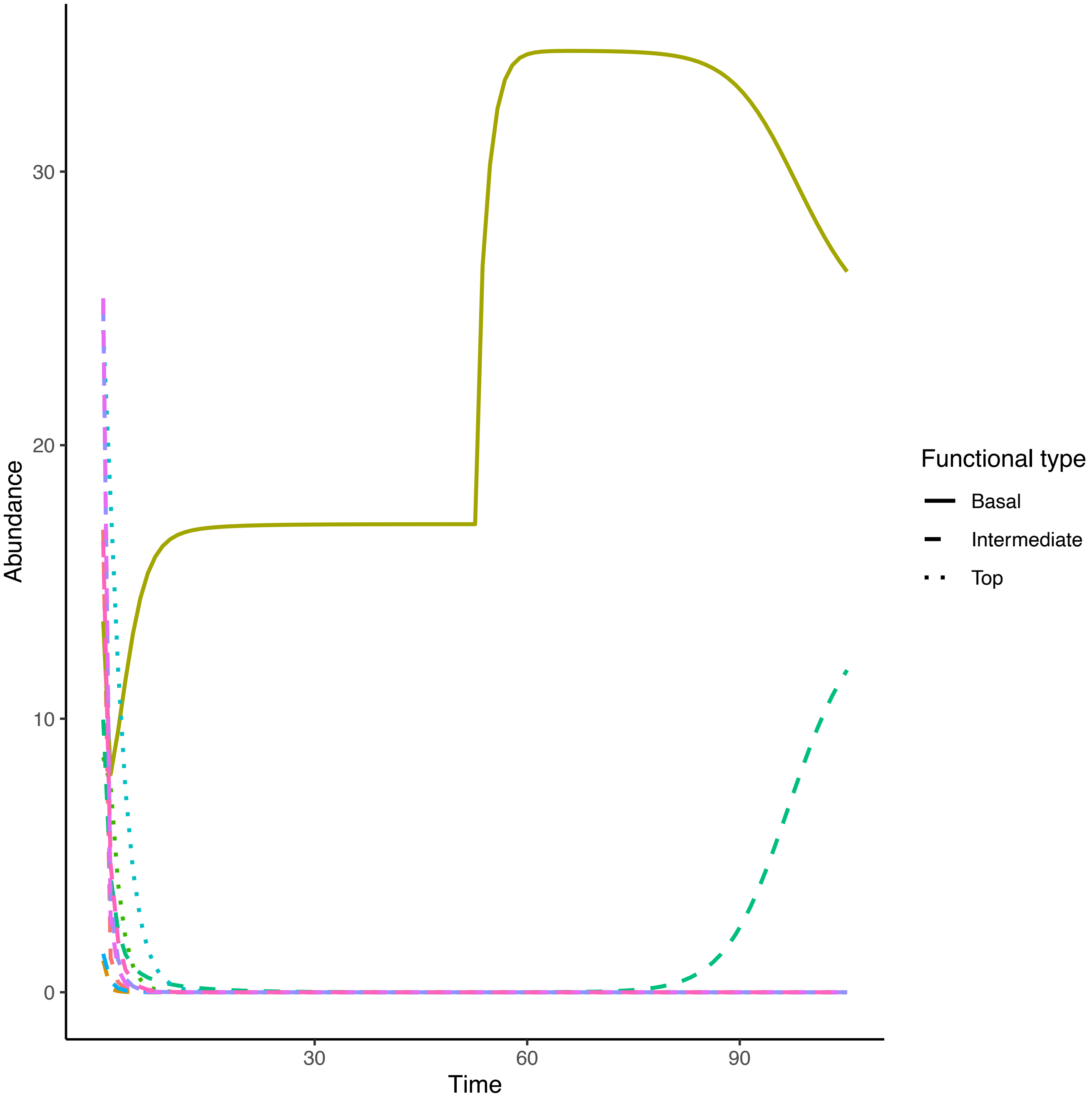

Abundance
30
20
10
0
Time
30
60
90
Functional type
Basal
Intermediate
Top

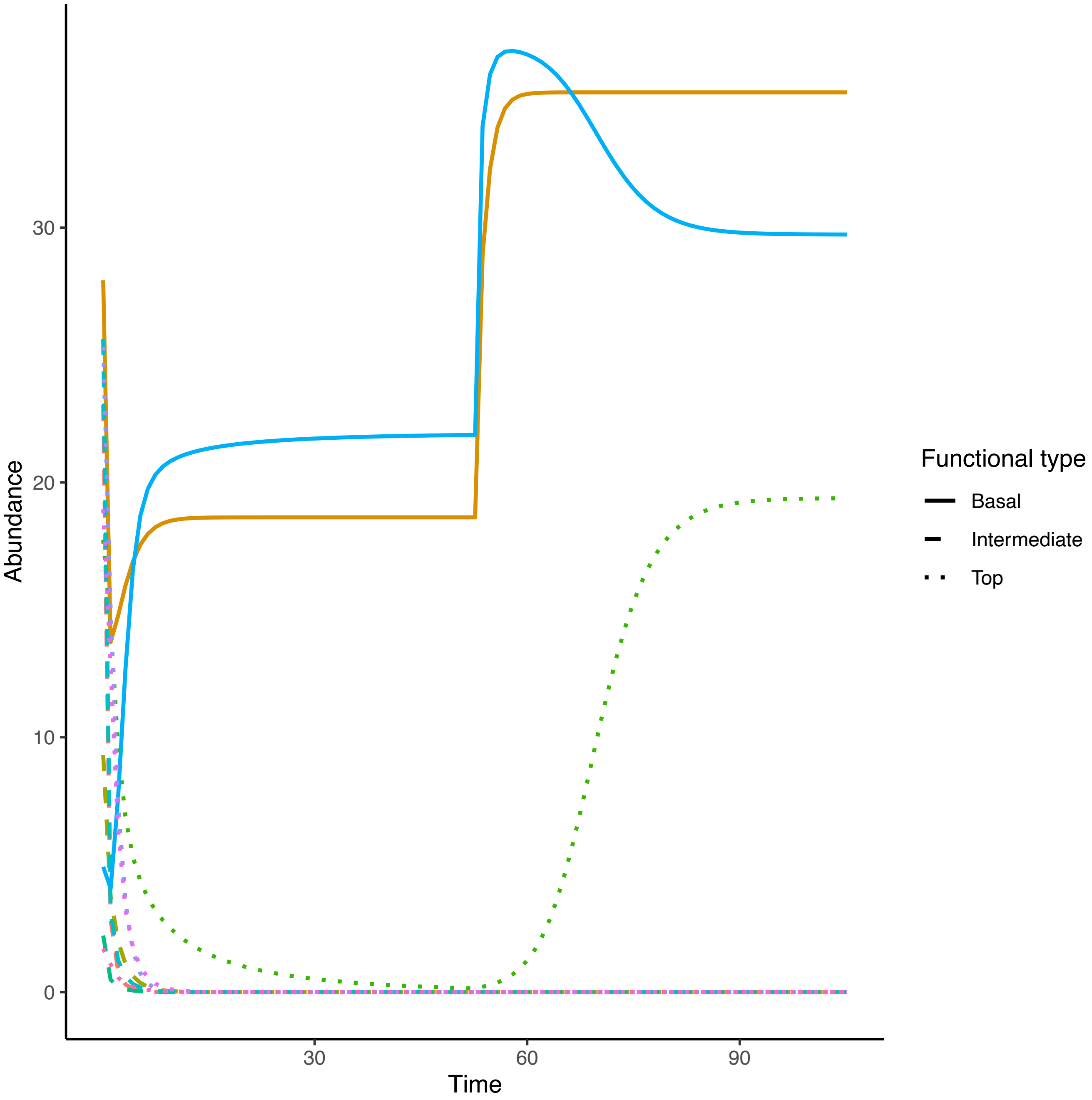
Abundance
Time
0
10
20
30
30
60
90
Functional type
Basal
Intermediate
Top

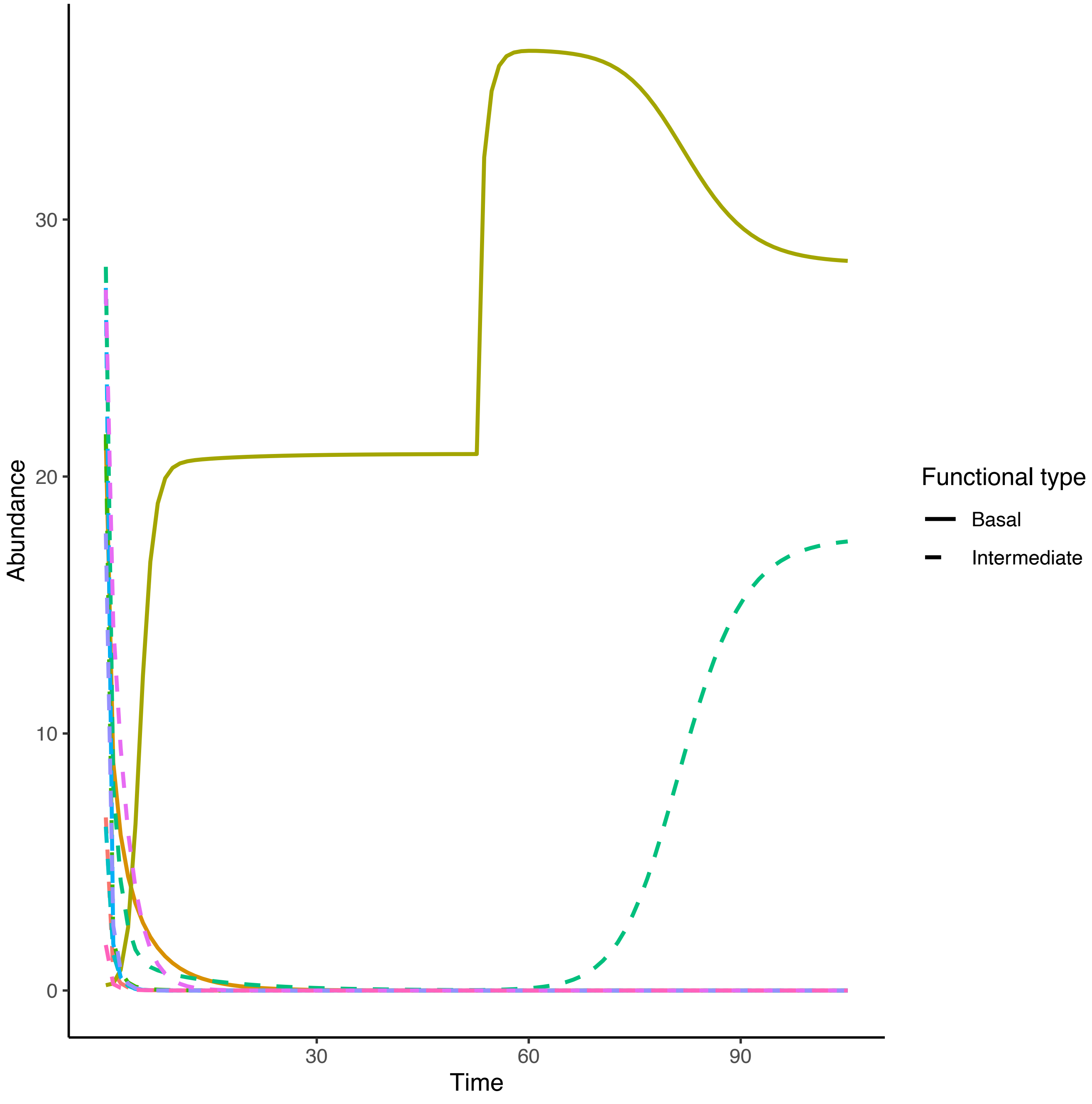

Abundance
Time
0
10
20
30
30
60
90
Functional type
Basal
Intermediate

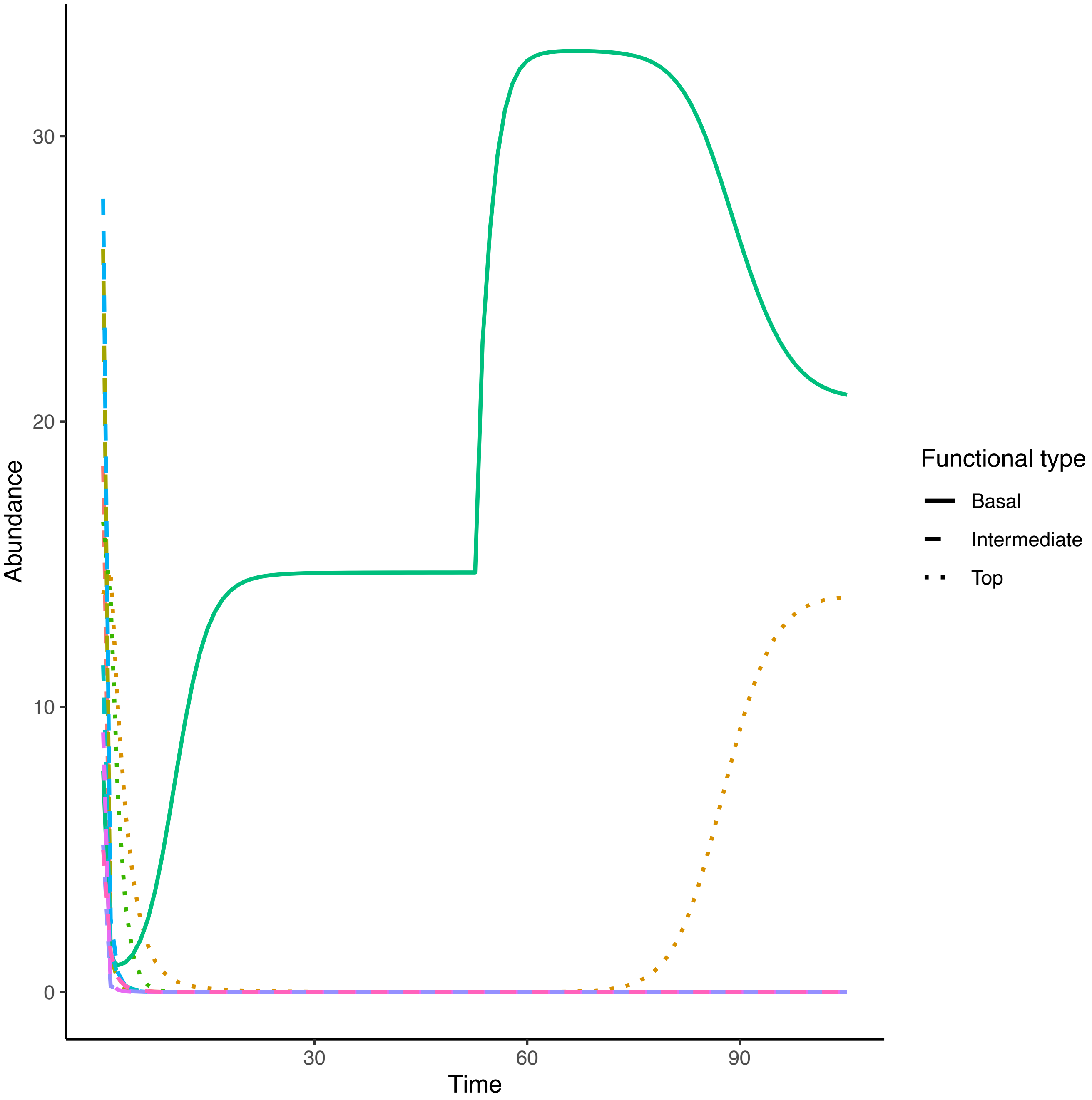

30
20
10
0
Abundance
30
60
90
Time
Functional type
Basal
Intermediate
Top

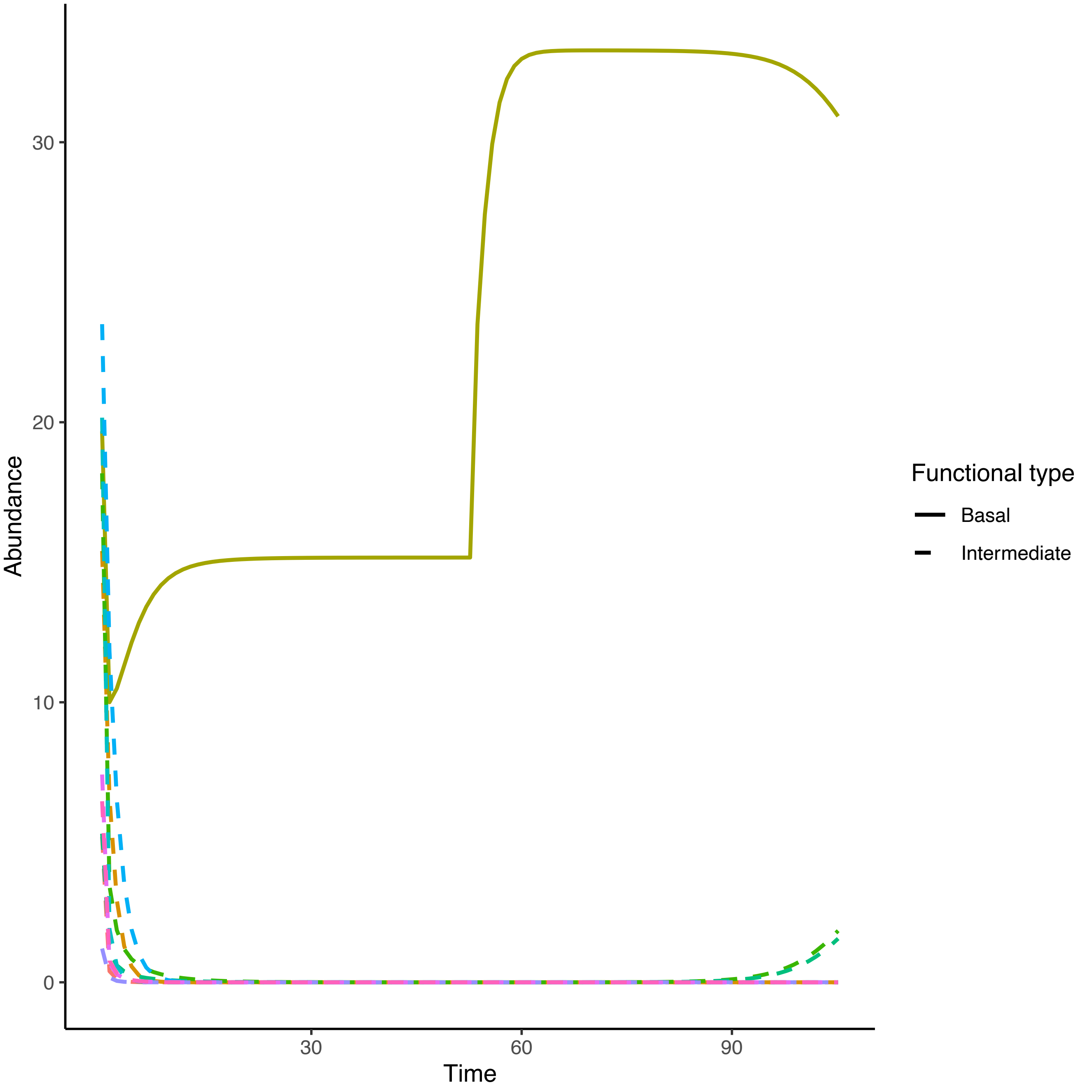
Abundance
30
20
10
0
30
60
90
Time
Functional type
Basal
Intermediate

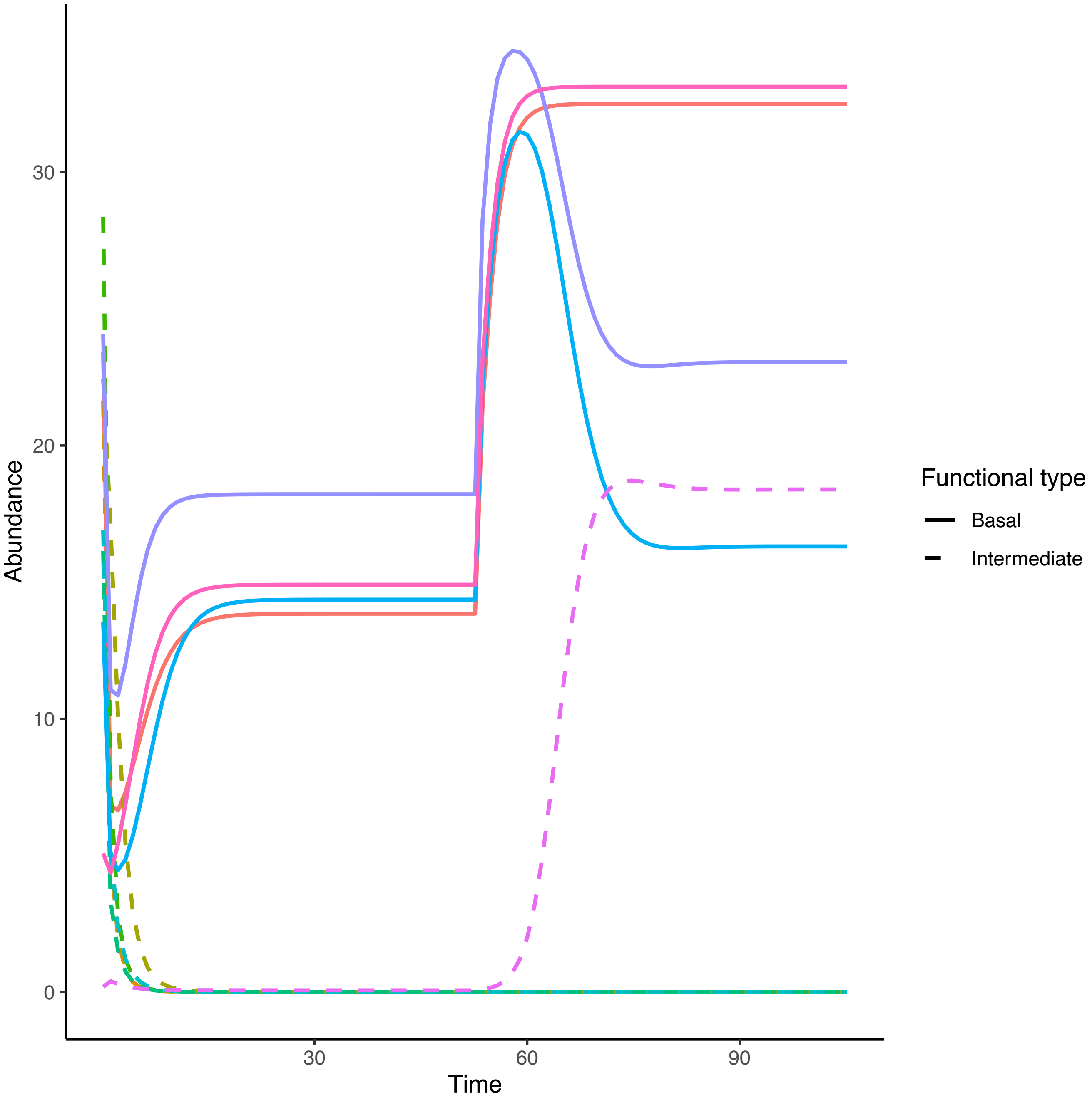
Abundance
30
20
10
0
30
60
90
Time
Functional type
Basal
Intermediate

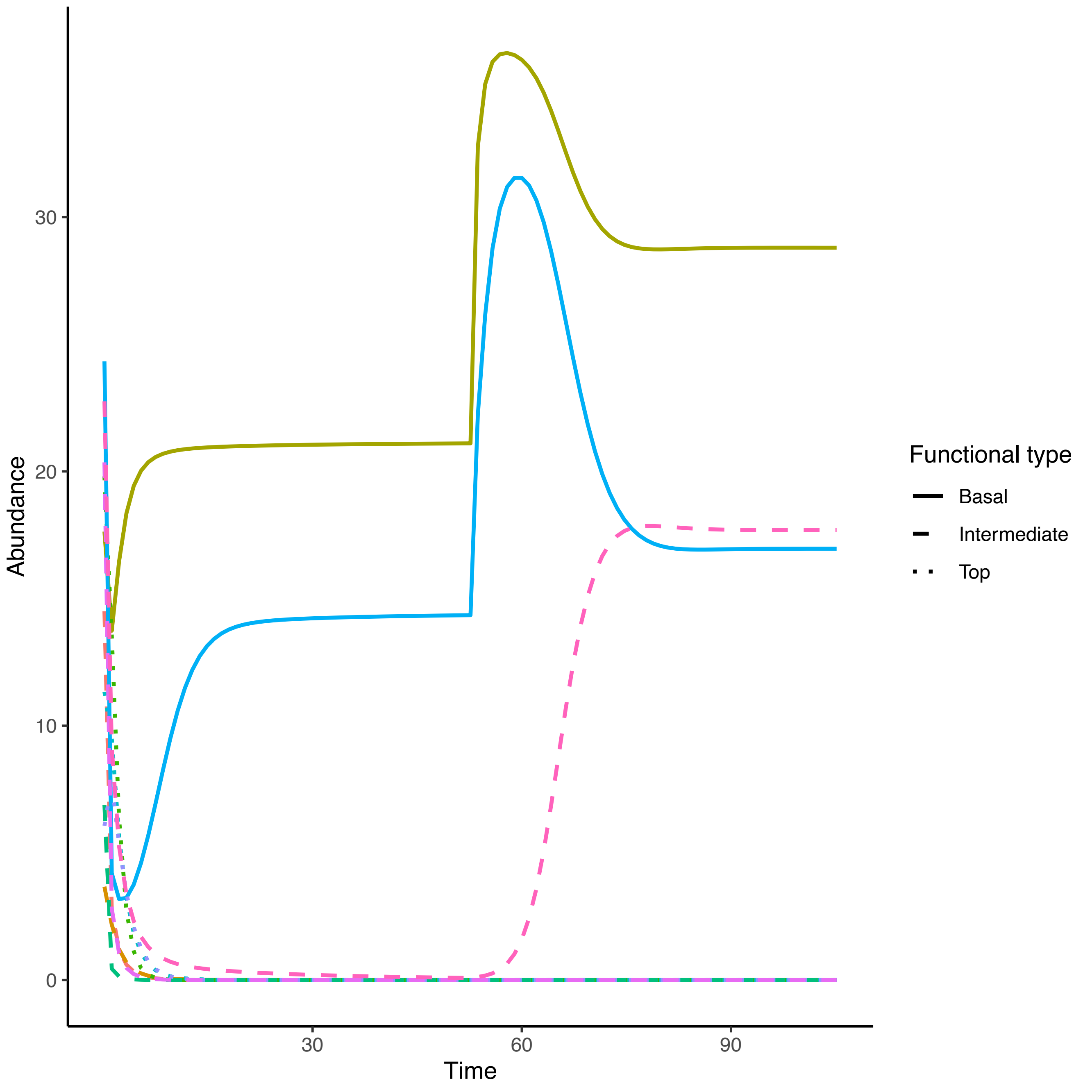

Abundance
Time
0
10
20
30
30
60
90
Functional type
Basal
Intermediate
Top

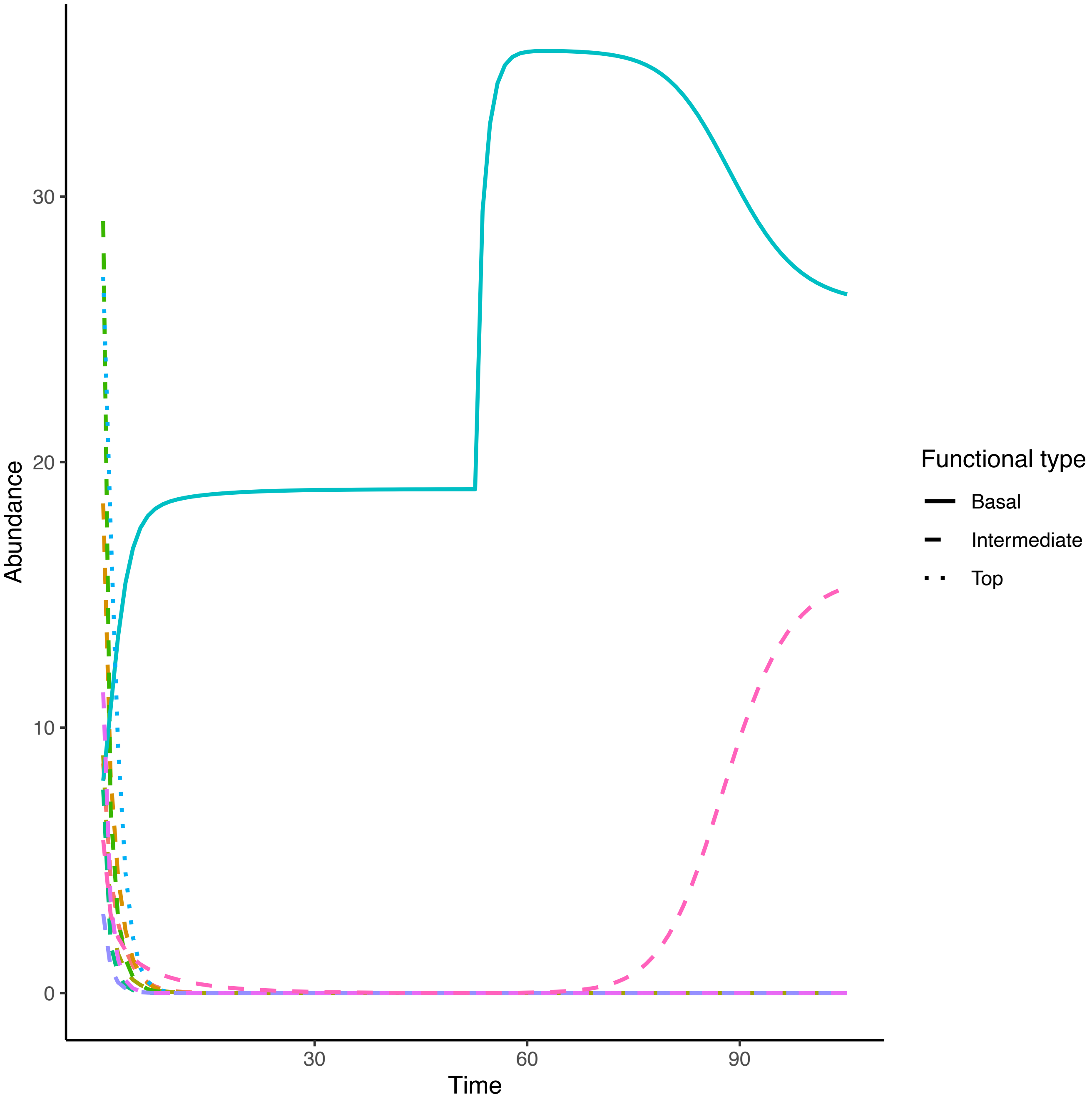

Functional type
Basal
Intermediate
Top
Abundance
Time
0
10
20
30
30
60
90

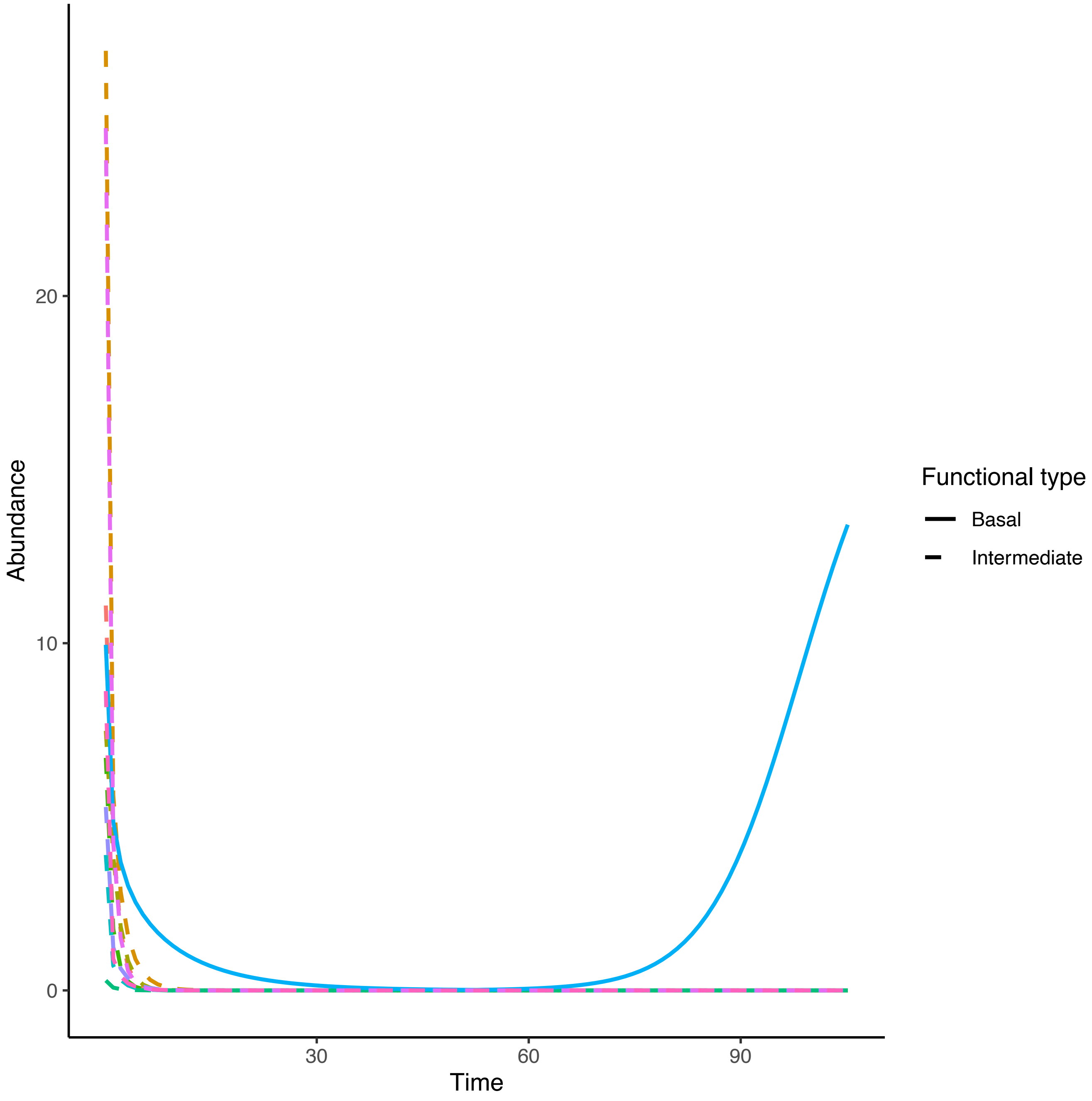

Abundance
20
10
0
30
60
90
Time
Functional type
Basal
Intermediate

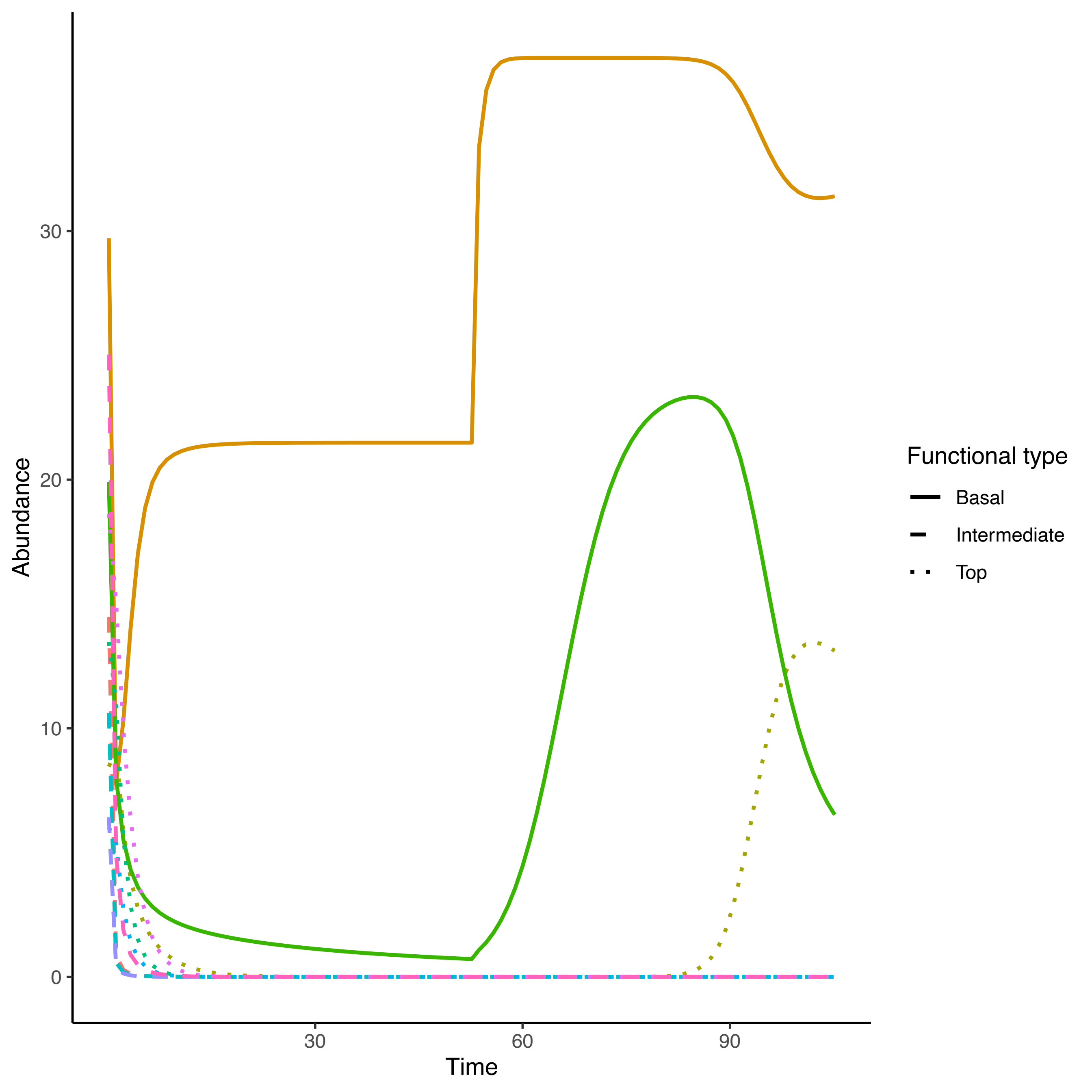

Abundance
30
20
10
0
30
60
90
Time
Functional type
Basal
Intermediate
Top

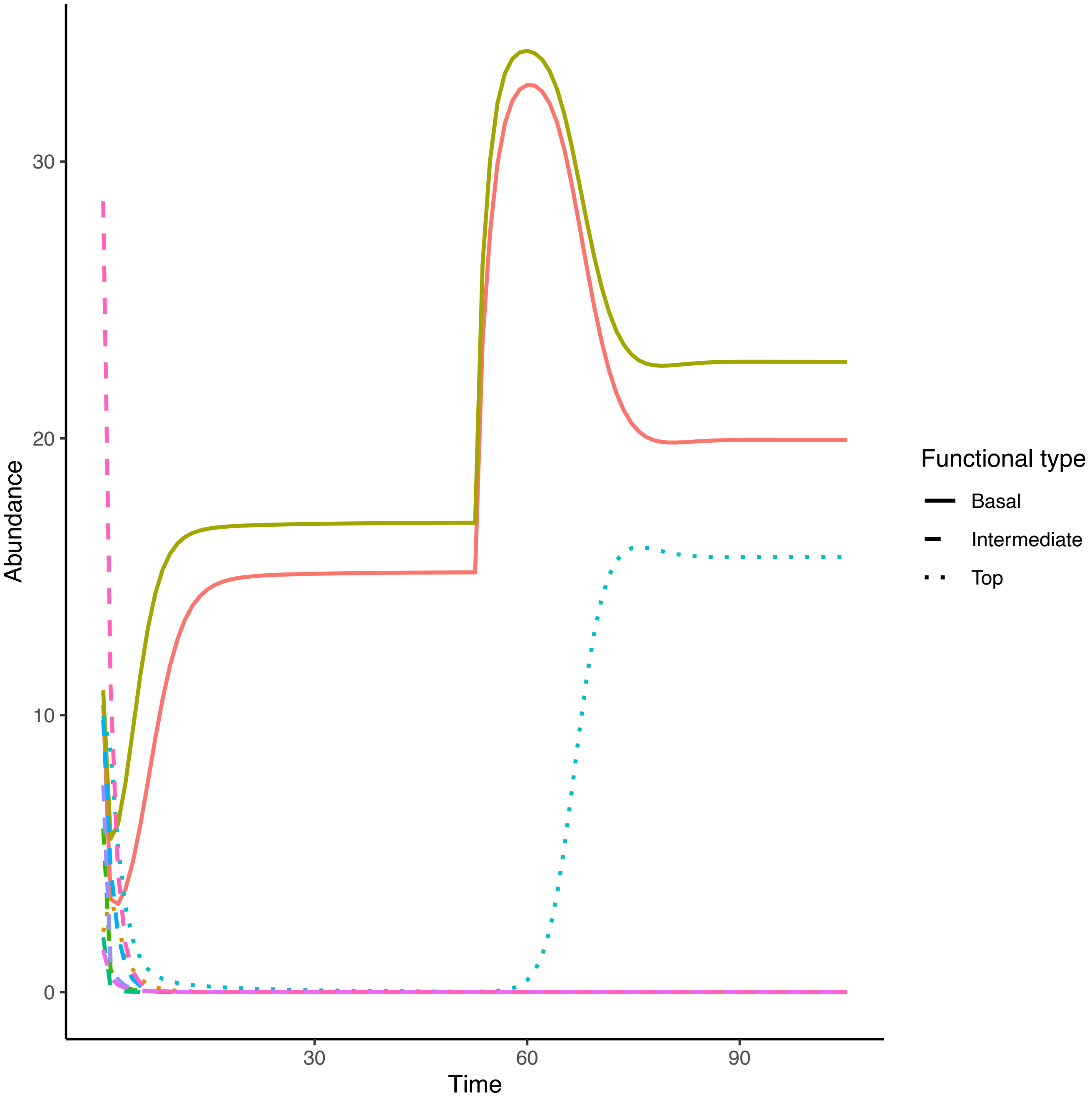

Abundance
Time
0
10
20
30
30
60
90
Functional type
Basal
Intermediate
Top

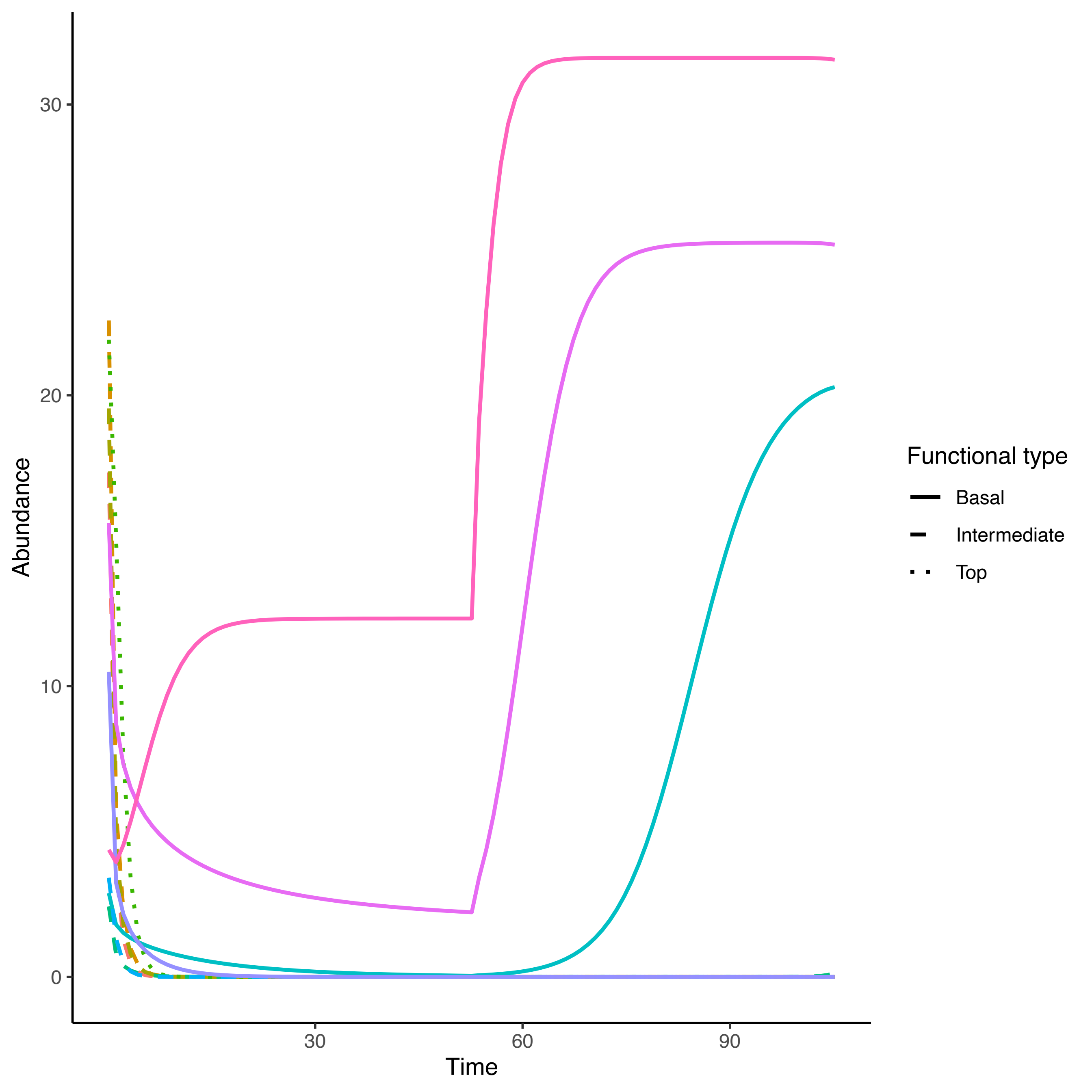

30
20
10
0
Abundance
30
60
90
Time
Functional type
Basal
Intermediate
Top

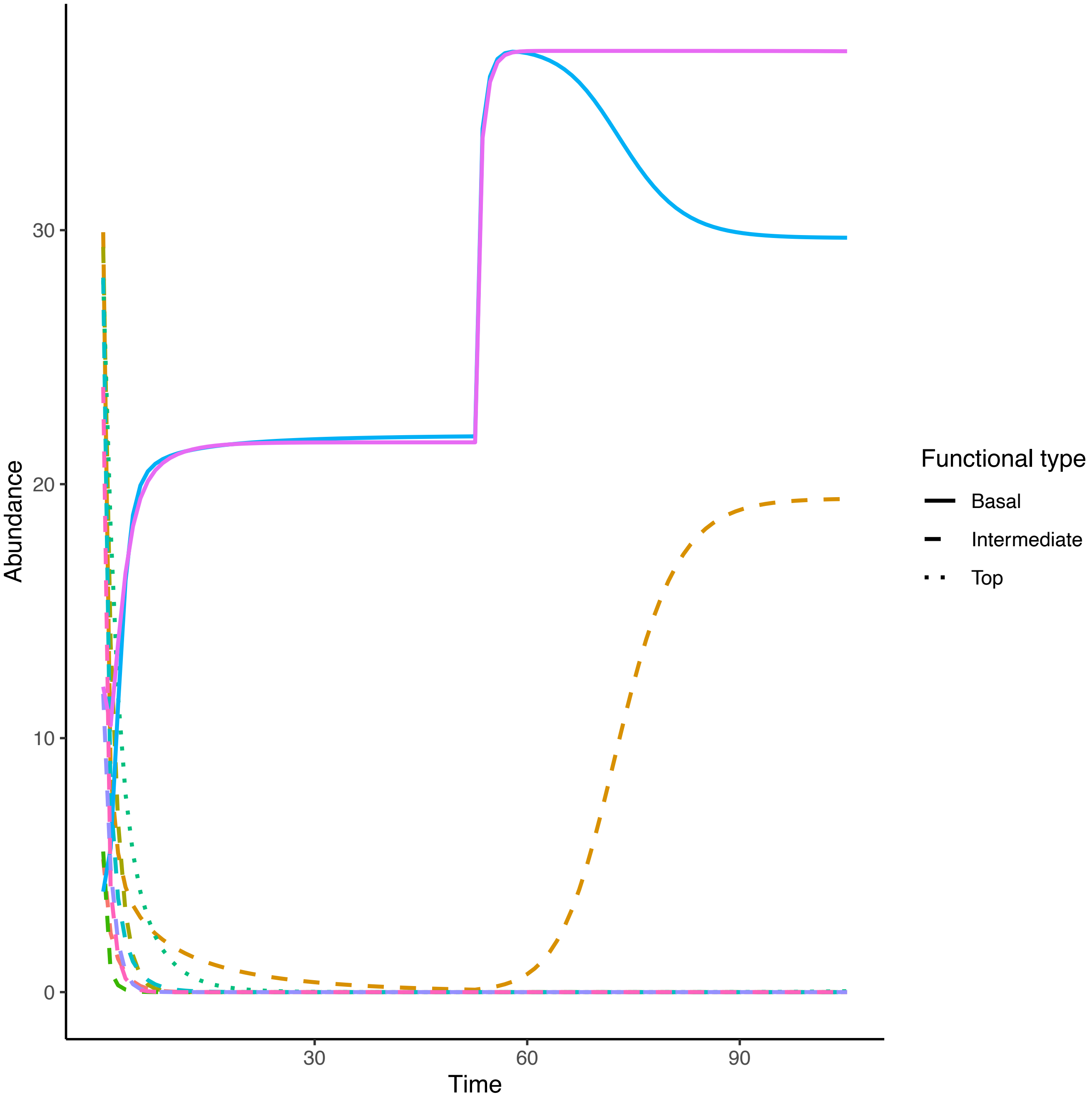

Abundance
0
10
20
30
Time
30
60
90
Functional type
Basal
Intermediate
Top

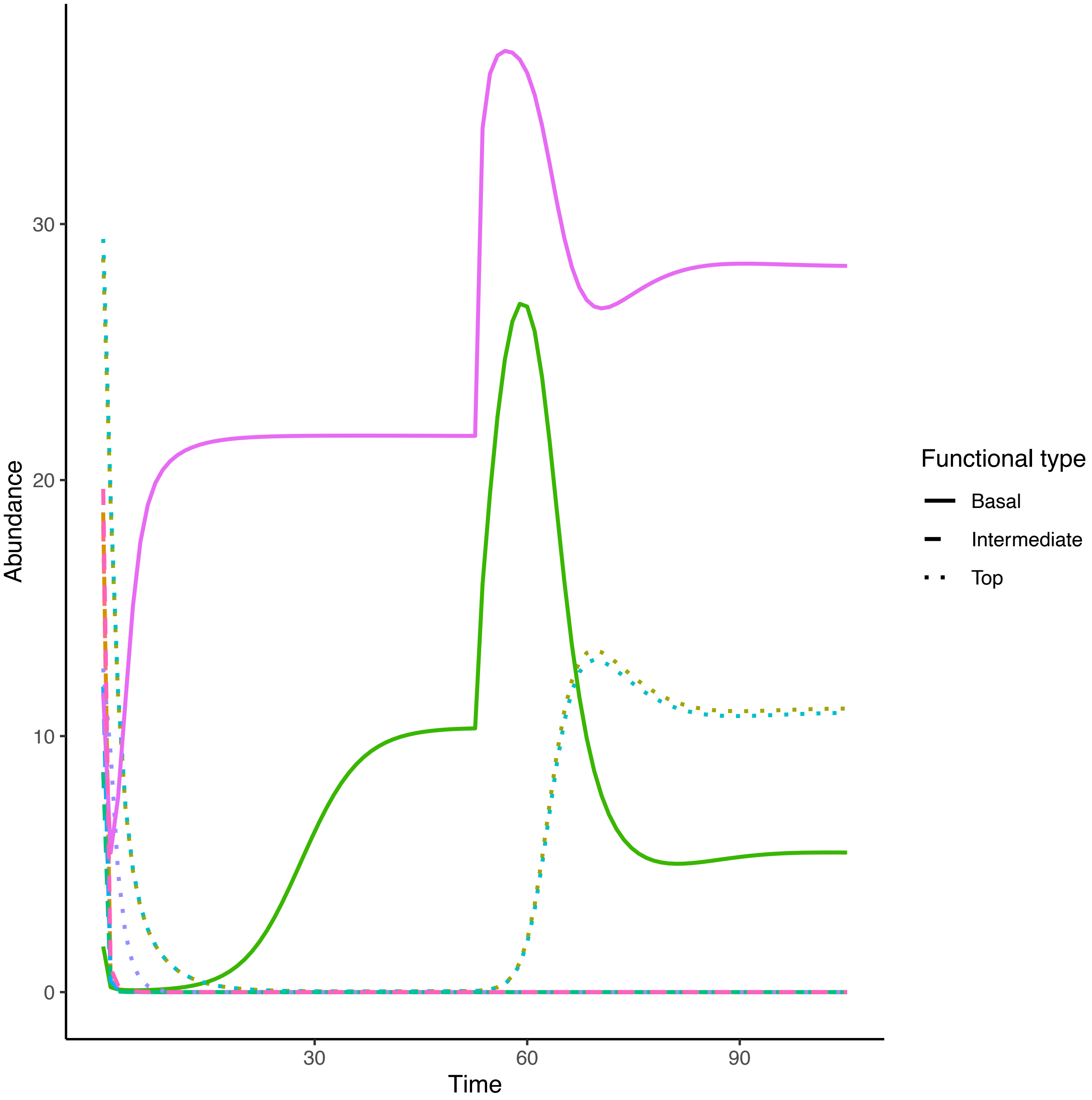

Abundance
30
20
10
0
30
60
90
Time
Functional type
Basal
Intermediate
Top

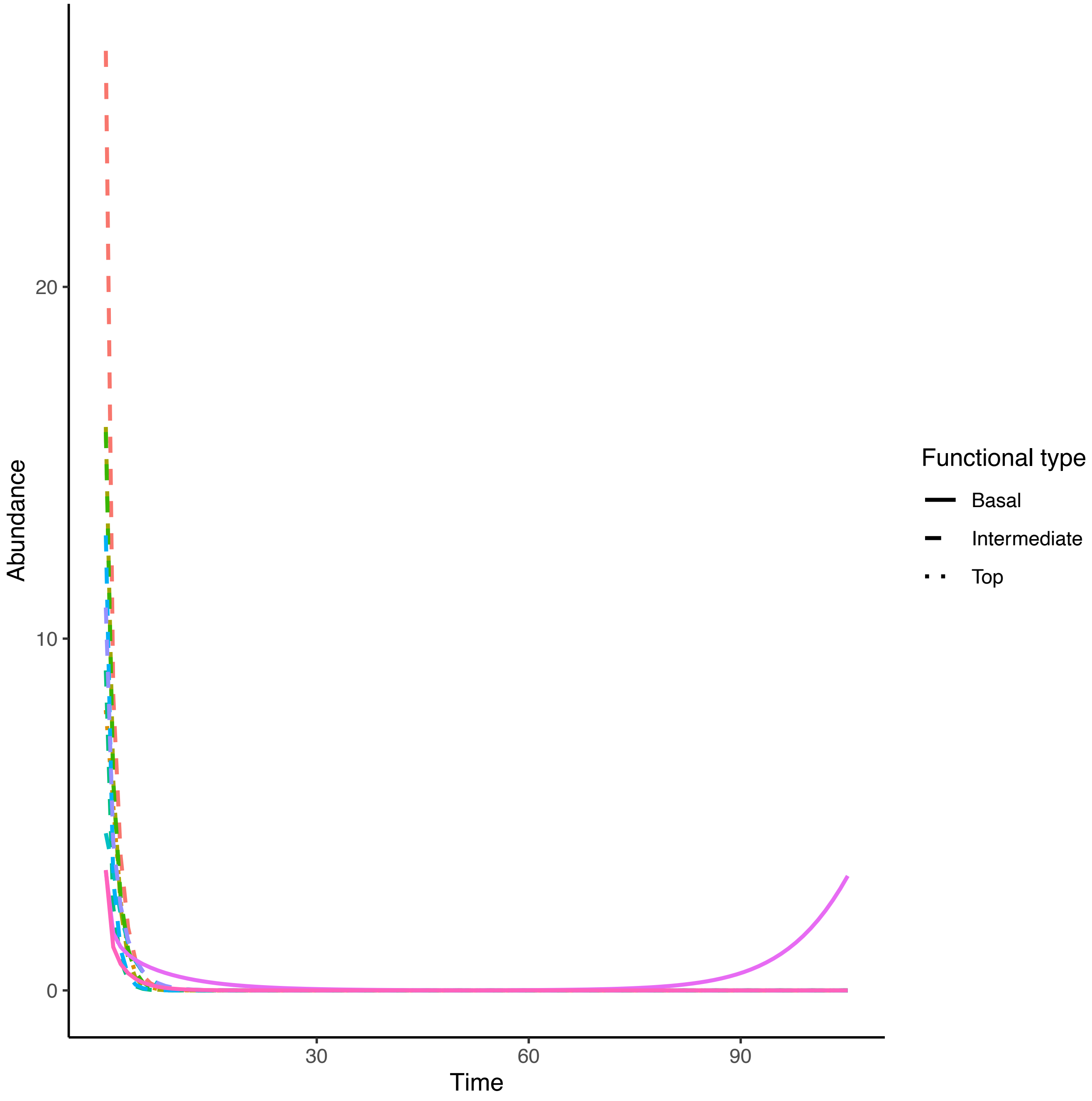

Abundance
20
10
0
30
60
90
Time
Functional type
Basal
Intermediate
Top

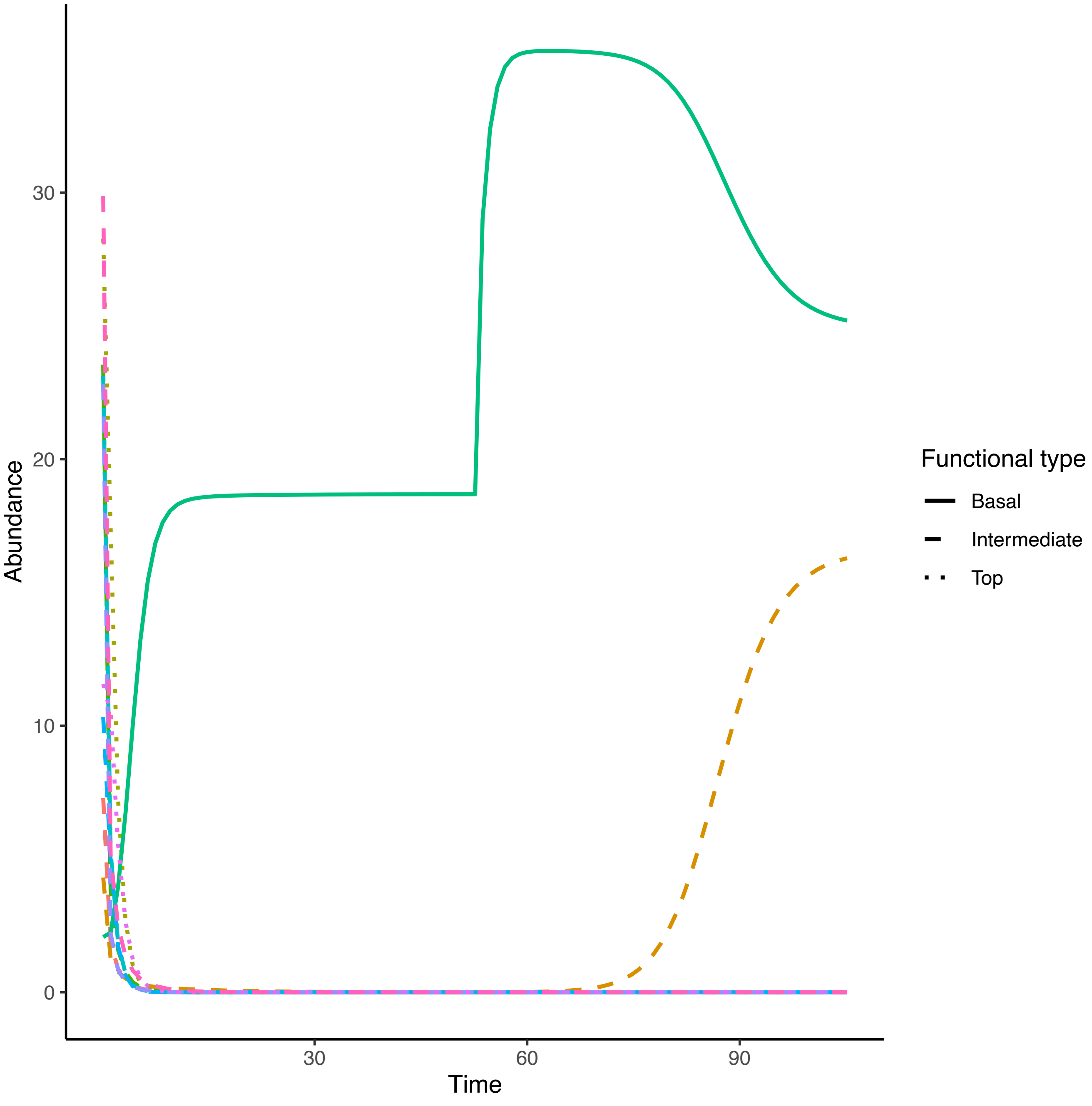

Abundance
Time
0
10
20
30
30
60
90
Functional type
Basal
Intermediate
Top

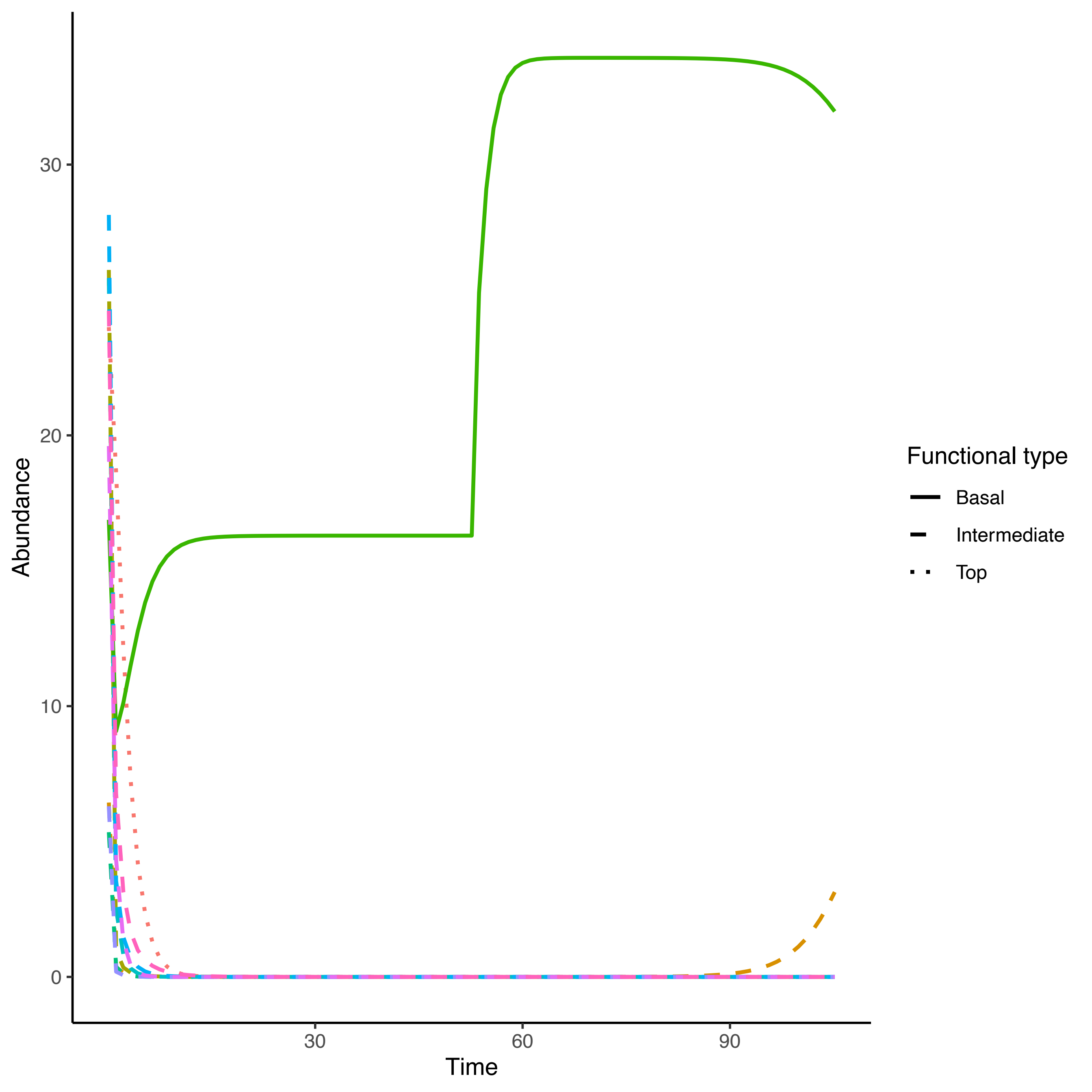

Abundance
30
20
10
0
30
60
90
Time
Functional type
Basal
Intermediate
Top

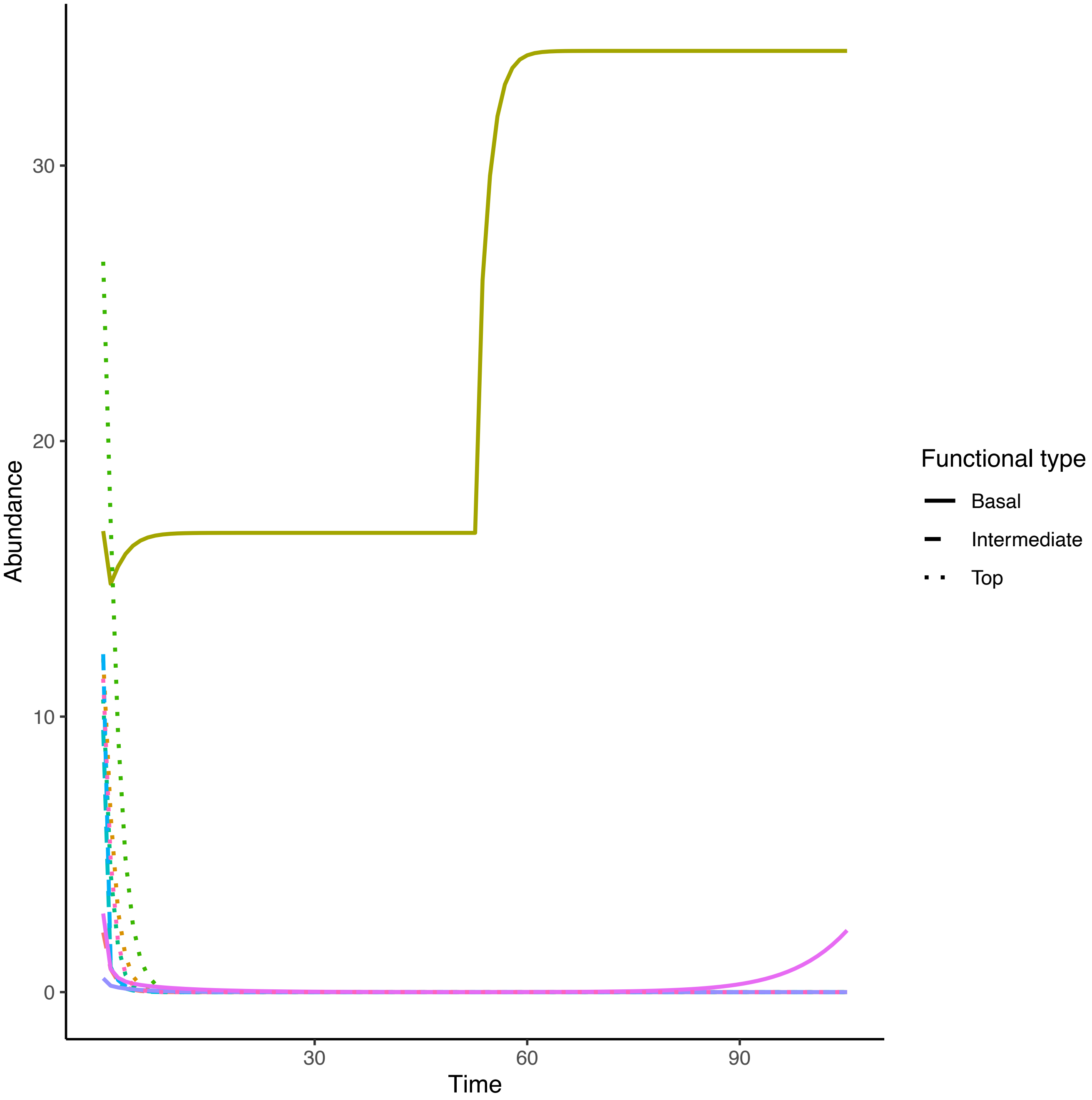

Abundance
Time
0
10
20
30
30
60
90
Functional type
Basal
Intermediate
Top

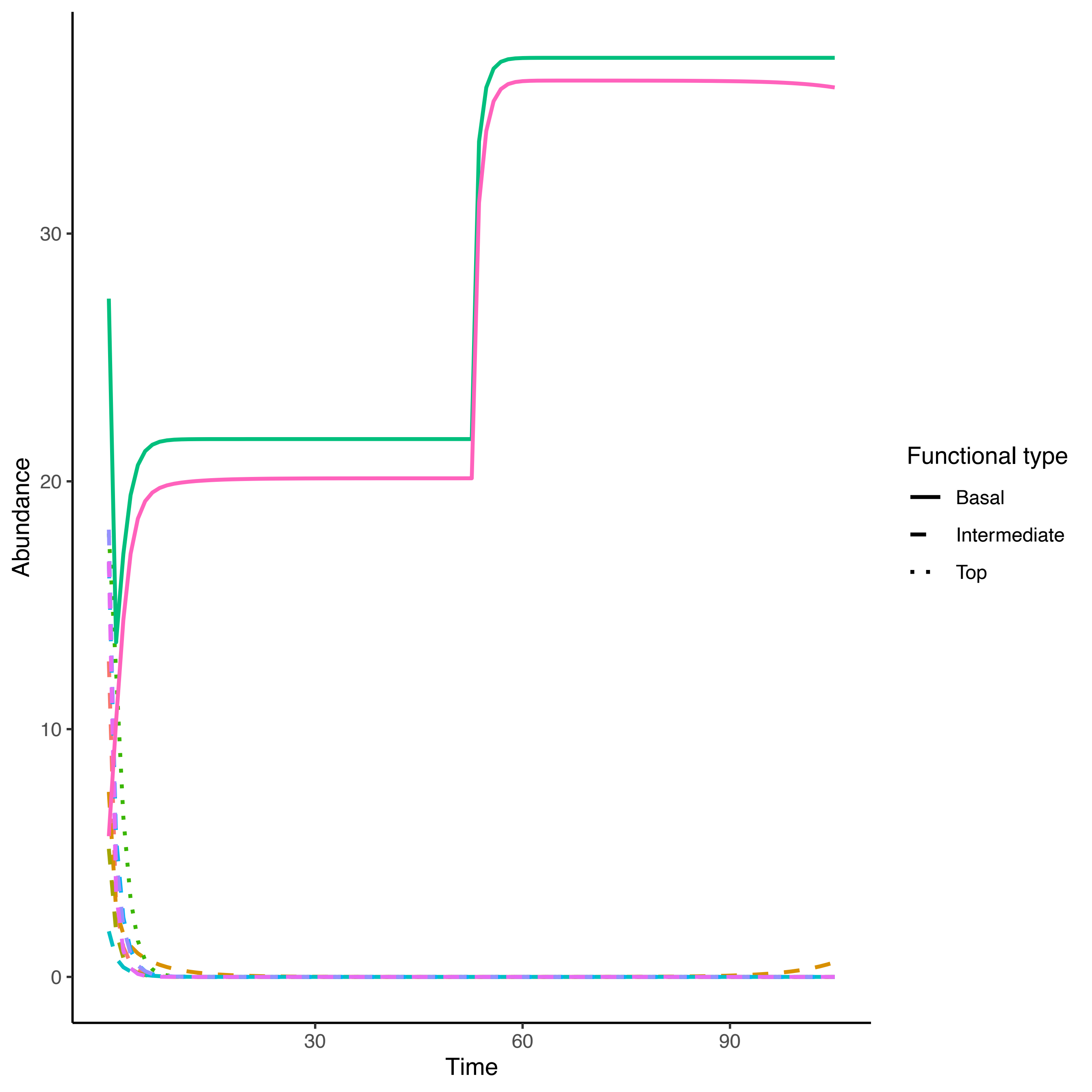

Abundance
30
20
10
0
Time
30
60
90
Functional type
Basal
Intermediate
Top

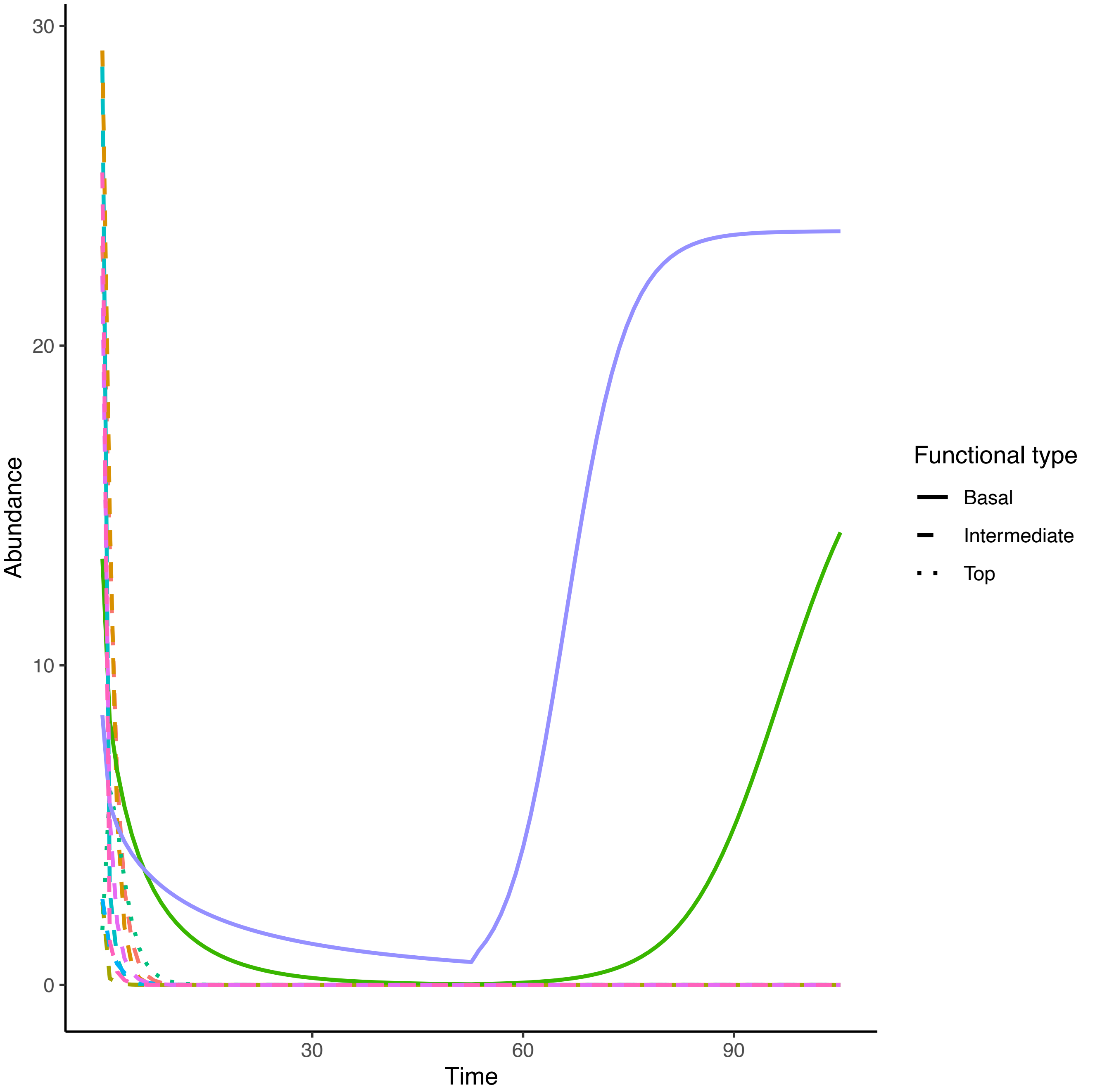

Abundance
30
20
10
0
30
60
90
Time
Functional type
Basal
Intermediate
Top

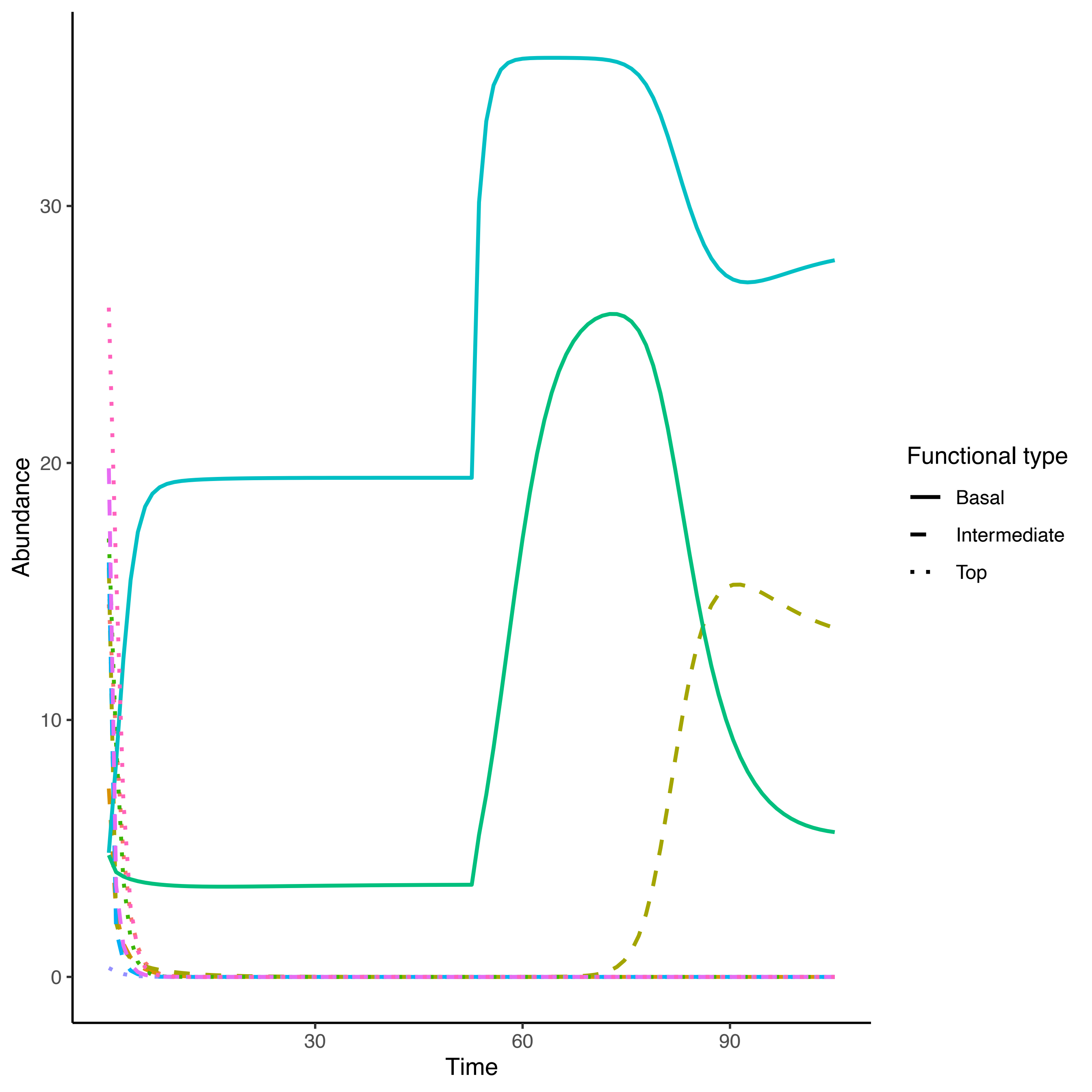

Abundance
Time
0
10
20
30
30
60
90
Functional type
Basal
Intermediate
Top

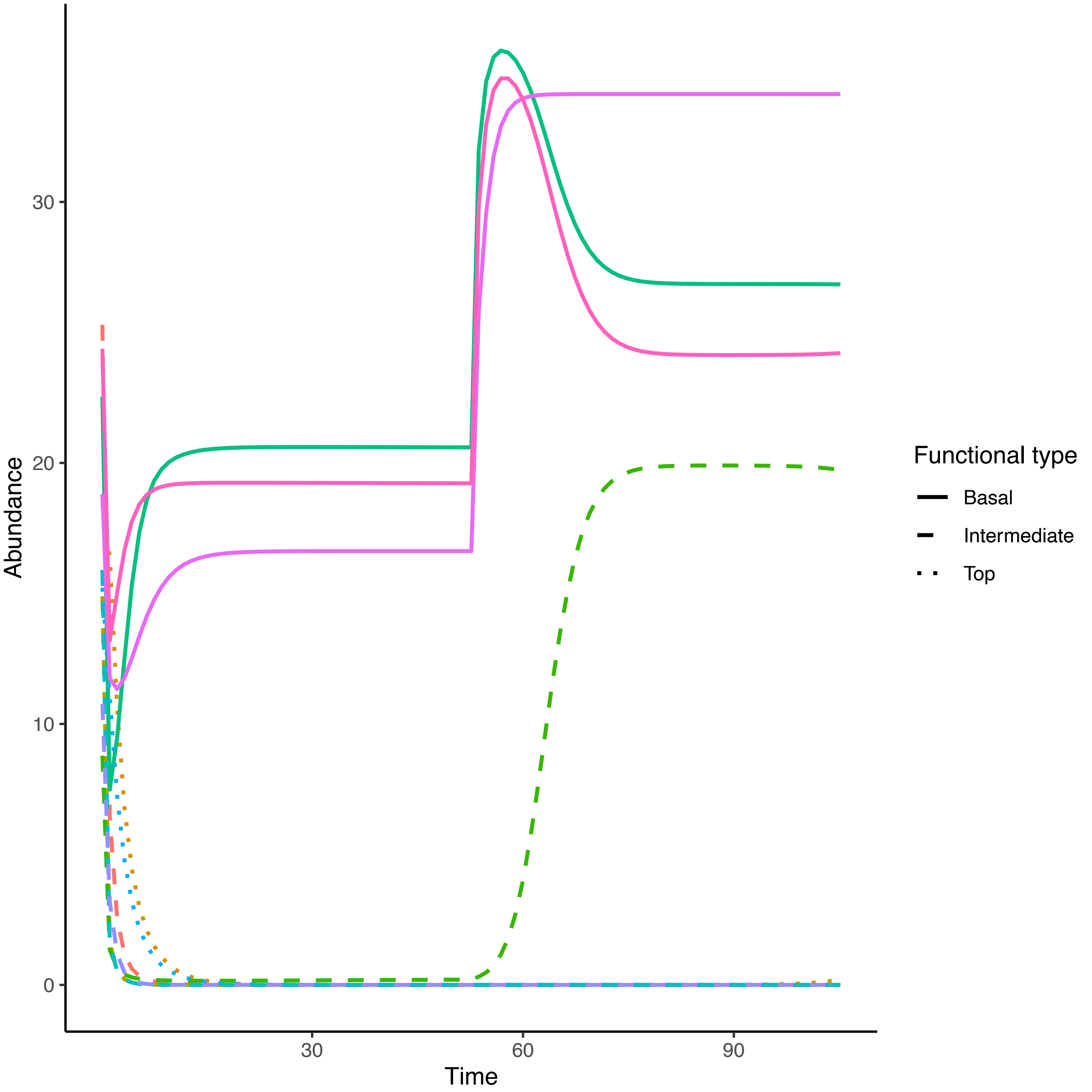
Abundance
Time
0
10
20
30
30
60
90
Functional type
Basal
Intermediate
Top

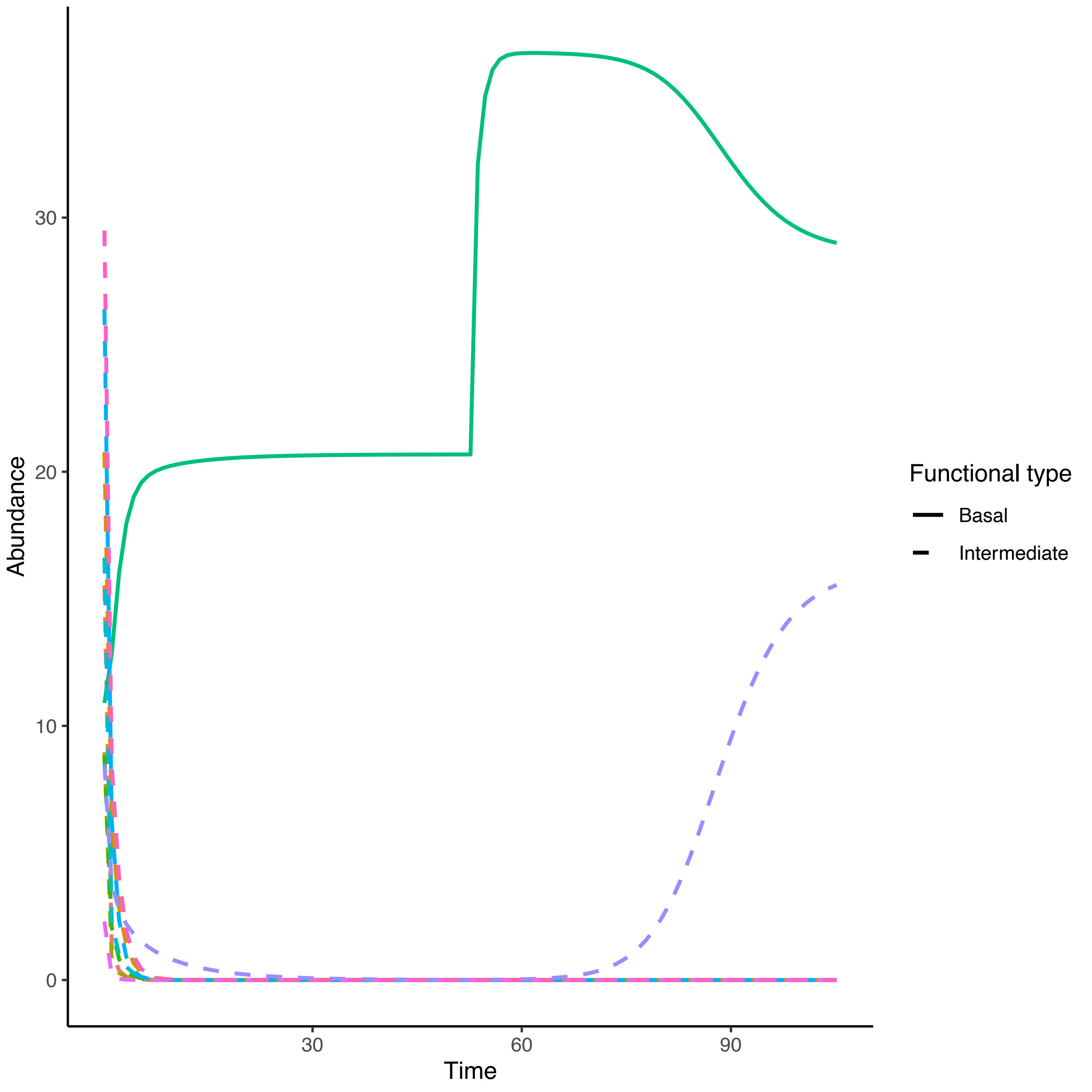

Abundance
Time
0
10
20
30
30
60
90
Functional type
Basal
Intermediate

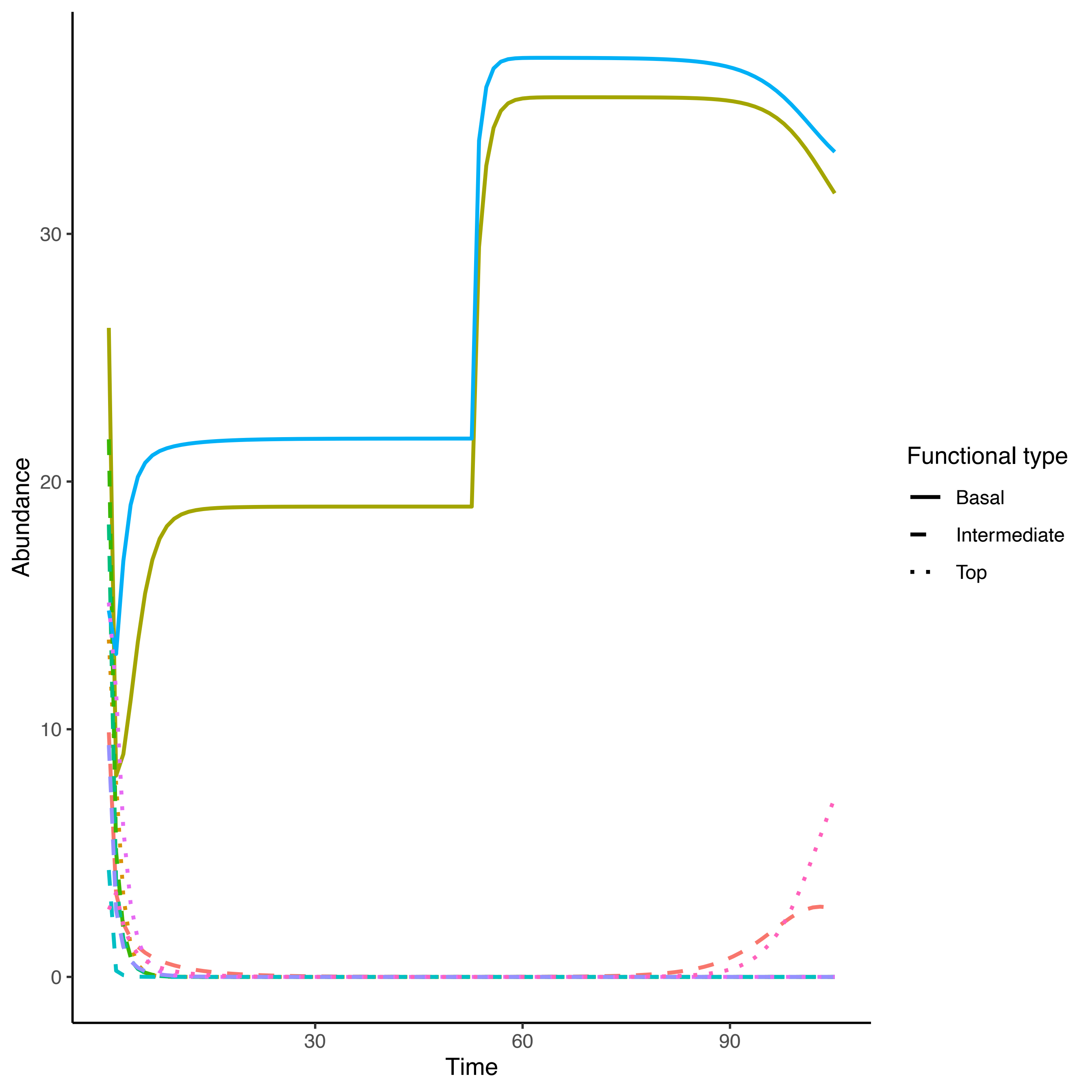
Abundance
30
20
10
0
30
60
90
Time
Functional type
Basal
Intermediate
Top

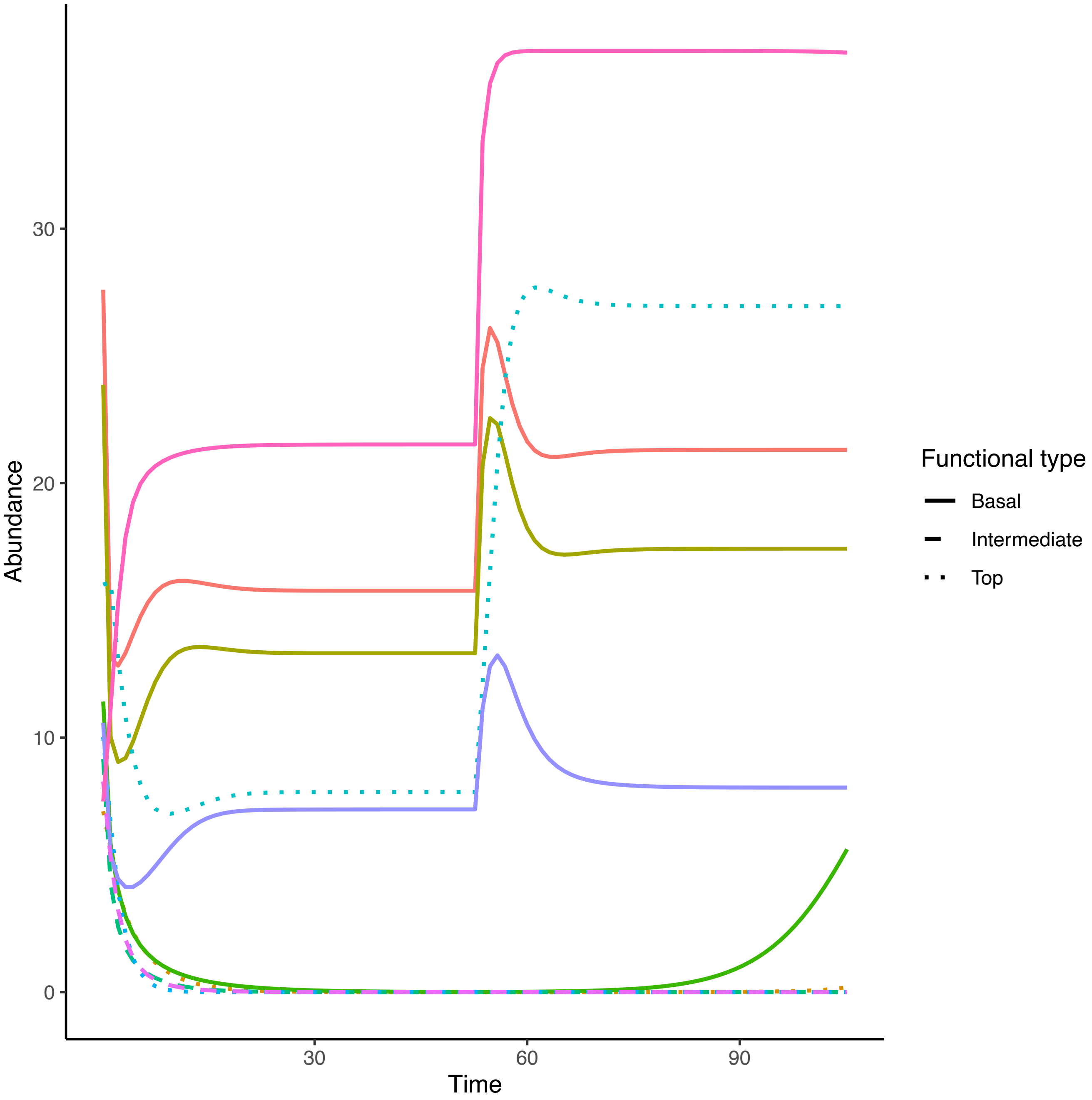

Abundance
30
20
10
0
30
60
90
Time
Functional type
Basal
Intermediate
Top

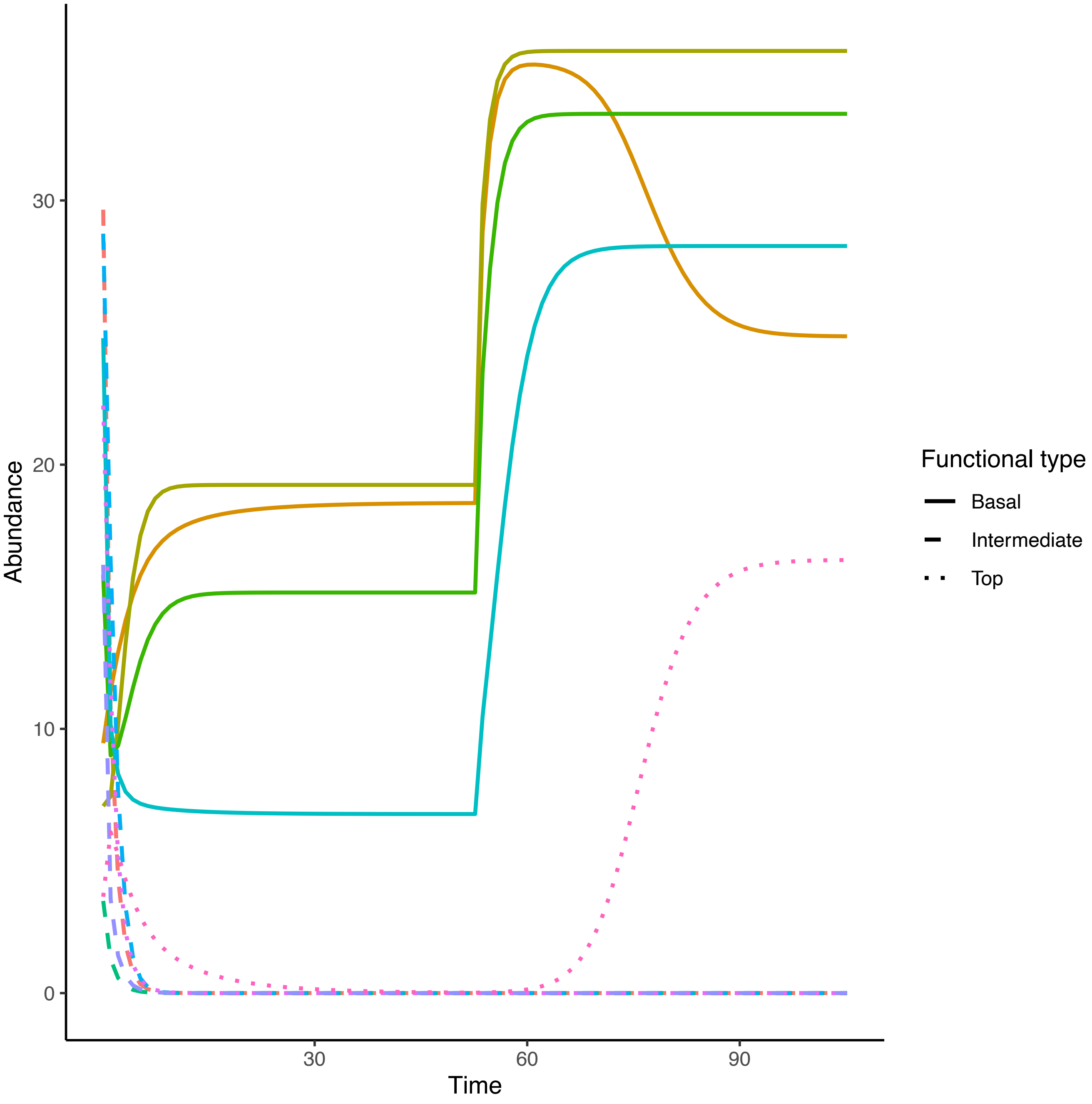

Abundance
30
20
10
0
30
60
90
Time
Functional type
Basal
Intermediate
Top

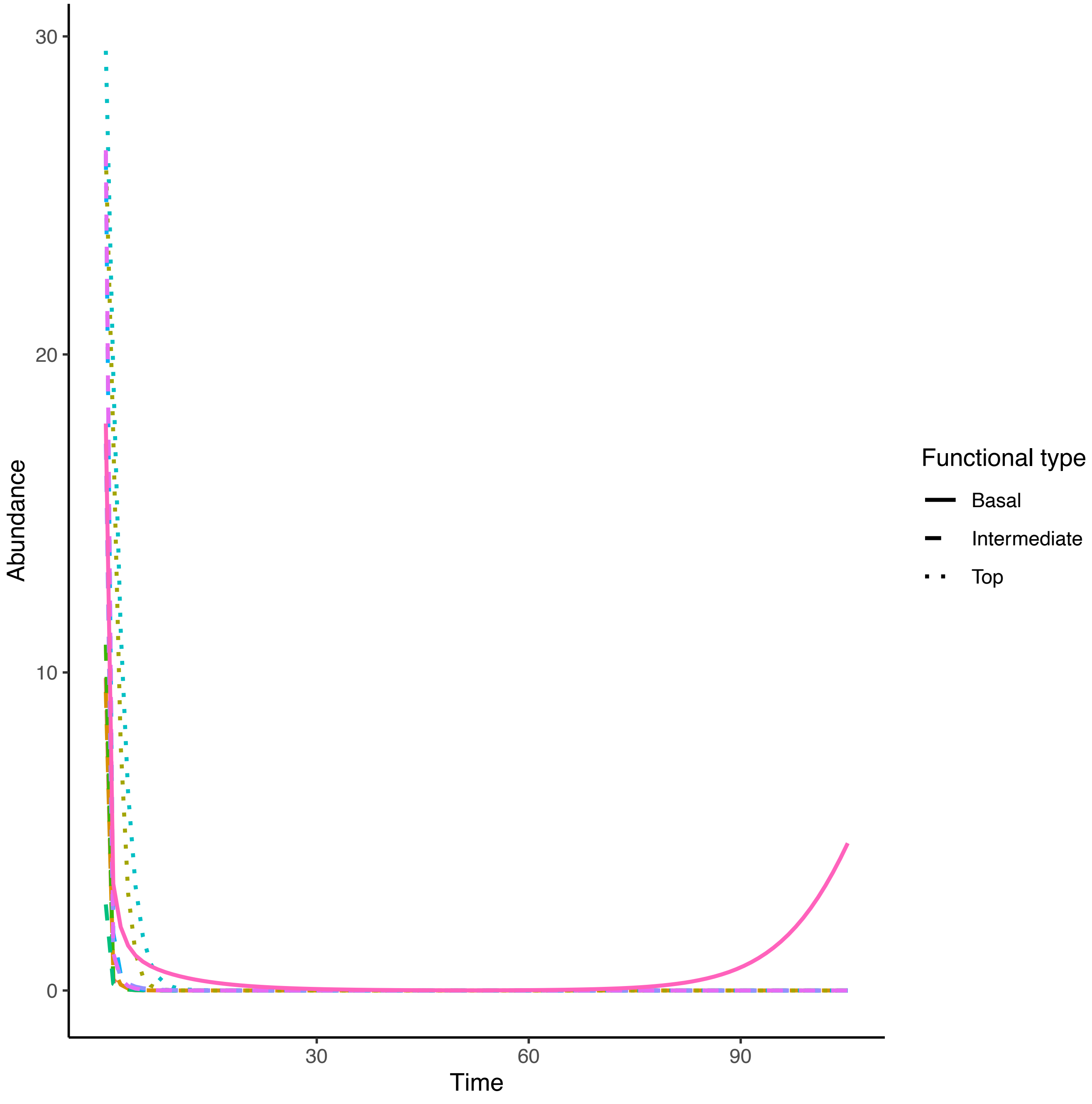

30
20
10
0
Abundance
30
60
90
Time
Functional type
Basal
Intermediate
Top

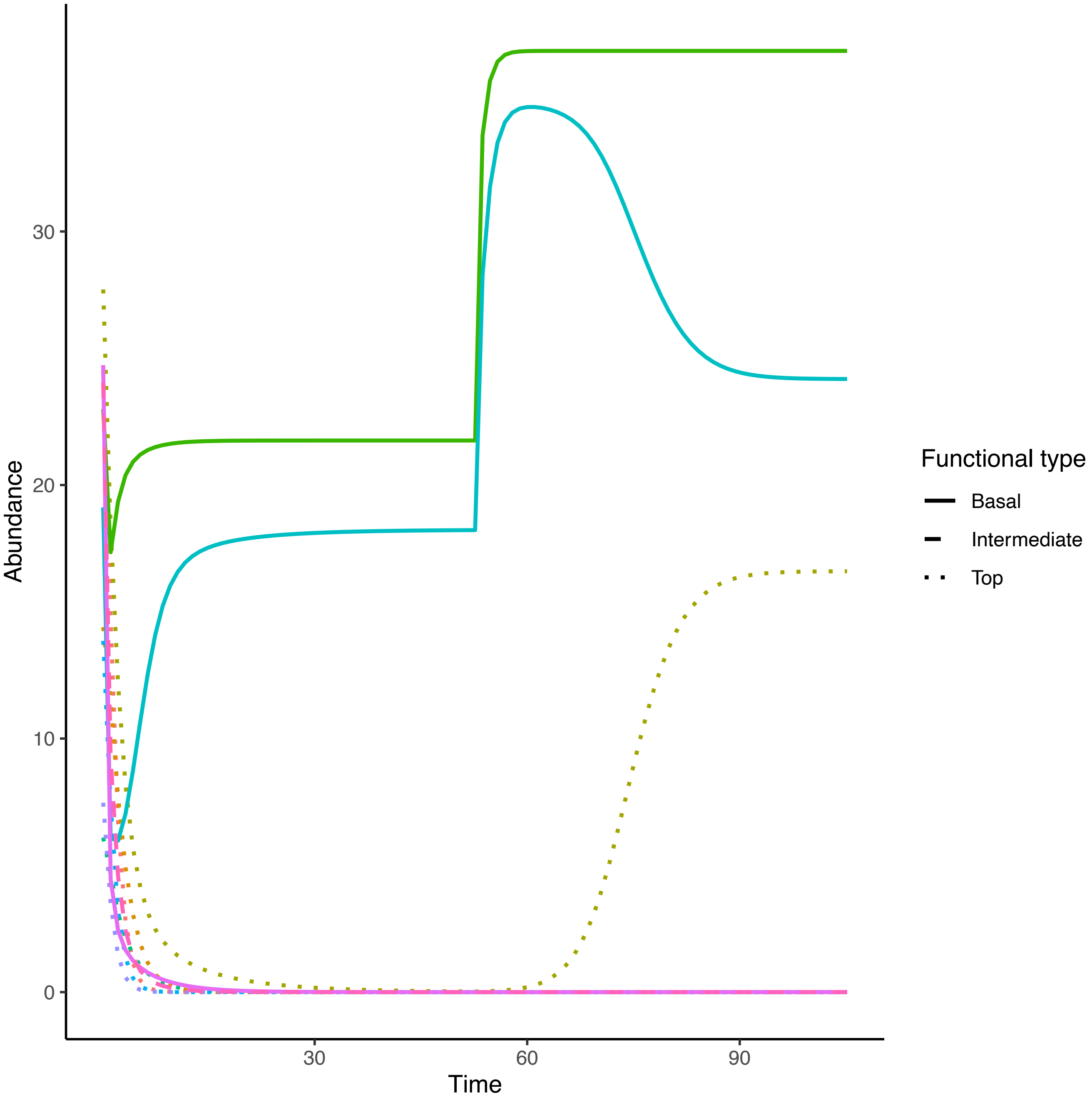
Abundance
0
10
20
30
Time
30
60
90
Functional type
Basal
Intermediate
Top

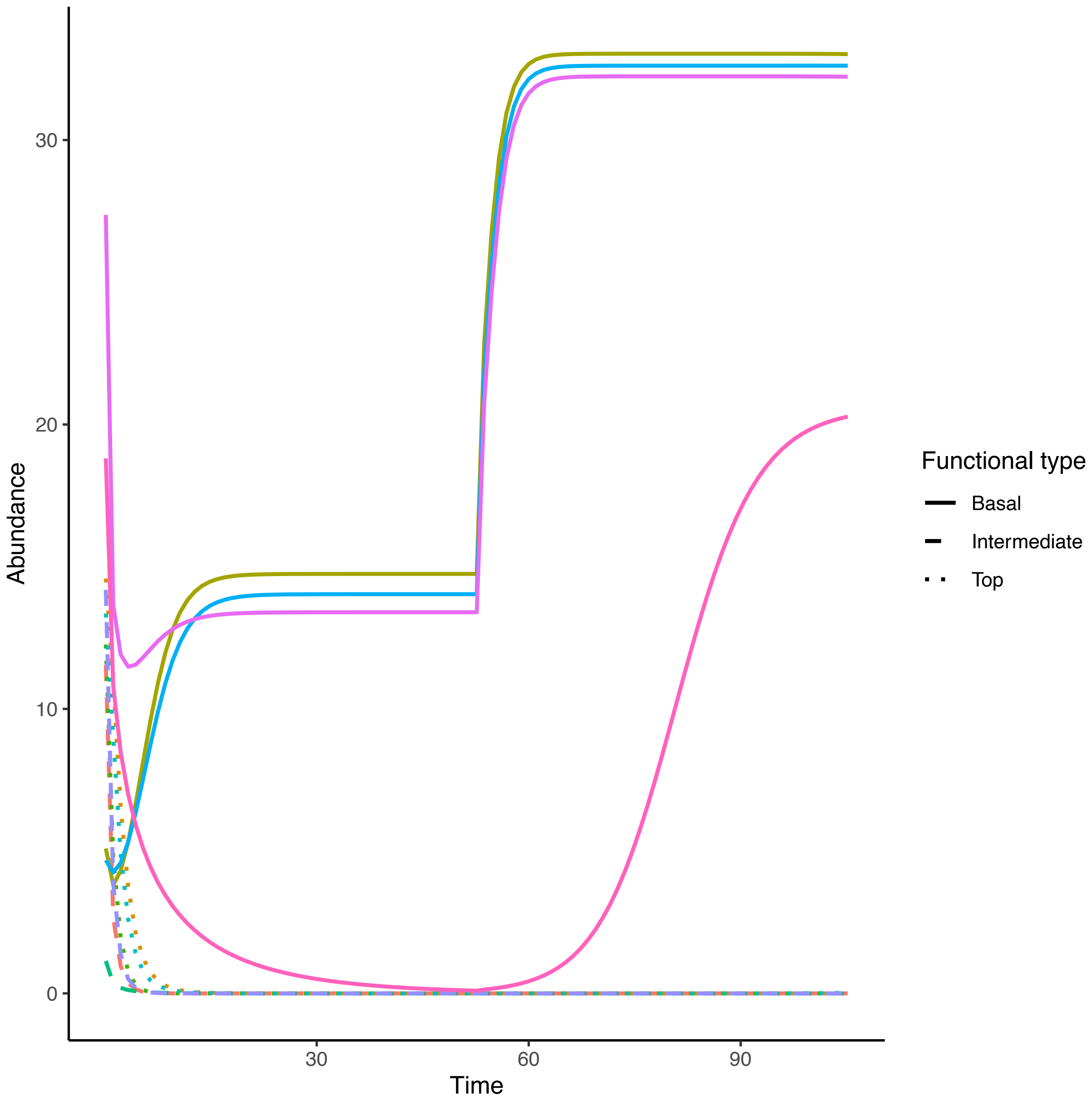

Abundance
0
10
20
30
Time
30
60
90
Functional type
Basal
Intermediate
Top

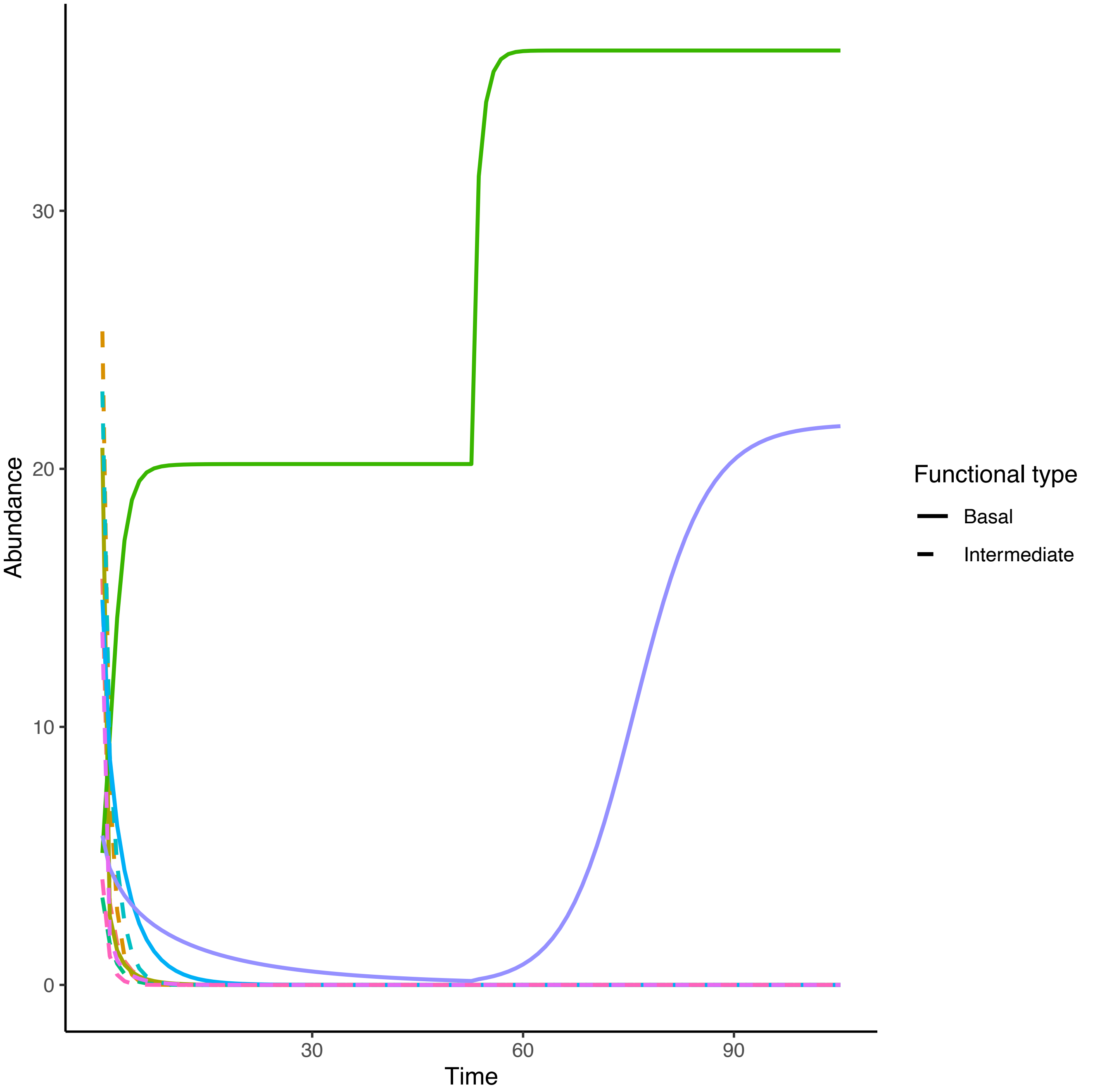

Abundance
30
20
10
0
30
60
90
Time
Functional type
Basal
Intermediate

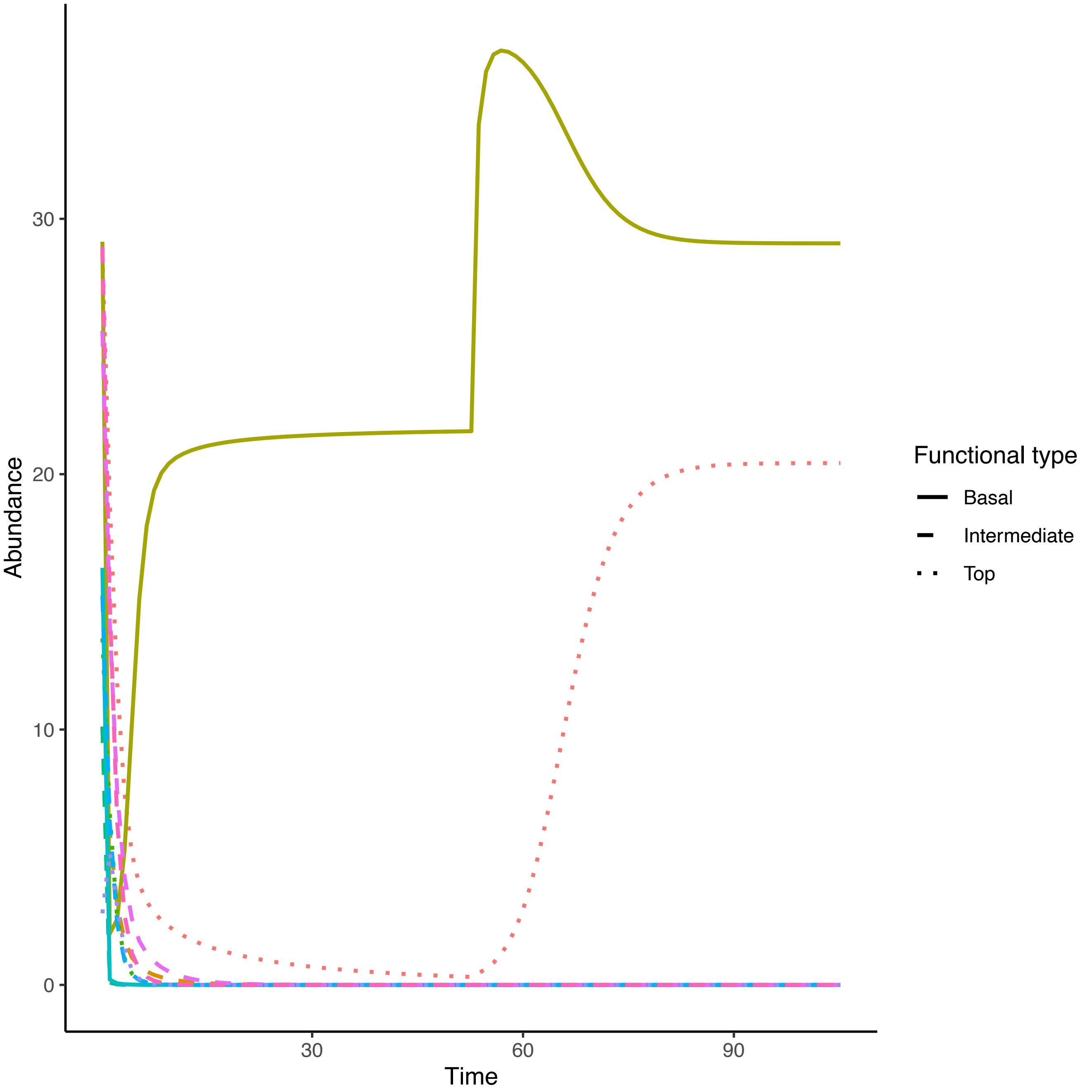

Abundance
0
10
20
30
Time
30
60
90
Functional type
Basal
Intermediate
Top

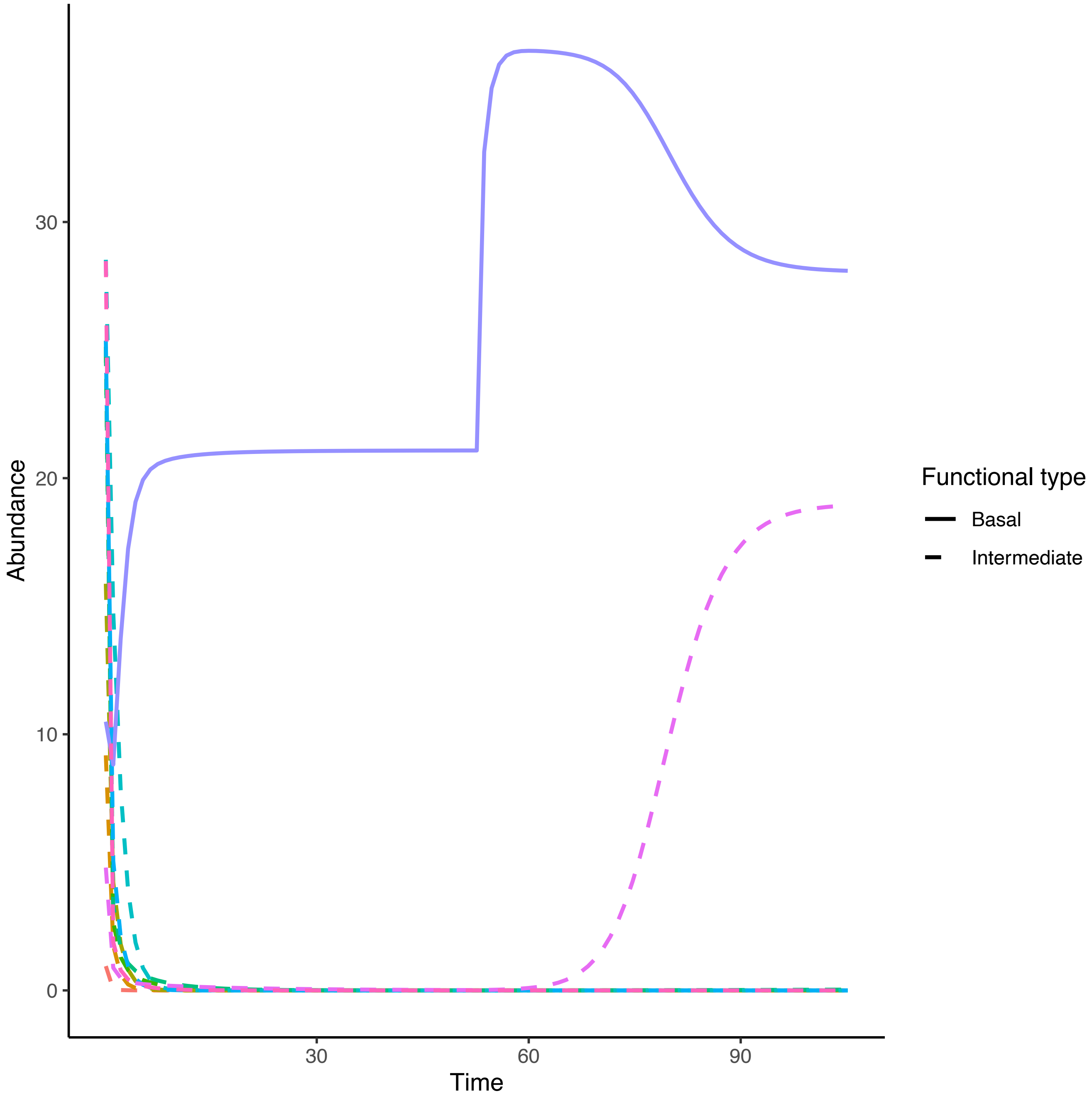

Abundance
30
20
10
0
Time
30
60
90
Functional type
Basal
Intermediate

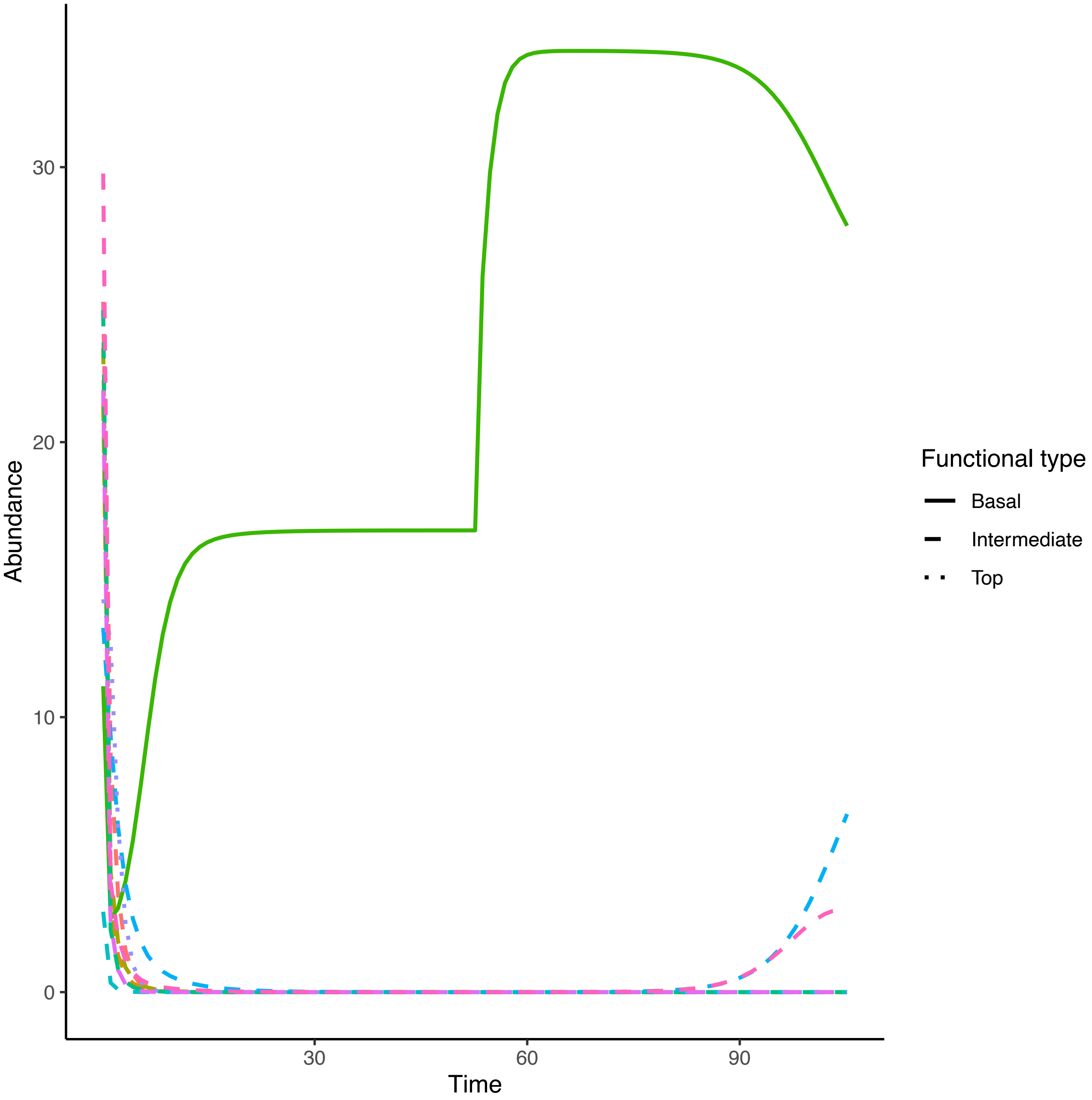

Abundance
30
20
10
0
30
60
90
Time
Functional type
Basal
Intermediate
Top

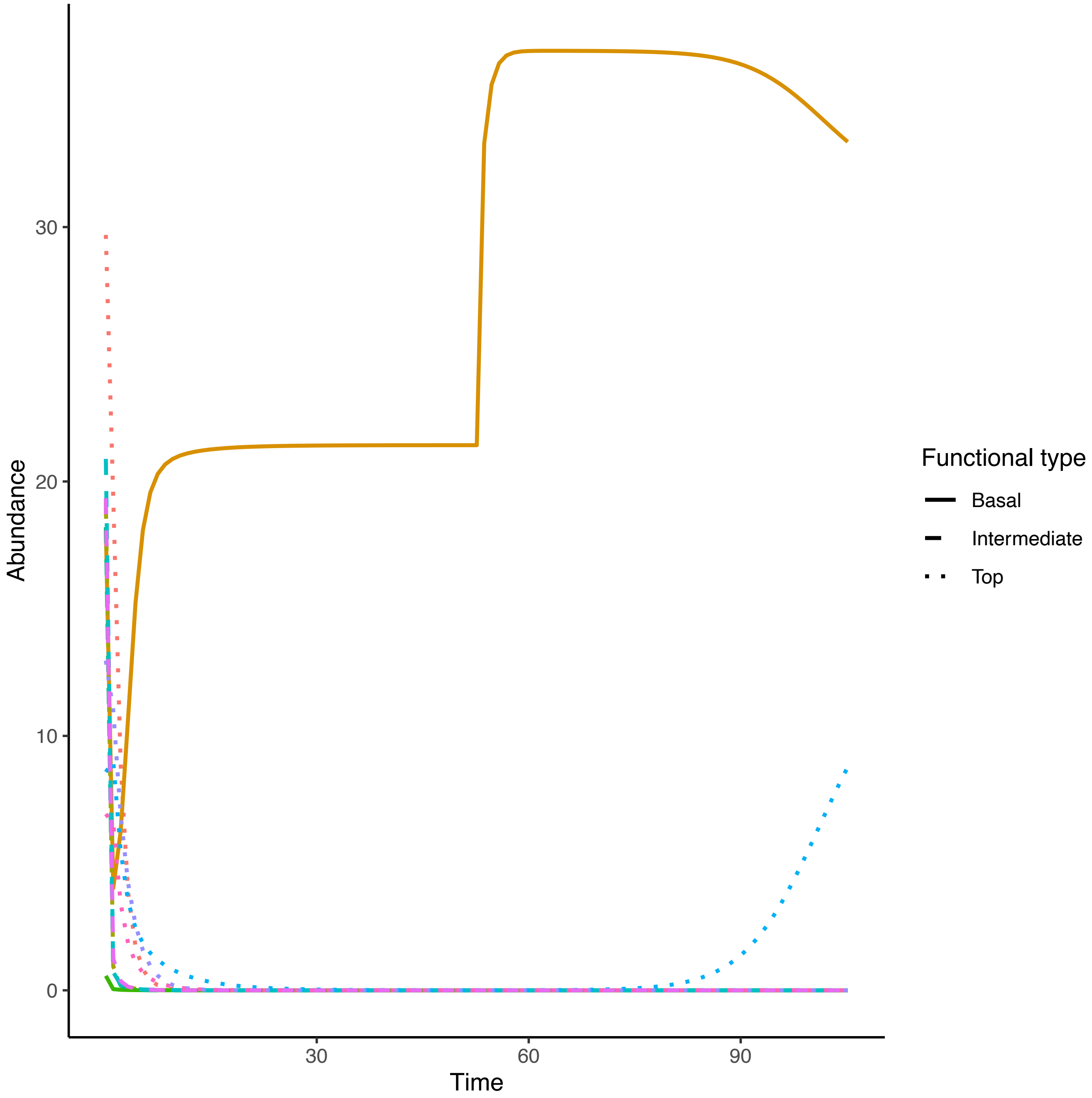

Abundance
30
20
10
0
30
60
90
Time
Functional type
Basal
Intermediate
Top

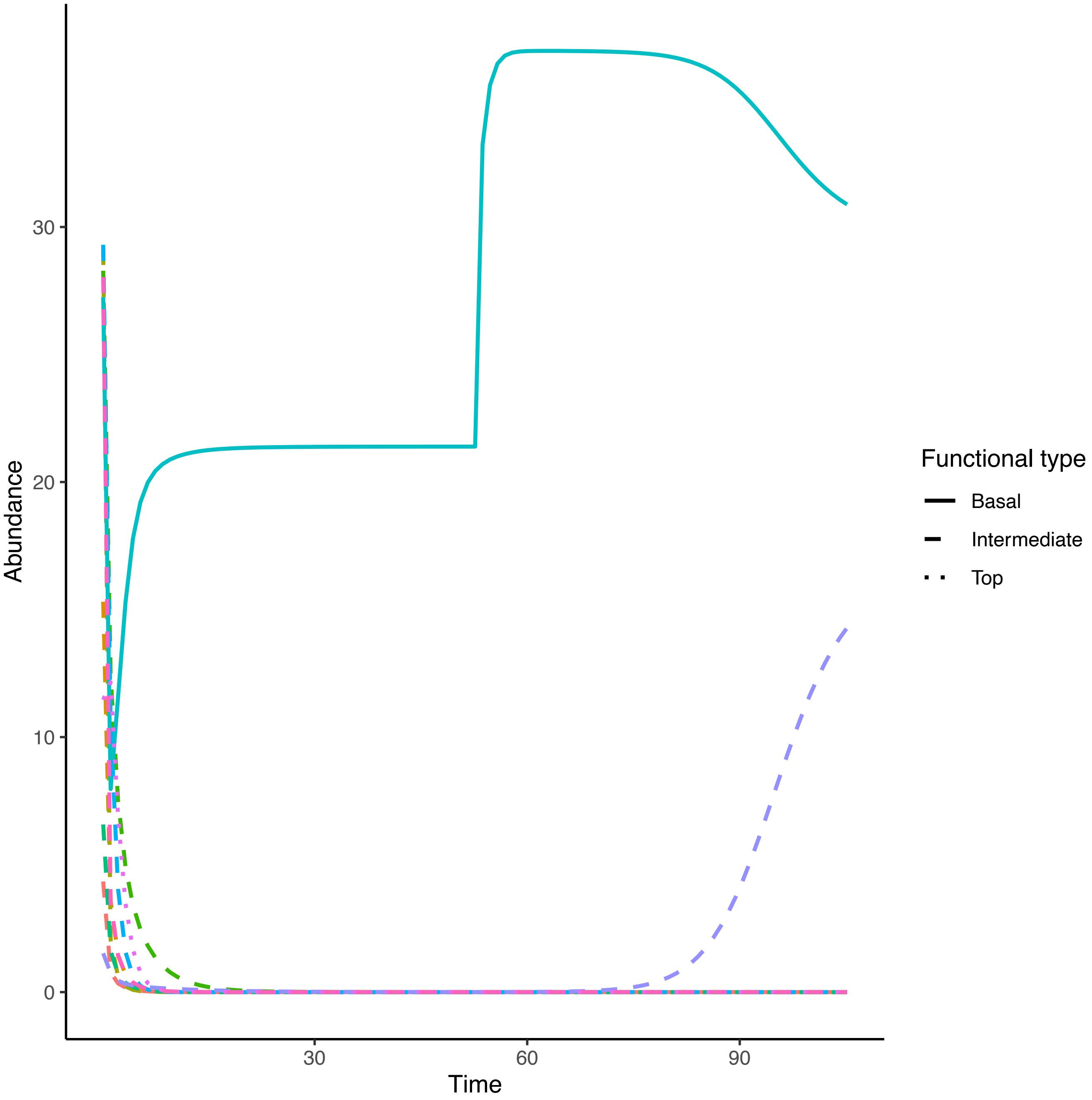

Abundance
Time
0
10
20
30
30
60
90
Functional type
Basal
Intermediate
Top

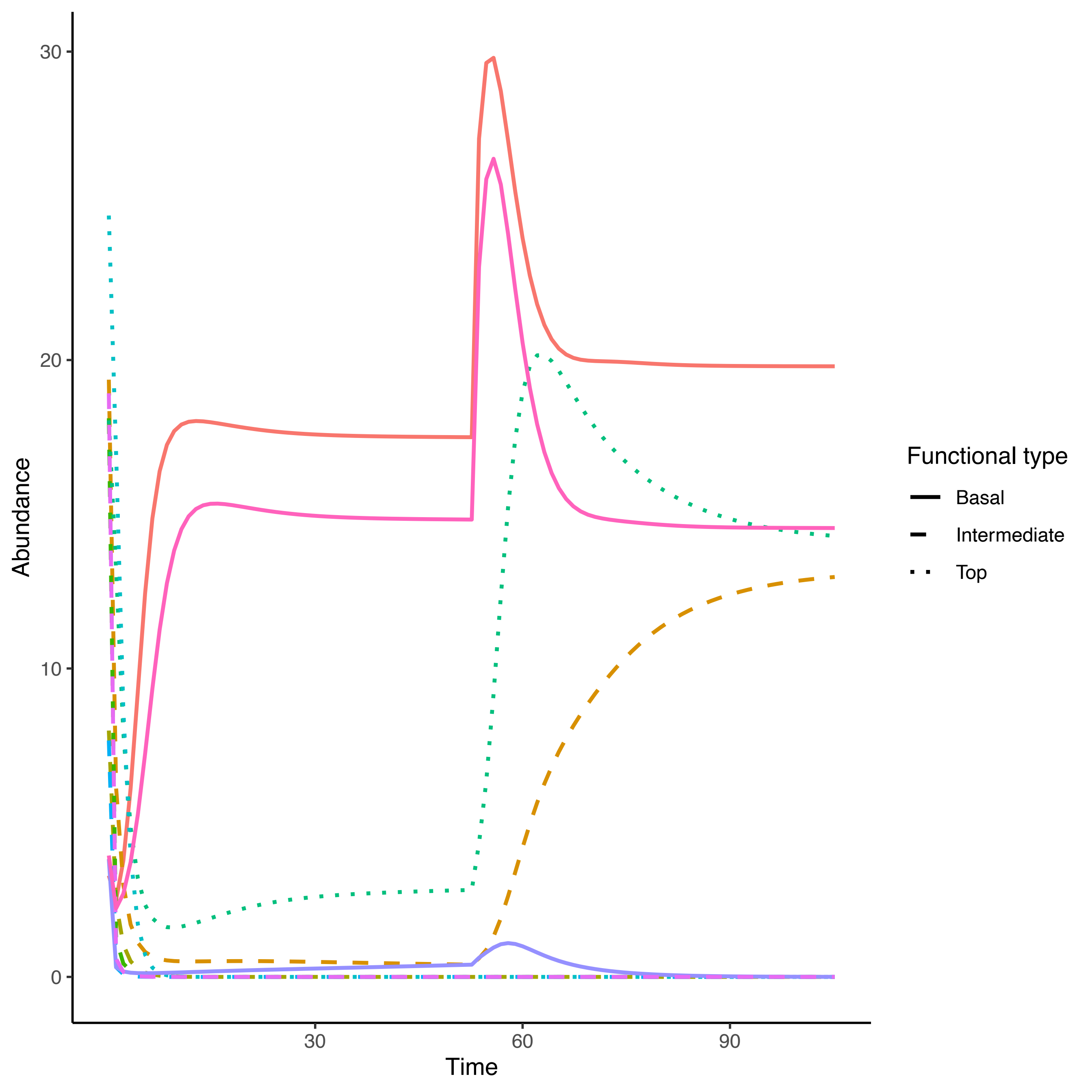

Abundance
30
20
10
0
30
60
90
Time
Functional type
Basal
Intermediate
Top

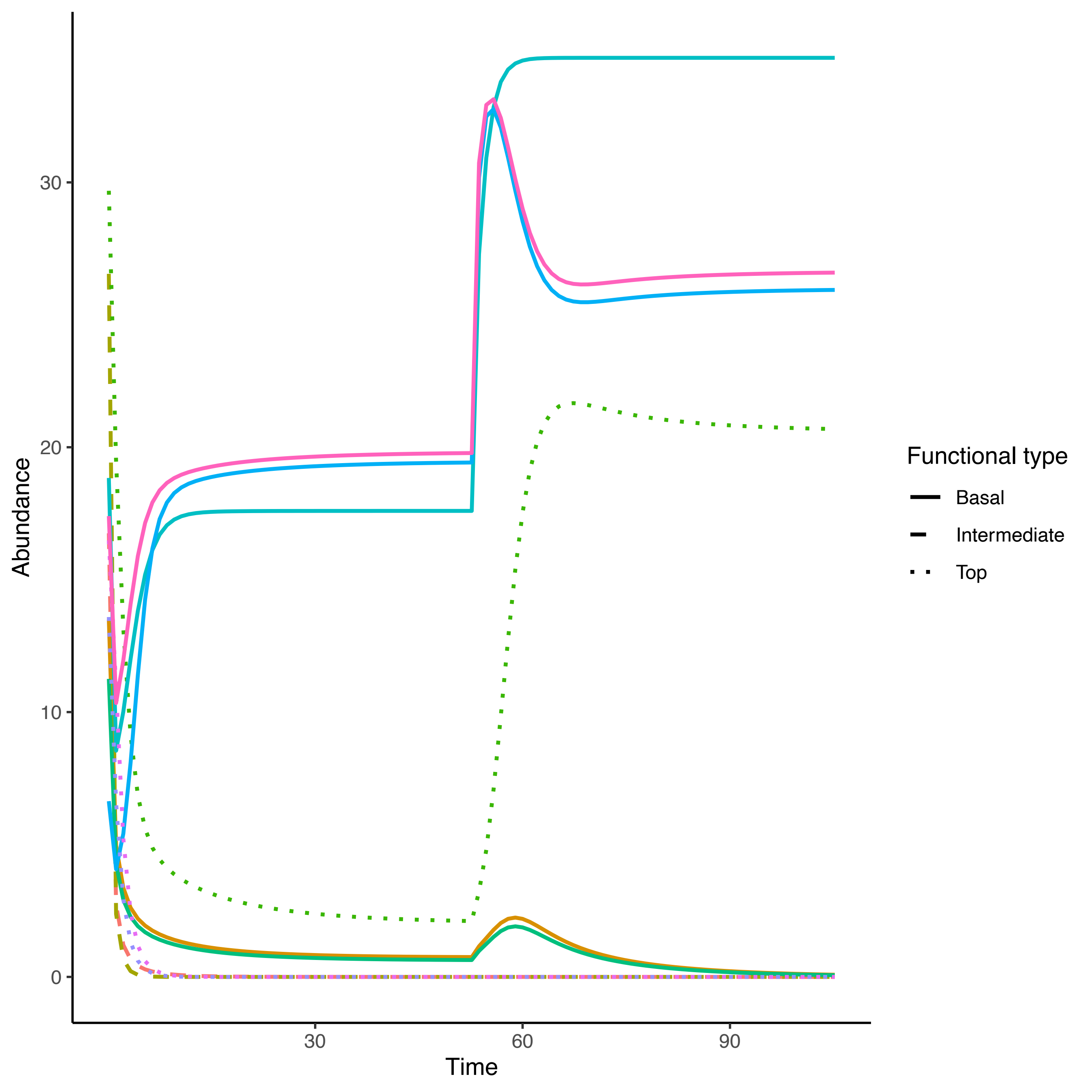
Abundance
Time
0
10
20
30
30
60
90
Functional type
Basal
Intermediate
Top

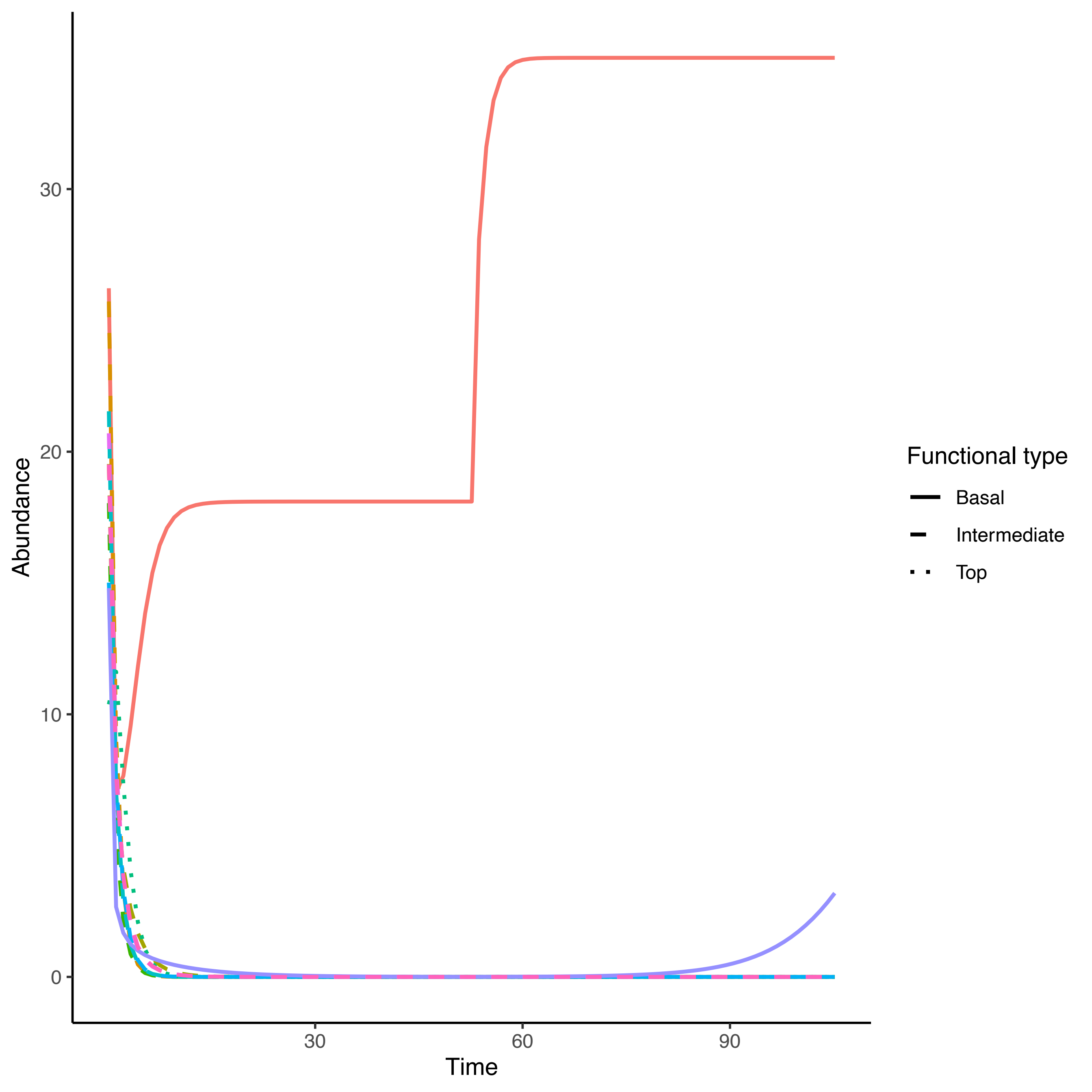

Abundance
Time
0
10
20
30
30
60
90
Functional type
Basal
Intermediate
Top

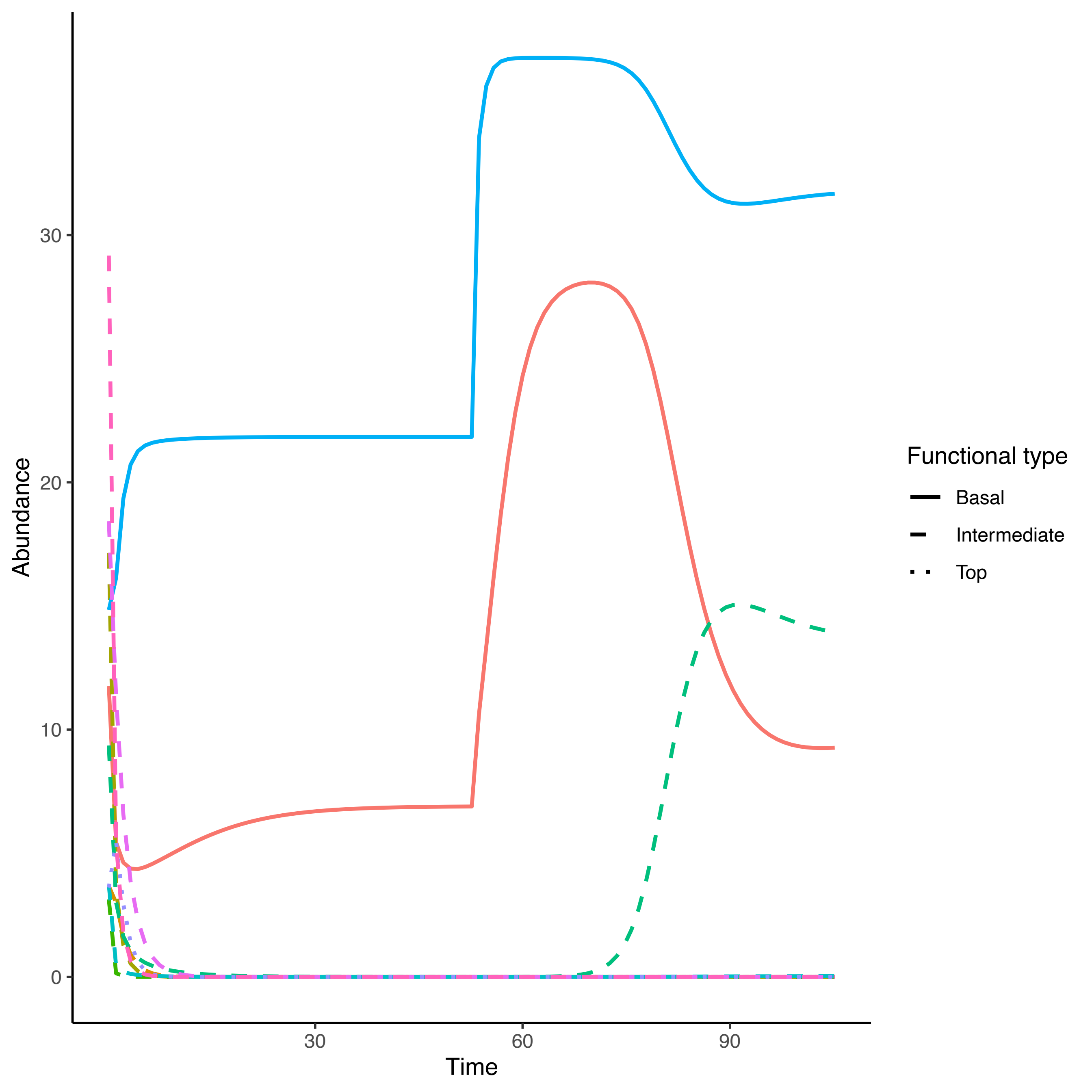

Abundance
0
10
20
30
Time
30
60
90
Functional type
Basal
Intermediate
Top

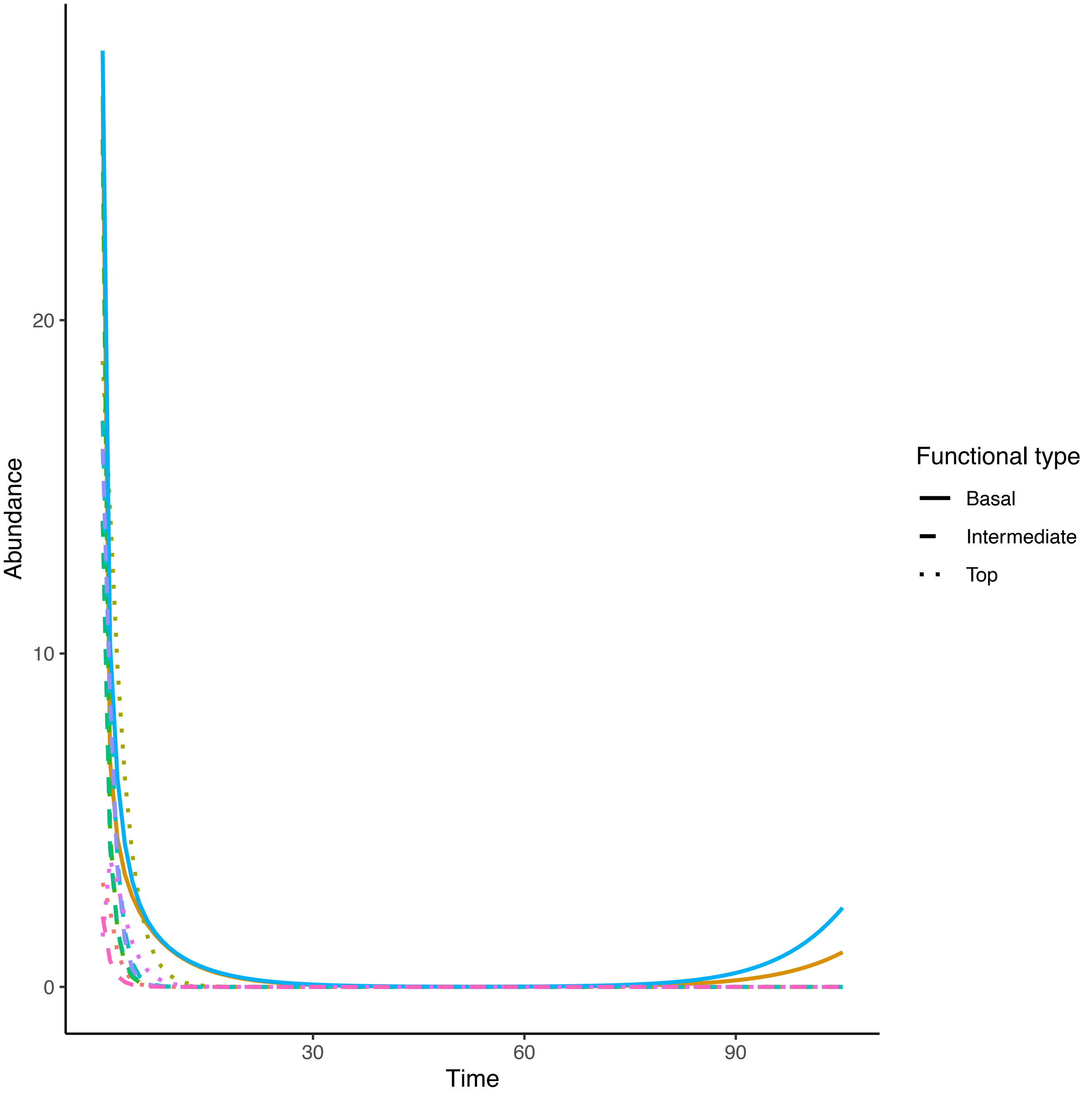

Abundance
20
10
0
Time
30
60
90
Functional type
Basal
Intermediate
Top

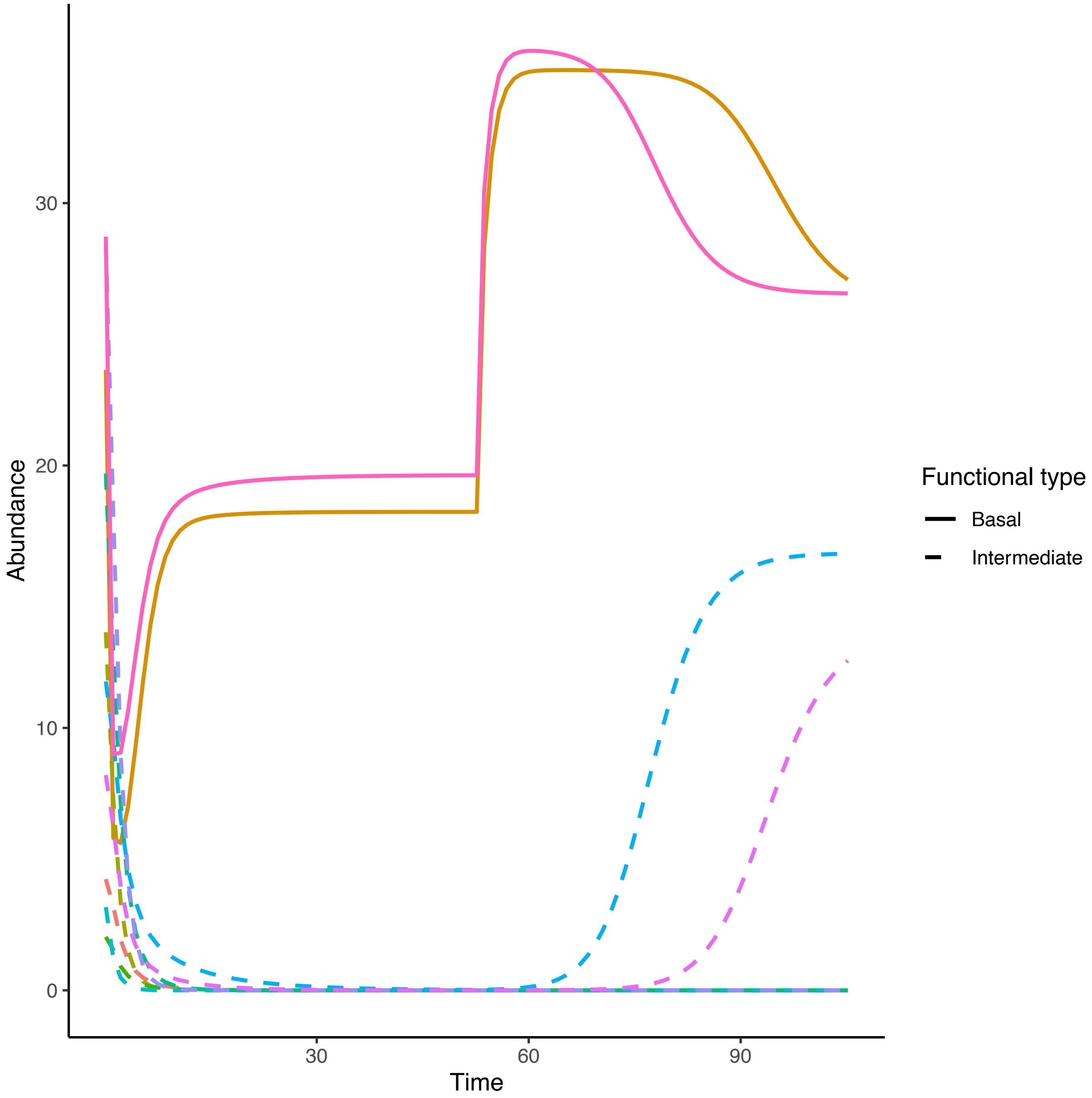

Abundance
30
20
10
0
30
60
90
Time
Functional type
Basal
Intermediate

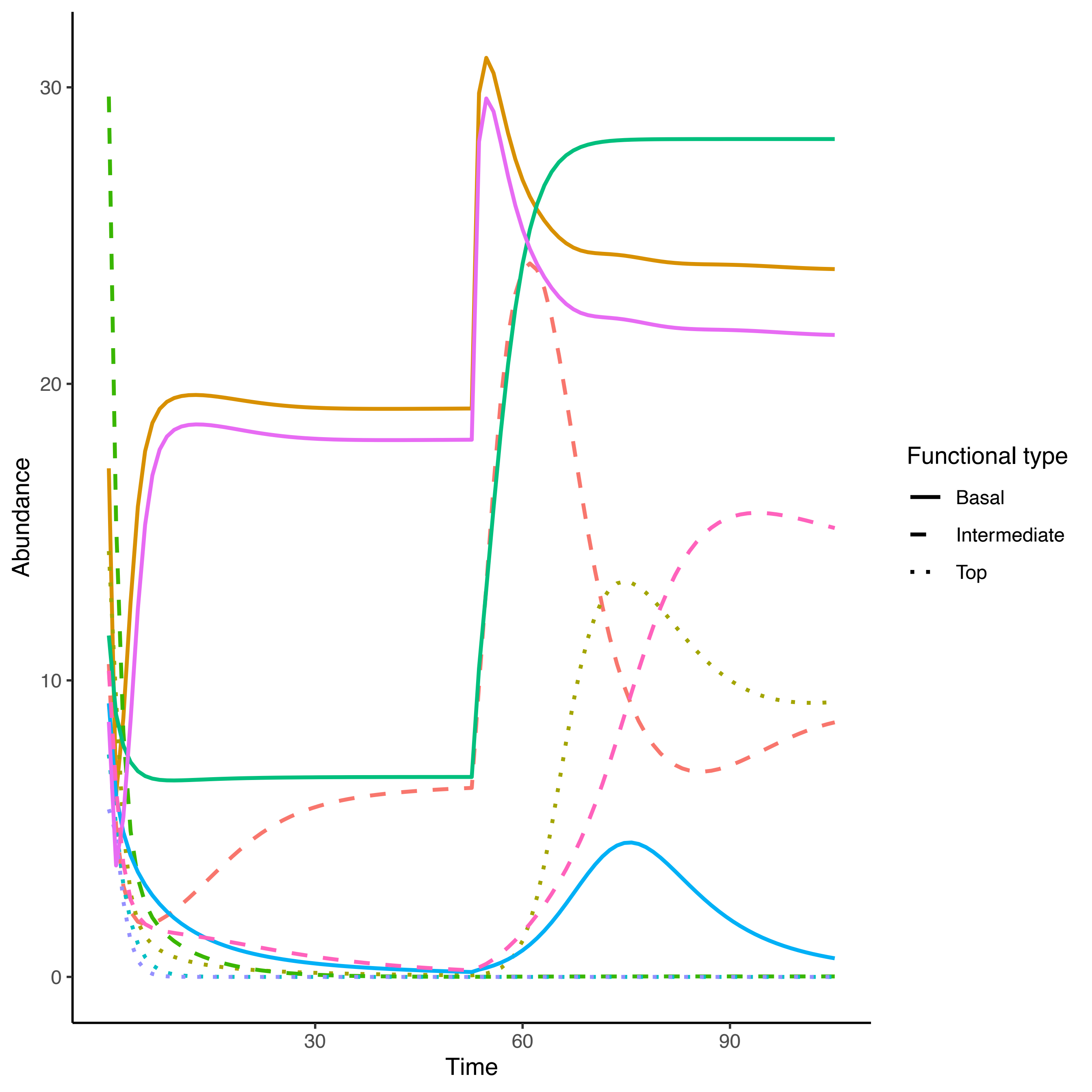
Abundance
0
10
20
30
Time
30
60
90
Functional type
Basal
Intermediate
Top
